# PriOmics: integration of high-throughput proteomic data with complementary omics layers using mixed graphical modeling with group priors

**DOI:** 10.1101/2023.11.10.566517

**Authors:** Robin Kosch, Katharina Limm, Annette M. Staiger, Nadine S. Kurz, Nicole Seifert, Bence Oláh, Stefan Solbrig, Marita Ziepert, Emil Chteinberg, Rainer Spang, Reiner Siebert, Helena U. Zacharias, German Ott, Peter J. Oefner, Michael Altenbuchinger

## Abstract

Mass spectrometry (MS)-based high-throughput proteomics data cover abundances of 1,000s of proteins and facilitate the study of co- and post-translational modifications (CTMs/PTMs) such as acetylation, ubiquitination, and phosphorylation. Yet, it remains an open question how to holistically explore such data and their relationship to complementary omics layers or phenotypical information. Network inference methods aim for a holistic analysis of data to reveal relationships between molecular variables and to resolve underlying regulatory mechanisms. Among those, graphical models have received increased attention as they can distinguish direct from indirect relationships, aside from their generalizability to diverse data types. We propose PriOmics as a graphical modeling approach to integrate proteomics data with complementary omics layers and pheno- and genotypical information. PriOmics models intensities of individual peptides and incorporates their protein affiliation as prior knowledge in order to resolve statistical relationships between proteins and CTMs/PTMs. We show in simulation studies that PriOmics improves the recovery of statistical associations compared to the state of the art and demonstrate that it can disentangle regulatory effects of protein modifications from those of respective protein abundances. These findings are substantiated in a dataset of Diffuse Large B-Cell Lymphomas (DLBCLs) where we integrate SWATH-MS-based proteomics data with transcriptomic and phenotypic information.

**GRAPHICAL ABSTRACT:** 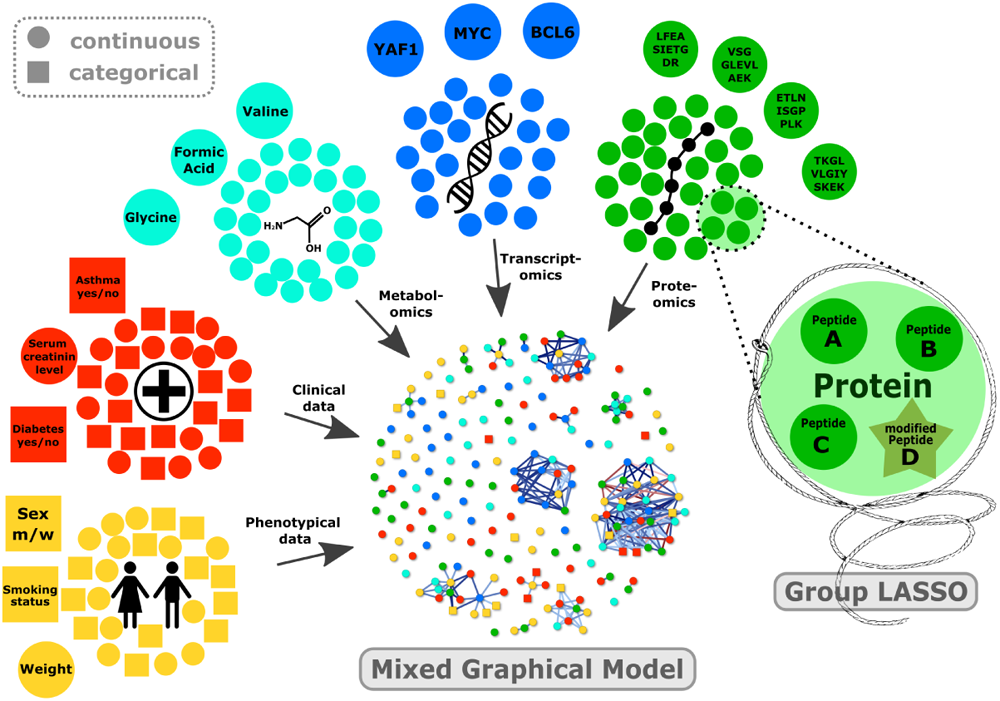

## INTRODUCTION

State-of-the-art proteomics approaches such as liquid chromatography coupled to quadrupole-time of flight tandem mass spectrometry (LC-QTOF-MS/MS) facilitate the high-throughput characterization of proteomes [1]. The field developed from assessing proteins, peptides and associated co- and post-translational modifications (CTMs/PTMs) in a few specimens, to large-scale studies in medicine and systems biology comprising hundreds of specimens to resolve regulatory mechanisms and to discover novel biomarkers [2, 3, 4, 5]. Studies of CTMs/PTMs have received increased attention in recent years, comprising among others phospho-proteomics to reveal kinase-substrate interactions in plants [6], the identification of high-risk groups in cancer [7], and the study of molecular mechanisms in cancer treatment [8].

Omics data integration refers to the study of biological processes using (multi-)omics data. This integration process requires statistical approaches capable of resolving the complex interdependencies among variables within and between different “tiers” of omics data. Naive pair-wise measures of association such as Pearson’s correlation were shown to be prone to false-positives as a consequence of indirect relationships mediated by one or more other variables [9, 10]. In order to resolve such spurious associations, molecular variables have to be considered in their multivariate context. Probabilistic Graphical Models are designed for this purpose [11, 10] and were recently adapted to the multi-omics setting [9] to study, for instance, gene regulation via promoter methylation. However, (multi-)omics datasets might demonstrate even higher complexity, which is not yet captured by existing approaches. First, (multi-)omics data are usually accompanied by complex and highly relevant phenotypical data that warrant direct incorporation into statistical model development. Second, some omics technologies require a tailored solution. For instance, the incorporation of genomic information, e.g., somatic mutations in cancer, should be incorporated as categorical variables. Similarly, state-of-the-art proteomics data require tailored solutions, as outlined in the following.

Bottom-up LC-MS/MS proteomic approaches like SWATH-MS (Sequential Window Acquisition of all THeoretical fragment-ion spectra mass spectrometry) quantify protein expression on the level of peptides following digestion of proteomes with endopeptidases such as trypsin. The measurement of thousands of peptides across hundreds of samples provides highly reproducible and consistent quantitative results [12]. Peptide intensities are typically computed by summing or averaging peak areas of the most intense fragment ions of a peptide. To infer the abundance of a protein, one or many proteotypic peptides are aggregated to yield protein intensities. Multiple strategies have been suggested for this process [13, 14, 15], of which the averaging of the most intense peptides per protein is one of the most common strategies [16]. These aggregation steps can potentially lower measurement noise, but they also have several major drawbacks. First, peptides contributing to the same protein intensity can differ with respect to measurement quality (noise) and, thus, the computed protein abundance can be blurred through low-quality peptide measurements. Thus, a specific peptide could be a better representative of the protein than the average value. Second, CTMs/PTMs, such as protein N-terminal acetylation, phosphorylation, glycosylation, and ubiquitination, can affect protein structure, function, and stability. For instance, the RNA and protein encoded by the gene *MYC* are known to be of high prognostic relevance in many cancer entities [17]. However, its phosphorylation patterns can confer additional prognostic value, as shown for medulloblastomas in [7]. In order to reveal the specific roles of protein MYC and its differentially phosphorylated isoforms, their respective quantities have to be considered. Naive aggregation can potentially hide such highly relevant information.

**Figure 1:**
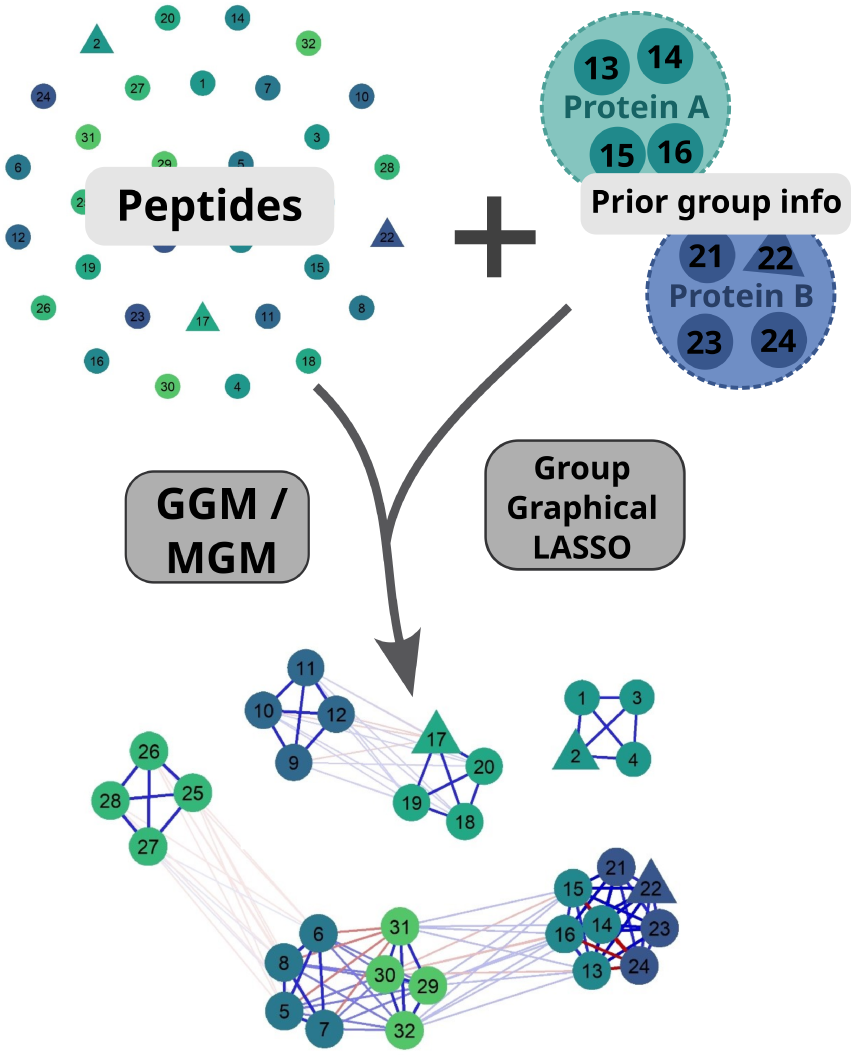
Concept of PriOmics. Peptides are grouped within a GGM or MGM by applying a Group Graphical LASSO, which penalizes the peptides according to their underlying proteins. Small circles indicate peptides, while peptides containing a co- or post-translational modification are indicated by triangles. Peptides with the same color contribute to the same protein.

**Figure 2:**
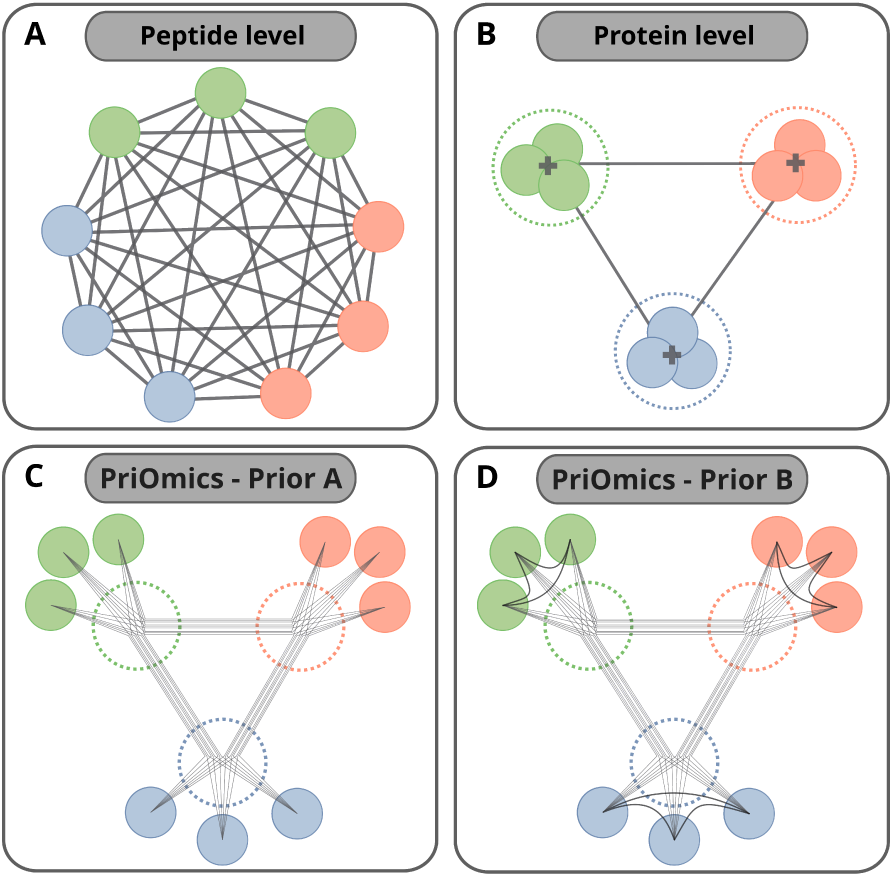
Visualization of different network inference concepts using high-throughput proteomics data. Figure (A) shows a naive network inferred on the level of peptides and Figure (B) on the level of proteins. PriOmics *Prior A* and *B* are shown in Figure (C) and (D), respectively. Both account for the protein affiliation but *Prior A* removes edges *a priori* between proteotypic peptides of a protein, while *Prior B* retains them. The former is more appropriate for technical, noisy copies of an identical variable (peptides which represent the same protein abundance) and the latter is more appropriate for variables which are related but likely represent different biological variables (for instance, different phosphorylation sites of the same protein).

Here, we introduce PriOmics, which is a probabilistic graphical modelling approach to account for redundant molecular variables (multiple peptides of the same protein) without a prior aggregation step. PriOmics models individual peptides but takes into account their corresponding protein identity. This allows PriOmics to disentangle regulatory effects of protein modifications from those of respective protein abundances. PriOmics can also incorporate categorical data types to facilitate the integration of complex proteomics data with pheno- and genotypical information. We demonstrate the performance of PriOmics both in simulation studies and in an application to proteome-transcriptome data integration in Diffuse Large B-Cell Lymphoma (DLBCL), where we also take into account phenotypical information on patient samples. The latter analysis reveals key molecular variables that drive the differentiation of DLBCLs into their cell of origin.

## MATERIALS AND METHODS

### Mixed graphical model

#### Probability density function

Mixed Graphical Models (MGMs) can simultaneously account for different data types. We will build on the probability density function

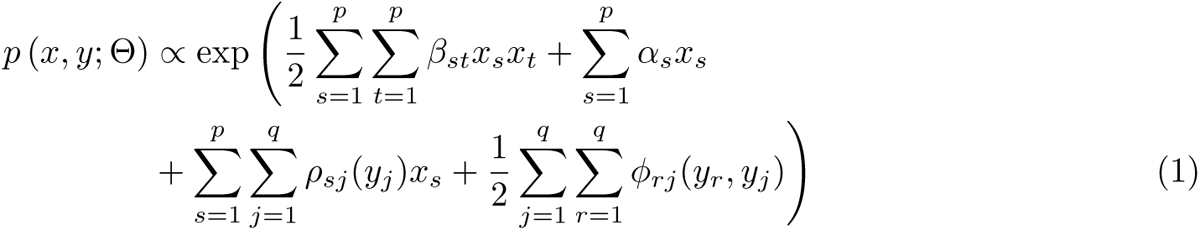

proposed by [18], which incorporates *p* standardized continuous variables *x_s_*, where *s* = 1*, . . . , p*, and *q* categorical variables *y_j_* with *L_j_* states, where *j* = 1*, . . . , q*. The model parameters are 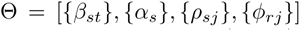. The term 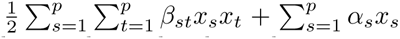 corresponds to a Gaussian Graphical Model (GGM), which encodes conditional independencies among multivariate gaussian distributed variables, and the term 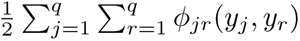 corresponds to a Markov Random Field 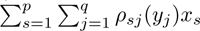 encodes the conditional dependencies between discrete variables *y_j_* and continuous variables *x_s_*. Following [18], Θ can be estimated using the negative pseudo-log-likelihood

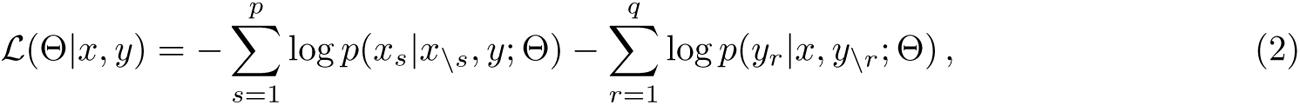

with the conditional probabilities 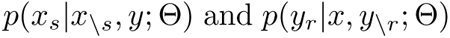, where *x\_s_* is the set of all continuous variables excluding *s*, and *y\_r_* the set of all categorical variables excluding *r*. Eq. (2) is a high-dimensional optimization problem that can be prone to overfitting if *p* and *q* are larger or of the same order of magnitude as the sample size *n*. For an appropriate regularization strategy see, e.g., [18].

### PriOmics

PriOmics builds on mixed graphical modelling to facilitate the integration of high-throughput proteomics data with additional complex data sources, such as complementary omics layers and/or phenotypical information. It models the data on the level of peptides to reveal potentially relevant regulatory mechanisms, and induces parsimonious network structures taking into account pre-specified peptide groups (see also Figure 1). Importantly, PriOmics allows to quantify our assumptions about biological data by selecting between two different grouping priors. A summary of these priors together with standard peptide-based and protein-based network inference concepts is shown in Figure 2.

#### Group priors

##### *Prior A* – grouped conditionally independent variables

Multiple proteotypic peptides likely represent the same biological variable in a network (here a protein abundance). However, this cannot be guaranteed. For instance, summing up or averaging only the intensities of unmodified peptides or both unmodified and modified peptides of a protein may mask important functional information. Thus, there are good reasons for either treating peptides as technical copies or as individual biological variables. Consequently, an appropriate prior should be a trade-off between both. Our suggested strategy is to enforce conditional independence among peptides, which belong to the same protein, and to group them via group regularization terms, as argued in the following.

The conditional probability 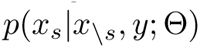 corresponds to the linear regression of response variable *x_s_* on the predictor variables *x\_s_* and *y* [19]. Let variable *x_s_* be member of a group *S* where *S* is a set of redundant variables (technical copies), *i.e.*, noisy copies of an identical biological variable. Then, first, if *x_s_* is the regression target, none of the variables 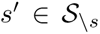 should be included as predictor variable; otherwise a perfect copy *x_t_* of *x_s_* would explain all variance of *x_s_*. This motivates the enforcement of conditional independence between *s* and *t* via setting *β_st_* = *β_ts_* = 0 during model development. Vice versa, if we consider variable *x_t_* with 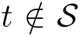 as regression target, then the variable which is represented by the members of group *S* contributes *| S|* times (via its copies). This constraint cannot be directly incorporated into model development, because it would violate Eq. (2). However, it is still possible to take into account the grouping property via appropriate regularization strategies. Namely, we employ a group-lasso regularization, which enforces group-wise sparseness parsimonious networks (see also Figures 1 and 2). To achieve this, we augmented Eq. (2) by regularization terms:

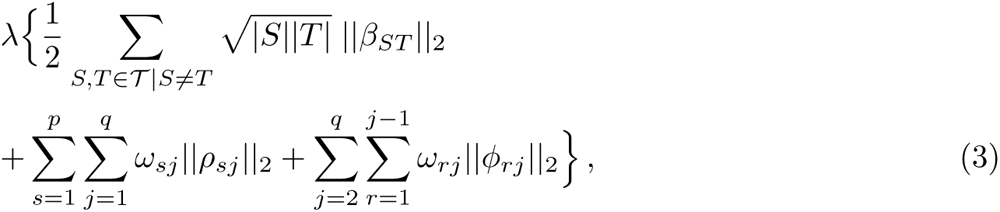

where *λ* is a regularization parameter and *|S|* gives the size of group *S*. Note that the weights *ω_ij_* of the individual terms in (3) are chosen such that only a single global hyperparameter *λ* has to be calibrated for model selection (compare [18]), and that all continuous variables are assumed to be standardized.

##### *Prior B* – grouped variables

This prior groups variables that are not ordinary copies of the same biological variable and, therefore, could potentially affect each other. Hence, we do not make any *a priori* assumptions about conditional independence among the grouped variables. A typical scenario where this prior will apply are differentially spliced mRNA isoforms of the same gene. Formally, *Prior B* accounts for the group membership only via regularization Eq. (3), and, thus, members of a group are allowed to be connected by an edge.

### Hyperparameter tuning

The calibration of hyperparameter *λ* was performed using either the Bayesian Information Criterion (BIC) [20] or the extended Bayesian Information Criterion (EBIC) [21], where we evaluated the pseudo-log-likelihood of Eq. (1) as suggested by [22] to reduce computational burden. Throughout this article, we used either BIC or EBIC with hyper-parameter *γ* = 0.5. The latter results in a moderately more conservative model. For each model we evaluated a series of 30 different *λ* values.

### Competing methods

Methods for the inference of mixed graphical models are still scarce. We evaluated the following competing methods:

#### Pseudo-log-likelihood (PLL-MGM)

We implemented the pseudo-log-likelihood based method suggested in [18], which forms also the backbone of PriOmics. Therefore, this implementation also serves as baseline to assess the performance of the suggested PriOmics group priors.

#### Node-wise lasso regression for mixed graphical models (NW-MGM)

We used the implementation of the node-wise MGM approach provided in the R-package ‘mgm’ [23].

### Simulations

We generated artificial data to evaluate the performance of PriOmics compared to PLL-MGM and NW-MGM. All methods were assessed on their capability to recover *a priori* defined edges, *i.e.*, non-zero model parameters. Data were simulated using Gibbs sampling, where ground truth parameter sets were simulated as follows: (0) initialize all parameters as zero, (1) sample off-diagonal elements of *β*, and off-diagonal blocks of *ρ* and *ϕ*, with the latter corresponding to individual continuous-discrete and discrete-discrete couplings, respectively, for given edge densities (Table 1), (2) replace sampled elements by values drawn from a uniform distribution *U* (*−*1, 1), (3) fill the diagonal of the joint precision matrix with the row-wise sum over all its absolute values and normalize it accordingly, and (4) replace the diagonal of 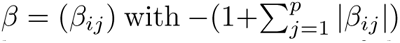 (for details see supplementary code). Redundant variables were simulated by generating noisy copies of the underlying protein variables with noise given by *ɛ ∼ N* (*µ* = 0*, σ* = 0.5).

### Performance measures

The performance of the algorithms was evaluated by comparing the estimated model parameters to the ground-truth Θ used to simulate *X* and *Y* . There, we analyzed if an association between two features (*i.e.*, the presence of an edge) could be reconstructed by considering the absolute values of edge strengths, and by calculating respective areas under the curve of receiver operating characteristics (AUC-ROC) and the AUC of precision-recall curves (AUC-PR).

## RESULTS

We will first illustrate the basic concept of PriOmics and will show that holistic analysis strategies of high-throughput proteomics data have to account for the fact that multiple proteotypic peptides likely represent the same underlying protein abundance. Otherwise, analyses are prone to erroneous conclusions. These findings will be substantiated in systematic simulation studies presented in subsequent sections. Finally, we will provide an exemplary application to a combined proteomics-transcriptomics dataset of DLBCLs.

### Graphical models, redundant variables and group-parsimonious networks

The inference of probabilistic graphical models is affected by redundant variables. To illustrate this, we generated artificial data according to the exemplary network structure shown in Figure 3A, consisting of 5 continuous variables (A, B, C, D, E) and 3 categorical variables with binary outcome (X, Y, Z). Then, we generated two noisy copies of B and three noisy copies of variables C and E, to simulate the possible presence of multiple peptides representing the same protein. The corresponding ground truth network is visualized in Figure 3B, where we contrast it with different approaches to reconstruct the underlying network (Figs. 3C to G). These are, first, the network structure estimated via ordinary univariate analysis, where we used linear regression for continuous-continuous edges, and logistic regression for continuous-binary and binary-binary edges (Fig. 3C). Since each edge is estimated twice, we show the respective average (only edges with BH-adjusted *p*-values *<* 0.05 are shown, Matthews correlation coefficient (MCC) =0.71). We observed a pattern typically seen in ordinary pair-wise association analysis, as exemplified by variables B and D, which are individually connected to variable E, but not to each other in the ground truth network Figure 3B. Erroneously, the univariate analysis in Figure 3C shows an edge between B and D as a consequence of B and D’s individual relationship to E. This observation illustrates that pair-wise association measures are prone to false positive associations and, thus, molecular variables have to be considered simultaneously in order to disentangle direct from indirect relationships. Similar observations were repeatedly made for pair-wise correlation networks [9]. Note that the latter are restricted to continuous variables only. As an alternative analysis strategy, we explored multivariate node-wise linear and logistic regressions (Fig. 3D). This analysis strategy takes into account that there might be indirect relationships, but it does not allow for variables to be technical replicates of each other. In fact, all genuine interactions were too weak and, consequently, were removed after BH-adjustment (MCC=0). This effect was substantially mitigated using the classical, regularized MGM as proposed by [18] (Figure 3E). However, still a high proportion of edges were not observed (MCC=0.72). In particular, when replicated variables were involved, an edge was usually only observed to individual members and not to all replicates, as exemplified for edges between variables representing B & E and A & C, respectively. Next, we evaluated PriOmics and obtained the networks shown in Figure 3F and G for *Prior A* and *B*, respectively. PriOmics *Prior A* correctly distinguished direct from indirect relationships, as can be seen from the absence of edges between B and E and between A and C, respectively. Moreover, it can deal with technical replicates of variables; if there is an edge between one peptide and another variable, then all peptides of the respective protein are connected to this variable. PriOmics - *Prior B* shows a similar behavior (Figure 3G). However, since *Prior B* does not strictly assume that variables of the same group are technical replicates, it infers also relationships between those replicates, at the price of reduced edge weights to other variables. In summary, PriOmics can deal with redundant variables, but requires *a priori* specification of the underlying data generating process. The latter is not always available and, as a consequence, performance can be compromised, as can be seen by comparing 3F and G (MCC=1. vs MCC=0.93); the prior was correctly specified for 3F (*Prior A*, as variables B, C, and E contain technical replicates) and incorrectly for 3G.

**Figure 3:**
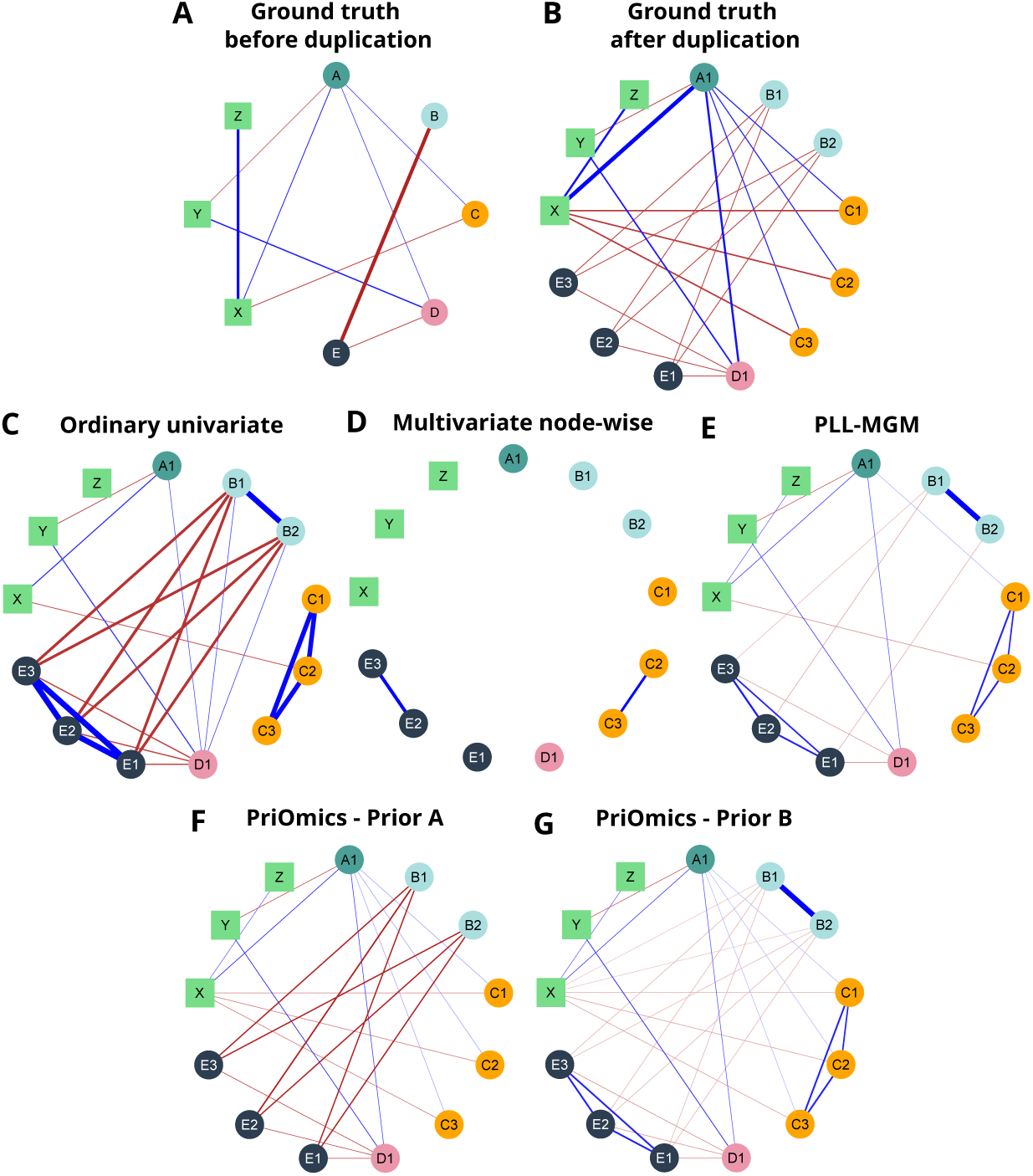
Illustration of network inference approaches on a dataset simulated according to the procedure described in Suppl. Figure S1. Circles correspond to continuous features and squares to categorical features. Equally colored continuous features represent noisy copies of the same feature. Blue colored edges indicate positive associations, while red edges indicate negative associations. The edge width and color depth indicate the association strength.

An important ingredient of this performance gain is the PriOmics regularization scheme. PriOmics adapts the MGM loss function to account for redundant variables by enforcing within-group edges to equal zero (*Prior A*) and makes use of regularization strategies taking into account the grouping of variables (*Prior A* and *B*).

The need for model regularization has been emphasized repeatedly for GGMs [24, 25] and MGMs [18]; for *p > n* the maximum likelihood estimate of the multivariate Gaussian is not defined and even if *n > p*, the parameter estimates can have a large variance if both are of the same order of magnitude. Thus, model regularization is mandatory. Standard lasso regularization enforces sparseness with respect to individual edges (see, *e.g.*, the graphical lasso [26]), while PriOmics utilizes group-lasso regularization terms to further encode the variable groups. To illustrate this feature, we repeated the previous analysis but now decreased the regularization parameter successively from *λ* = 1.6 to *λ* = 0. Each individual *λ* was applied on both regularization strategies, *Prior A* (Suppl. Figure S3) and *B* (Suppl. Figure S4). We make the following observations:

1. Networks become increasingly connected upon decreasing *λ*.
2. Variables that belong to the same variable group are either all connected to a neighbor or not connected.

The latter feature is a consequence of the group-lasso terms and has important consequences not just for model performance but also for model interpretability; edges are always drawn to all members of a group and, thus, respective edge weights can be directly compared. The latter feature is lost when regularization enforces sparseness in individual edges, as seen in Suppl. Figure 5, where PLL-MGM was evaluated for different values of the regularization parameter.

### Simulation studies

The former example illustrated the basic features of PriOmics. To further demonstrate the performance gain of PriOmics compared to other methods, we performed several simulation studies, exploring both different sample sizes and different edge densities (the proportion of edges).

#### PriOmics improves on network inference in settings with redundant variables

We evaluated PriOmics and the competing methods for a fixed number of 200 continuous and 30 discrete variables for different sample sizes *n* (*n ∈ {*100, 175, 250, 375, 500, 750, 1000, 1500, 2000*}*), where the subsets were always drawn from the next bigger one. The continuous variables were included as noisy copies to account for a scenario similar to that of different peptides representing the same protein (see supplementary code). We performed three different simulation scenarios A, B, and C with varying edge densities *η* for continuous-continuous, continuous-discrete, and discrete-discrete edges as summarized in Table 1. Each simulation scenario was repeated 10 times to provide error estimates.

**Table 1:**
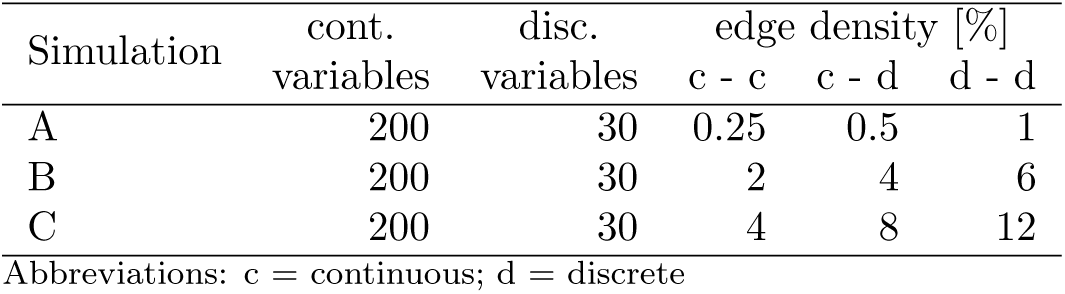
Description of simulated datasets.

| Simulation | cont. | disc. | edge density [%] |  |  |
| --- | --- | --- | --- | --- | --- |
|  | variables | variables | c - c | c - d | d - d |
| A | 200 | 30 | 0.25 | 0.5 | 1 |
| B | 200 | 30 | 2 | 4 | 6 |
| C | 200 | 30 | 4 | 8 | 12 |

Abbreviations: c = continuous; d = discrete

First, we compared the performance of PriOmics to PLL-MGM and NW-MGM with respect to the reconstruction of continuous-continuous edges, with results shown in Figure 4A to C. The AUC-ROCs for *Prior A* were consistently higher than those obtained for the MGMs without peptide grouping, with improvements of up to 34.8% for simulation A, 31.1% for simulation B, and 27.1% for simulation C. Similarly, *Prior B* performed better than PLL-MGM and NW-MGM for all sample sizes, but improvements were smaller than for *Prior A*, see Figure 4. In general, AUCs increased with *n*, and interestingly, the largest improvements were observed for the large sample sizes, while improvements for samples sizes *n <* 250 were smaller but still better for both PriOmics *Prior A* and *B*. In general, the edge recovery of estimated models was better for small edge densities *η* with maximum AUCs of 0.962, 0.925, and 0.749 for simulation scenarios A, B, and C, respectively. Next, we verified the reconstruction of continuous – discrete edges (Fig. 4, middle Figure D to F) and discrete – discrete edges (Fig. 4, bottom Figure G to I). For all methods, AUCs had a tendency to be lower than for the reconstruction of continuous – continuous edges, with PriOmics clearly out-competing NW-MGM and showing similar performance to PLL-MGM for both *Prior A* and *B*. One should note that the model priors directly affect continuous-continuous edges via prior Eq. (3), while their effect on continuous – discrete and discrete – discrete is only an indirect one, which might explain the minor differences to PLL-MGM. Corresponding results considering area under the precision-recall curves (AUC-PR) and MCCs are given in Suppl. Figure S6. In summary, this simulation study shows that both PriOmics *Prior A* and *B* substantially improve edge recovery compared to the state-of-the-art approaches PLL-MGM and NW-MGM through systematically addressing redundant variables by removing within-group edges (*Prior A*) and by a regularization scheme, which *a priori* takes into account the grouping of variables (*Prior B*).

### Application to DLBCL data

Non-Hodgkin lymphomas (NHL) arising from B- or T-cells represent a highly heterogeneous group of diseases with diffuse large B-cell lymphomas (DLBCL) being the most frequent, accounting for almost one third of NHL [27]. Routinely, DLBCL patients are stratified simply according to the stage of the disease (i.e., the sites of manifestations), or taking addtional phenotypic and clinical variables into account, such as age, general constitution (*i.e.*, performance status), serum lactate dehydrogenase (LDH) level or the occurrence of B symptoms (*i.e.*, fever, drenching night sweats, and loss of more than 10 percent of body weight over 6 months), according to the International Prognostic Index (IPI) [28]. On the molecular level, DLBCL are commonly differentiated according to their cell of origin (COO) into germinal center B-cell like (GCB) and activated B-cell like (ABC) depending on the expression of specific genes [29, 30]. More recently, aggressive lymphomas expressing the ”double hit” or the ”molecular high grade” signatures have been proposed as a separate subtype [31, 32]. These signatures characterize GCB tumors with genetic and dark zone biological features similar to both follicular lymphomas and Burkitt lymphomas. In addition, molecular classification schemes based on transcriptomics and proteomics were proposed in [33, 34, 35, 36] and, more recently, more sophisticated genetic classifiers [37, 38]. These recent developments warrant a holistic approach for revealing different pathomechanisms underlying DLBCL and predicting treatment response. Here, we will perform an exemplary analysis of a high-throughput proteomics dataset of DLBCL using PriOmics.

#### Description of the dataset and preprocessing

##### Dataset

The dataset includes 344 DLBCL specimens from the German High-Grade Lymphoma Study Group (DSHNHL) [36], comprising proteomics data of formalin-fixed paraffin-embedded (FFPE) tissue slides generated using microLC-SWATH-MS, including in total the label-free mass spectrometric intensities of 7,720 unmodified peptides and 666 peptides with co- and/or post-translational modifications. Two additional transcriptomics datasets were available for 330 of the 344 patients, consisting of 145 genes measured by NanoString nCounter [35] and 296 genes measured by HTG EdgeSeq (see overview in Table 2), both containing typical genes relevant for COO-subtyping and genes characterizing the tumor-microenvironment. The ”HTG EdgeSeq Pan B-Cell Lymphoma Panel” was used following the instructions from HTG Molecular Diagnostics. Briefly, FFPE samples underwent among others heat treatment and proteinase K incubation in the HTG EdgeSeq system. The RNA protected DNA probes were purified with AMPure XP beads and quantified using the KAPA Library Quantification Kit. Library concentrations were determined using the HTG RUO library calculator. Libraries were sequenced using an Illumina NextSeq 550 device. Raw counts of the probes were extracted from FASTQ using HTG EdgeSeq analyzer software. In addition to the omics data, 12 categorical variables (nine binary variables & three multi-label variables) were gathered (Table 3). First, we applied PriOmics to the proteomics dataset including the clinical and phenotypical features. We refer to this analysis as Dataset 1 (DS1). Second, we applied PriOmics to the joint dataset (DS2), consisting of proteomics, transcriptomics, and clinical and phenotypical data. Datasets DS1 and DS2 are summarized in Table 2.

**Table 2:** Number and type of features included in the DLBCL dataset.

|  | Peptides |  | Genes | Pheno/Clin | Patients |
| --- | --- | --- | --- | --- | --- |
|  | no CTM/PTM | CTM/PTM |  |  |  |
| DS1 | 2,033 | 198 | 0 | 12 | 344 |
| DS2 | 2,033 | 198 | 145+296 | 12 | 330 |

Abbreviations: DS = Dataset; CTM = co-translational modification; PTM = post-translational modification; Pheno/Clin = phenotypical or clinical

##### Preprocessing

The proteomics data contained a total of 80 missing values (0.0000023 % of all measurements), which were imputed with the respective patient sample mean value. To improve data quality, we only included those peptides, which showed a local FDR value [39] lower than 0.2 in at least 80% of patients (0.4 in at least 60% of patients for the modified peptides). The final datasets included 2,033 unmodified and 198 modified peptides, which could be attributed to 1,056 individual proteins. One of the modified peptides was found twice that differed only in charge state. The most common modification observed in 120 out of 197 (60.9%) modified peptides was co-translational protein N-terminal acetylation with (*n*=87) or without (*n*=33) prior cleavage of the N-terminal methionine by methionine aminopeptidases [40]. These peptides can be easily detected without prior enrichment, as N-terminal acetylation of the alpha-amino group of peptides improves the occurrence and abundance of so-called b and y ions in collision-induced dissociation, thus facilitating their detection by SWATH-MS [41]. Since protein N-terminal acetylation affects all copies of a protein, abundance of the acetylated N-terminal peptide is expected to correlate well with the abundance of any other unmodified peptide of the same protein. Therefore, they are unlikely to provide any additional biological insight. This may not be the case for PTMs. Genuine PTMs observed in the present study included the post-translational phosphorylation of 8 threonine and 4 serine residues, respectively, of which 8 had been reported previously, as well as the acetylation and trimethylation of the epsilon-amino group of lysine residues. In contrast, other modifications observed, such as formylation of the epsilon-amino group of lysine residues of proteins other than histones, N-terminal carbamylation, deamidation of asparagine and glutamine, and oxidation of methionine are typical artefacts occurring during formalin fixation of tissues, urea-based protein extraction, and trypsin digestion at neutral or alkaline pH. Thus, they will not provide any meaningful biological insight. Both, the proteomics and transcriptomics data were normalized by (1) dividing each value by the corresponding patient mean, (2) performing a log_2_ transformation, and (3) by standardizing the variables.

**Figure 4:**
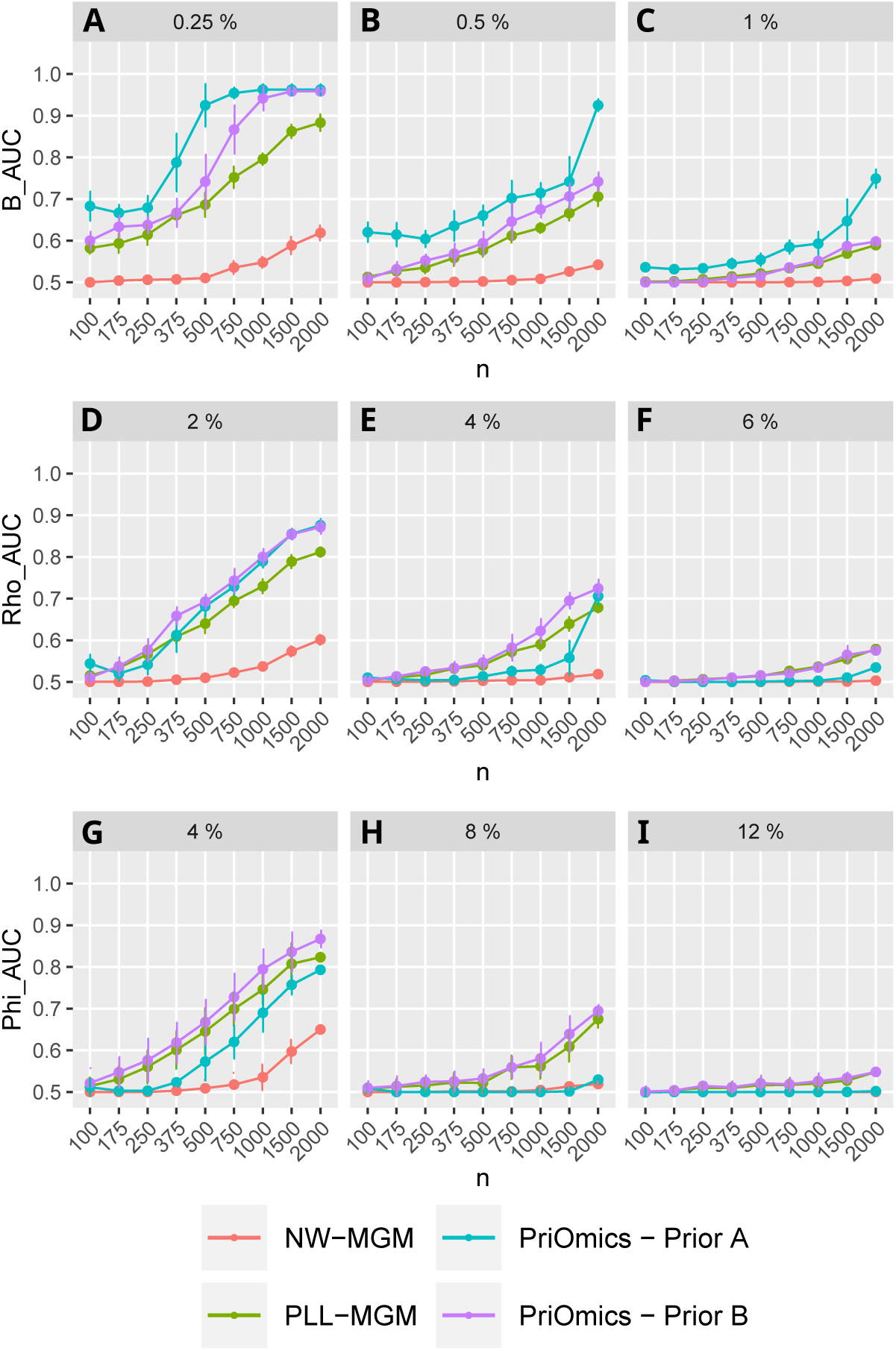
Performance comparison of a node-wise lasso regression MGM implementation (’NW-MGM’), a pseudo-log-likelihood MGM implementation (’PLL-MGM’), and the PriOmics implementations with different priors (’*Prior A*’ and ’*Prior B* ’) on simulated data. The plots show the average AUC versus sample size *n*. The upper row shows results for the continuous-continuous edges, the middle row for the continuous-discrete edges, and the last row for the discrete-discrete edges. The columns correspond to three different simulation studies with edge densities *η* given in the figure headings, as summarized in table 1. The error bars correspond to *±*1 standard deviation.

**Table 3:** Clinical and phenotypic features of the DLBCL dataset.

| Variable name | Description |
| --- | --- |
| AGE | Patient age > 60 |
| LDH | Serum lactate dehydrogenase elevated |
| ECOG | Performance status 3 or 4 (chronically bedridden) |
| STAGE | Disease stage III or IV according to the Ann Arbor staging system |
| EXBM | Two or more extranodal sites of disease (outside of a lymph node) |
| BULK | Presence of bulky disease (tumor larger than 10 cm) |
| BSYMP | Presence of B symptoms (e.g., fevers, night sweats and unexplained weight loss) |
| BM | Bone marrow (BM) involvement |
| GENDER | male or female |
| COO | DLBCL COO subtype classification:<br>GCB, ABC or unclassified |
| DHL | Double-Hit lymphoma status:<br>0 (no), 1 (yes) or unlabelled |
| DEL | Double-Hit expression status:<br>0 (no), 1 (yes) or unlabelled |

DHL is defined by the presence of MYC and BCL2 or BCL6 rearrangements, while DEL is defined by the expression of MYC and BCL2. Abbreviations: DLBCL = Diffuse Large B Cell Lymphoma; COO = Cell of origin; GCB = germinal center B cell-like; ABC = activated B cell-like

##### PriOmics model development

PriOmics was applied to DS1 in three different scenarios; using *Prior A* and *B* to encode the protein affiliation and without prior assumptions (i.e., PLL-MGM), respectively. The more comprehensive dataset DS2 was processed using the priors to group peptides according to their associated protein. The transcriptomics data were modelled as individual variables without underlying group property. All PriOmics models converged using a stopping criterion of *ɛ <* 10*^−^*^6^, defined according to [42]. In total, models for 30 different penalization parameters (*λ*) were evaluated. In a first instance, we selected models using the lowest EBIC with *γ* = 0.5 resulting in *λ* values of 1.130, 0.783, and 0.425, respectively, for models with *Prior A*, *Prior B*, and no prior assumptions. Since the selected models yielded only few non-zero associations as shown in Suppl. Table S1, we subsequently chose the BIC as model evaluation criterion. This yielded denser networks and facilitated a more detailed biological interpretation. Models with *λ* values of 0.332, 0.294, and 0.141, respectively, for *Prior A*, *Prior B*, and no priors were found to be optimal for regularization. A summary of the inferred models can be found in Suppl. Table S1 and Suppl. Figures S22-24. Analogously, the model selection for DS2 was conducted with the BIC using *λ* values of 0.294, 0.294, and 0.180, respectively, for *Prior A*, *Prior B*, and no prior. Further details of model selection are depicted in Suppl. Figures S25-27.

#### Holistic omics data integration with phenotypical variables in DLBCL

We used PriOmics to integrate proteomics and transcriptomics data with phenotypical variables summarized in Table 3 to obtain an overall picture of the variables’ interrelationships in DLBCL. This analysis provides both, sanity checks of the algorithm as well as results that would have remained hidden using routine analysis strategies. Unless otherwise mentioned, the analyses were performed with BIC as model selection criterion.

##### PriOmics reduces false positive findings

We first applied PriOmics and competing methods to the combined proteomics and phenotypical data (DS1). PriOmics yielded networks with 46,987 and 44,354 edges, corresponding to edge densities of 1.87% and 1.69% for *Prior A* and *Prior B*, respectively, without taking into account edges within protein groups to maintain comparability. Edges obtained by both approaches were consistent with an overlap of 63.3% (Figure 5B). PLL-MGM yielded a similar edge density of 2.05%, but the individual edges agreed only moderately to those of PriOmics with an overlap of 10.3% and 11.4%, respectively, for *Prior A* (Figure 5C) and *B* (Figure 5D). One should note that if a peptide is connected to a variable, then presumably the remaining peptides of the same protein should be also connected to this variable. This is always the case for PriOmics, as this prior knowledge is encoded via regularization. However, PLL-MGM yielded highly inconsistent results, where, on average, only 56.80% of the expected edges were selected. It is noteworthy that PLL-MGM and *Prior B* differ only in model regularization and as such this observation might be attributed to the *l*_1_ regularization of PLL-MGM. The latter induces sparseness in individual edges only and not edge groups. The node-wise MGM approach yielded an almost vanishing edge density of 0.041% and, therefore, was excluded from further comparison.

Finally, we compared the PriOmics networks to those obtained by two naive univariate screenings, one using a significance threshold for BH-adjusted p-values of *α* = 0.05 and a more restrictive model with *α* = 10*^−^*^28^. The latter threshold yielded an edge density of 1.83% and was chosen such that it agreed approximately with the PriOmics networks. The univariate approach with *α* = 0.05, however, yielded a dense network covering 62.44% of all possible edges, suggesting a huge number of false positive findings. It is noteworthy that while this univariate approach selected the majority of all possible edges, it did not capture all of those selected by PriOmics *Prior A* (7,425, Fig. 5E). The more restrictive univariate model with *α* = 10*^−^*^28^ showed a substantial disagreement to PriOmics *Prior A* with only 9.43% consistent edges (Fig. 5F). Moreover, the edges were inconsistent with respect to the underlying protein grouping, as suggested by an average edge proportion within protein groups of 35.78%. A Venn diagram illustrating the overlap of the inferred edges between all methods applied is shown in Suppl. Fig. S9.

##### Associations to Cell of Origin (COO)

The differentiation of DLBCLs according to their COO into GCB and ABC is used widely in patient stratification in randomized clinical trials [43]. Recently, proteomics based GCB/ABC subtyping was suggested [36]. Thus, we focused first on a PriOmics analysis of the proteomics and phenotypical data only (without transcriptomics data) to explore whether the COO first-order neighborhood (Fig. 6) resembled the signature suggested by [36] and whether it included additional potential proteomics markers for COO subtyping (the first-order neighborhood should contain the most directly associated proteins). The ABC neighborhood (Fig. 6A) of the PriOmics *Prior A* model comprises in total 19 peptides from 12 different proteins, where the tryptic peptide at the protein N-terminus of PDLIM1 ([1Ac]-TTQQIDLQPGPWGFR) was acetylated after co-translational cleavage of the N-terminal methionine by methionine aminopeptidase. Edges to COO are chosen to represent the differentiation of ABC vs. “unclassified”. The corresponding network visualizing the differentiation between GCB and “unclassified” is shown in Suppl. Figure S10A & B, where most edge signs are flipped. This is in line with the fact that those molecular features, which are positively associated with ABC, are usually negatively associated with GCB. In this analysis, “unclassified” DLBCLs are considered as an individual entity, as motivated by the unsupervised clustering performed in [44]. Interestingly, one of two variables, which differentiate both ABC and GCB with the same edge sign from unclassified DLBCL, is a peptide proteotypic for DNMT1 (DNA (cytosine-5)-methyltransferase 1). DNMT1, which belongs to a family of DNA methyltransferases that catalyze the transfer of the methyl moiety from S-adenosylmethionine to DNA, plays an important role in maintaining genome-wide methylation patterns during DNA replication [45]. DNMT1 is frequently expressed in both GCB and non-GCB DLBCL, but more strongly so in the former [46]. More importantly, however, GCB and ABC DLBCLs have been suggested to feature specific and distinct epigenetic alterations including DNA promoter methylation profiles [47]. In contrast, unclassified DLBCLs have been described to display nonspecific and, on average, lower DNA methylation levels. This appears to be consistent with the present finding that DNMT1 can distinguish both ABC and GCB DLBCLs from unclassified cases [48]. The second protein that differentiates both ABC and GCB with the same edge sign from unclassified DLBCL, is switch-associated protein 70 (SWAP70). SWAP70 is a Rac family guanine nucleotide exchange factor that independently of RAS, transduces signals from tyrosine kinase receptors to RAC [49]. Following B cell stimulation, expression of SWAP70 increases rapidly and it translocates from the cytoplasm to the nucleus, where it plays a role in immunoglobulin class switching, as well as to the plasma membrane, where it associates with the B cell receptor [50]. SWAP70 plays an important role in the regulation of cellular actin dynamics [51] and is required for polarization and homing of B cells into secondary lymphoid organs [52]. Its ability to activate RAC1 may help to understand why SWAP70 distinguishes both, ABC- and GCB-like DLBCL from unclassified cases. SWAP70 has been observed to regulate NOTCH2 [53], which is frequently mutated in unclassified DLBCL, defining together with BCL6 gene rearrangements the so-called BN2 subgroup of DLBCL [38]. However, even though BN2-like DLBCLs are associated with either NOTCH2 gain-of-function mutations or a relative increase in NOTCH2 expression, NOTCH2 is rarely activated in these tumors [54]. Hence, it appears reasonable, albeit to be proven, that decreased expression of SWAP70 and, thus, reduced activation of RAC1 and NOTCH2, is a common feature of unclassified, mostly BN2-like DLBCLs that distinguishes them from other DLBCLs.

**Figure 5:**
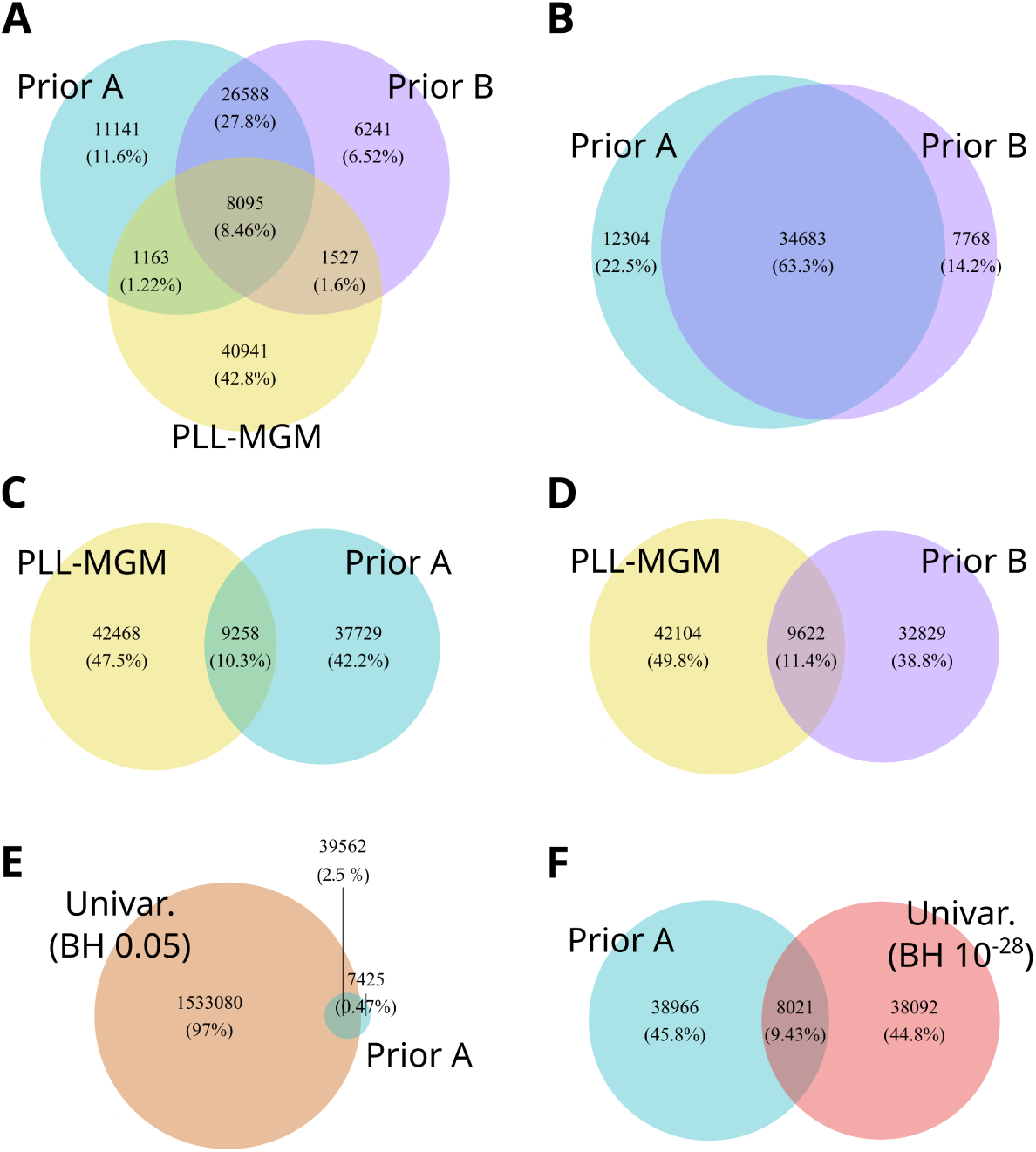
Venn diagrams display the overlap of inferred associations in DS1 between multiple network analysis strategies, i.e., PriOmics – *Prior A* & *B*, pseudo-log-likelihood-MGM, and two univariate approaches with BH-adjusted threshold cutoffs of 0.05 and 10*^−^*^28^.

**Figure 6:**
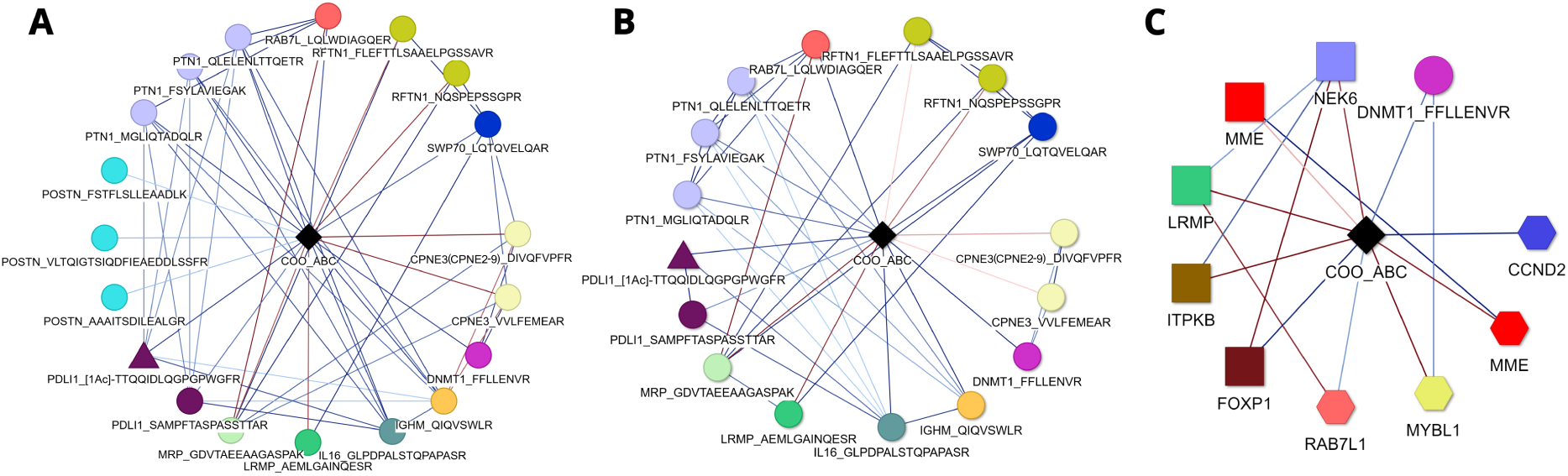
First order neighborhood networks of “ABC”-labeled DLBCLs compared to ‘unclassified’ DLBCLs. Figures A & B were based on PriOmics models calculated on the proteomics dataset (DS1) with *Prior A* or *B*, respectively. The model in Figure C included additional transcriptomics data (DS2) with *Prior A* for proteomics data and no prior for RNA data. Genes from the first data source are depicted as squares, genes from the second data source as hexagons, unmodified peptides as circles, peptides with a CTM or PTM as triangles, and discrete variables as diamonds. Adjacent nodes of the same color represent the protein affiliation. Edge colors indicate positive (blue) or negative (red) associations. Edge color intensity indicates the association strength. The node labels refer to the gene or protein name together with the amino acid sequence of the associated peptide, separated by an underscore. CTMs and PTMs are displayed in square brackets ([1Ac] = Acetylation). Edge weights are understood relative to the baseline COO=“unclassified”.

The majority of identified proteins in the COO first-order neighborhood are established COO markers, such as LRMP (*i.e. IRAG2*, *JAW1* ; Inositol 1,4,5-triphosphate receptor associated 2) [55, 56, 57, 34, 58, 59, 60], IL16 (Pro-interleukin-16) [55, 34, 33], IGHM (Immunoglobulin heavy constant mu) [34, 57, 36, 59, 60], and PTN1 (Tyrosine-protein phosphatase non-receptor type 1) [36, 55]. We further confirmed the sign of the association for each of the proteins. For instance, the downregulation (negative association) of PTN1 and IGHM in GCB-DLBCLs was already described by Reinders et al. [36]. Further, PDLI1 (i.e., PDLIM1; PDZ and LIM domain protein 1) was positively associated with the ABC subtype in the PriOmics model. Its corresponding gene *PDLIM1* was found to be upregulated in CD5+ cases that are associated with ABC [61]. However, only a minority of DLBCL patients showed CD5 expression. The loss of the closely related gene family member PDLIM2 has been reported as a defect commonly shared by classical Hodgkin lymphoma (cHL) and anaplastic large cell lymphoma (ALCL), indicating a potential tumor suppressing functionality [62]. Further studies showed that *PDLIM1* acts as cancer inducing regulator [63], *e.g.*, in breast cancer [64]. One of the two peptides representing PDLI1 was acetylated at the free alpha-amino group of the protein N-terminus after prior co-translational cleavage of the protein N-terminal methionine ([1Ac]-TTQQIDLQPGPWGFR; AA positions 2-17). This common protein modification affects approximately 80% of all human proteins including PDLI1 [40]. The minor discrepancy in the edge weights assigned to both peptides of PDLI1 (0.017 for SAMPFTASPASSTTAR (AA positions 123-138) and 0.055 for [1Ac]-TTQQIDLQGPGPWGFR) and the positive correlation obtained for *Prior B* (Fig. 6B) suggest, that the two peptides are technical replicates and that abundance of PDLI1 rather than its differential acetylation is a marker for COO. To further explore how results are affected by the selected prior, we compared the previous COO neighborhood to the one obtained by using *Prior B* (potential interrelationships between peptides of the same protein) (see Figure 6 A & B). The established edges to COO were almost identical to those obtained for *Prior A*. While the connections to three peptides of the protein POSTN disappeared, additional edges arose among grouped peptides (those are enforced to zero for *Prior A*). Note that the assumptions of both *Prior A* and *B* might be partially violated; most peptides affiliated with the same protein might be technical replicates (violating the assumptions of *Prior B*) but not necessarily all (violating the assumptions of *Prior A*). Thus, the observation that both analyses strongly coincide in their results serves as an important sanity check for PriOmics. For comparison, we also show the COO neighborhood established by PLL-MGM, which comprises 41 peptides, each representing a different protein (Suppl. Figure S11). Thus, in this case it remains to be elucidated whether the individual peptides or the underlying protein abundance are associated with COO. The latter information becomes directly accessible using PriOmics by considering the individual edge weights, as demonstrated by the previous example of PDLI1. As mentioned afore, *Prior B*, in contrast to *Prior A*, does not strictly assume that tryptic peptides derived from the same protein are technical replicates. Data-independent acquisition of tryptic peptides by SWATH-MS and their assignment to particular proteins is based on a peptide library generated previously using data-dependent acquisition followed by removal of tryptic peptides that are not unique, *i.e.*, that match more than one protein. Interestingly, closer inspection of the two PriOmics models in Figure 6 revealed a difference in association strength with the COO label between the two peptides believed to be proteotypic for CPNE3 in the PriOmics *Prior B*, but not the *Prior A* model. A renewed search against the UniProtKB/Swiss-Prot database [65] revealed that only the peptide VVLFEMEAR was indeed proteo-typic for CPNE3, while the peptide DIVQFVPFR was shared by the CPNE paralogs 2 thru 9. Thus, the two peptides represent different variables. Differences in strength of association with the COO label were also noted for RFTN1, with RFTN1-NQSPEPSSGPR showing a much stronger negative association with COO-ABC than RFTN1-FLEFTTLSAAELPGSSAVR. In this case, the difference in association may be due to phosphorylation of S220, the first serine residue from the N-terminal end of NQSPEPSSGPR, which is a reported phosphorylation site of RFTN1 [66]. Since phosphopeptides are typically present at substoichiometric levels when compared to nonmodified peptides of a protein and are also less efficiently ionized and prone to neutral loss of the phosphate group in collision-induced dissociation, which translates into reduced intensities of sequence informative fragment ions, failure to detect both the phosphorylated and the nonmodified peptide is common [67]. Nevertheless, differences in phosphorylation of a protein may be detected indirectly via differences in the amount of unmodified protein detected. In contrast to CPNE3 and RFTN1, the three tryptic peptides derived from PTN1 showed no differences in association strength with COO-ABC in the PriOmics *Prior B* model, thus indicating that these three peptides are indeed technical replicates.

Besides investigating the explorative property of the PriOmics models, we also tested their predictive performance. In detail, we estimated the probabilities of identifying the correct COO-status of the DLBCL specimens included in DS1 and DS2. To that end, it was necessary to find optimal cut-offs that define the probability ranges for each of the three COO subtypes. The PriOmics *Prior A* model resulted in an overall agreement rate of 0.79 on dataset DS1. Compared to the true COO labels of the 66 ABC-like, 100 unclassified, and 178 GCB-like DLBCLs, as determined by NanoString nCounter gene expression analysis using the Lymph2Cx DLBCL COO Classifier [68], the PriOmics *Prior A* model yielded ROC-AUCs of 0.906, 0.755, and 0.937, respectively, for ABC, unclassified, and GCB DLBCL specimens. A classification plot depicting the predicted and true COO labels for the 344 DLBCL specimens is shown in Suppl. Figure S12. The predicted COO labels with a PriOmics *Prior A* model on DS2 resulted in an overall agreement rate of 0.87, indicating a stronger predictive impact of gene transcripts than peptides, which does not come as a surprise, as SWATH-MS may yield precise results for repeated measurements of a specimen, but still be prone to matrix effects leading to differential suppression of ionization when analyzing a large collection of DLBCL tissues varying significantly in composition of lymphoma and stroma cells. We obtained ROC-AUCs of 0.964, 0.819, and 0.988, respectively, for ABC, unclassified, and GCB DLBCLs. The true class labels were distributed as follows: 63 ABC, 96 unclassified and 171 GCB specimens. The corresponding classification plot is shown in Suppl. Figure S13. Comparing the same COO-status predictions to labels determined via HTG, we achieved overall agreement rates of 0.722, 0.728 and 0.731, each respectively for *Prior A*, *B*, and PLL-MGM in DS1. Slighly higher agreement rates were obtained in DS2 with 0.749, 0.746 and 0.755, respectively for *Prior A*, *B*, and PLL-MGM.

**Table 4:**
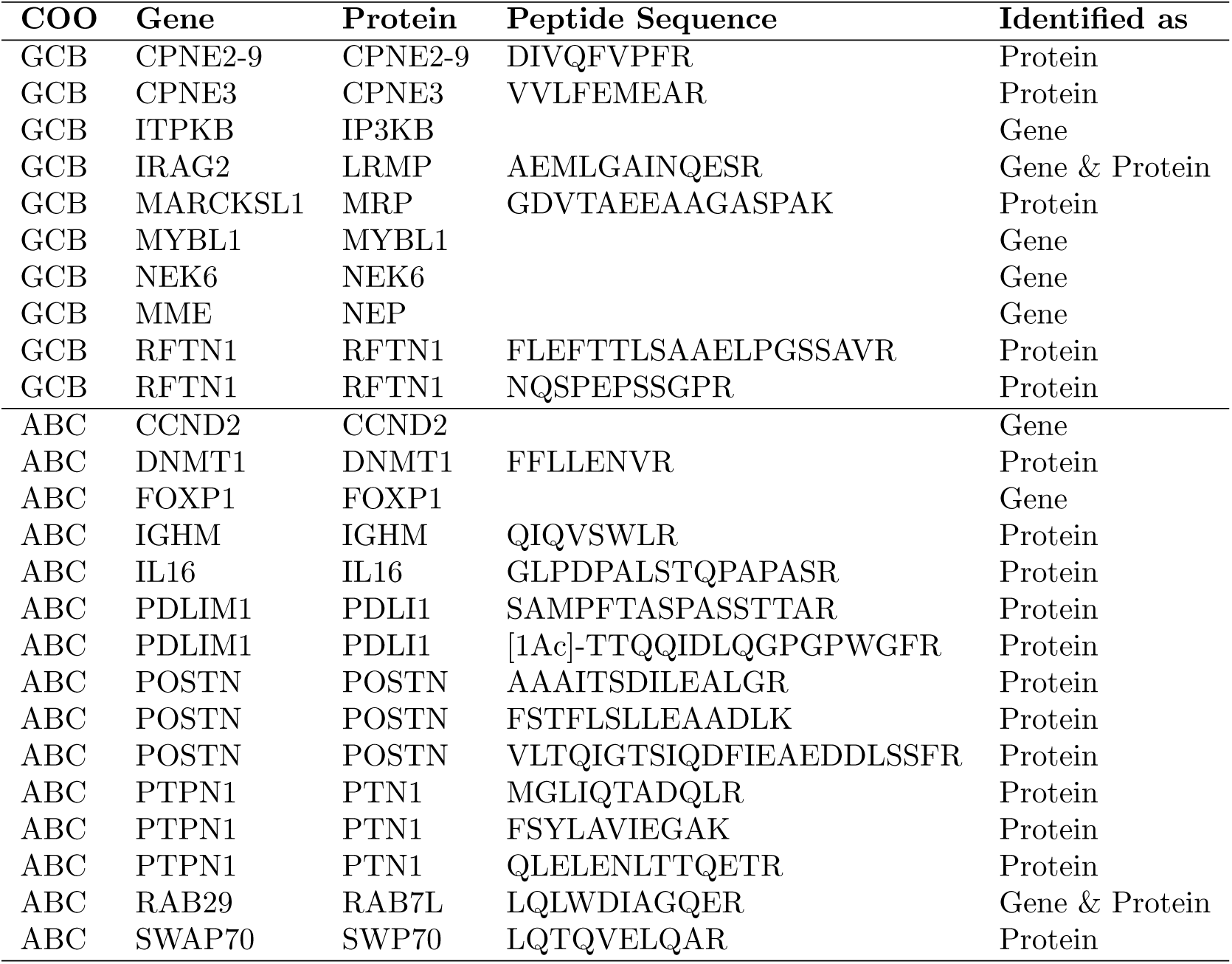
Associations between genes/proteins to GCB-DLBCL or ABC-DLBCL patients identified by PriOmics *Prior A* based on the proteomics dataset (DS1) and/or the combined transcriptomics+proteomics dataset (DS2)

Upon incorporation of additional transcriptomic features (dataset DS2), the network of the ABC neighborhood (Figure 6C) revealed associations for eight genes and one peptide. The gene *MME* (Neprilysin) occurred twice, due to its presence in both gene datasets. Again, most edge signs were flipped comparing ABC to GCB specimens (Suppl. Figure S10C) with “unclassified” specimens serving as reference. The only protein detected in both, DS1 and DS2, was DNMT1, which yielded congruent signs of the edge weights. Interestingly, RAB7L (*i.e.*, *RAB29*, Ras-related protein Rab-7L1) and LRMP were observed as protein entity in DS1 and as gene entity in DS2, which might indicate an increased predictive role of RNAs compared to respective proteins. Further, genes *CCND2* (G1/S-specific cyclin-D2), *MYBL1* (Myb-related protein A), *FOXP1* (Forkhead box protein P1), *ITPKB* (Inositol-trisphosphate 3-kinase B), and *NEK6* (Serine/threonine-protein kinase Nek6) were associated with COO. Out of the 20 genes for ABC/GCB classification proposed by Wright et al. [55, 69], PriOmics identified eight genes/proteins. The importance of FOXP1 as a main driver of DLBCL pathogenesis, especially in ABC-DLBCLs, was stated in multiple studies, though different FOXP1 isoforms might have different functions [70, 71]. Further, CCND2 was repeatedly described as a prognostic marker [72, 73]. In a combined study of three cohorts with 250 GCB-and ABC-DLBCL patients each, Liu et al. identified 87 differentially expressed genes (DEGs) in the ABC group [60]. Ten of these DEGs were also present in the PriOmics based networks of DS1 and/or DS2, namely MYBL1, MME, *MARCKSL1* (*i.e.*, MRP; MARCKS-related protein), ITPKB, LRMP, POSTN, IGHM, FOXP1, CCND2, and *RAB29*. All edge signs were in line with these findings, except POSTN. From the 19 COO-associated genes and proteins identified with PriOmics, SWP70, CPNE3, and six other markers were also found in a study comparing EZB-DLBCL subtypes [38] to non-EZB-DLBCLs [74]. An overview of the COO-associated genes and proteins identified by PriOmics *Prior A* is given in Table 4. Details about the particular edge strengths of the COO-associated variables are displayed in Supplementary Figures S15-S20.

##### Associations to clinical and phenotypical features

As a further sanity check, we explored the networks of the remaining categorical features, learned on the full dataset (proteomics + transcriptomics + phenotypic/clinical), as shown in Figure 7. The neighborhood of “gender male” in our PriOmics model was negatively associated with peptides proteotypic for proteins RS4X (40S ribosomal protein S4, X isoform, encoded by gene *RPS4X*) and DDX3X (ATP-dependent RNA helicase DDX3X; *i.e.*, a DEAD-box helicase family member), as well as to gene *NLGN4X* (Neuroligin-4, X-linked). In humans, *RS4X*, *DDX3X*, and *NLGN4X* are located on the X chromosome and their levels of expression have been observed to be higher in females, at least in certain tissues, due to escape from X-inactivation [75, 76]. DDX3X is involved in several tumor-related processes [77]. Further, studies showed that mutations in *DDX3X* can result in worse clinical outcomes in DLBCL patients [78]. Gong et al. showed a functional cooperation between *DDX3X* mutations and *MYC* -translocated DLBCL samples [79]. Lacroix et al. extended these findings by demonstrating the crucial role of *DDX3* for lymphomagenesis [80]. Female mice with *Ddx3x* loss-of-function mutations did not develop tumors, while male mice substituted the functional role of Ddx3x with a high expression of the paralog *Ddx3y* leading to lymphomagenesis.

**Figure 7:**
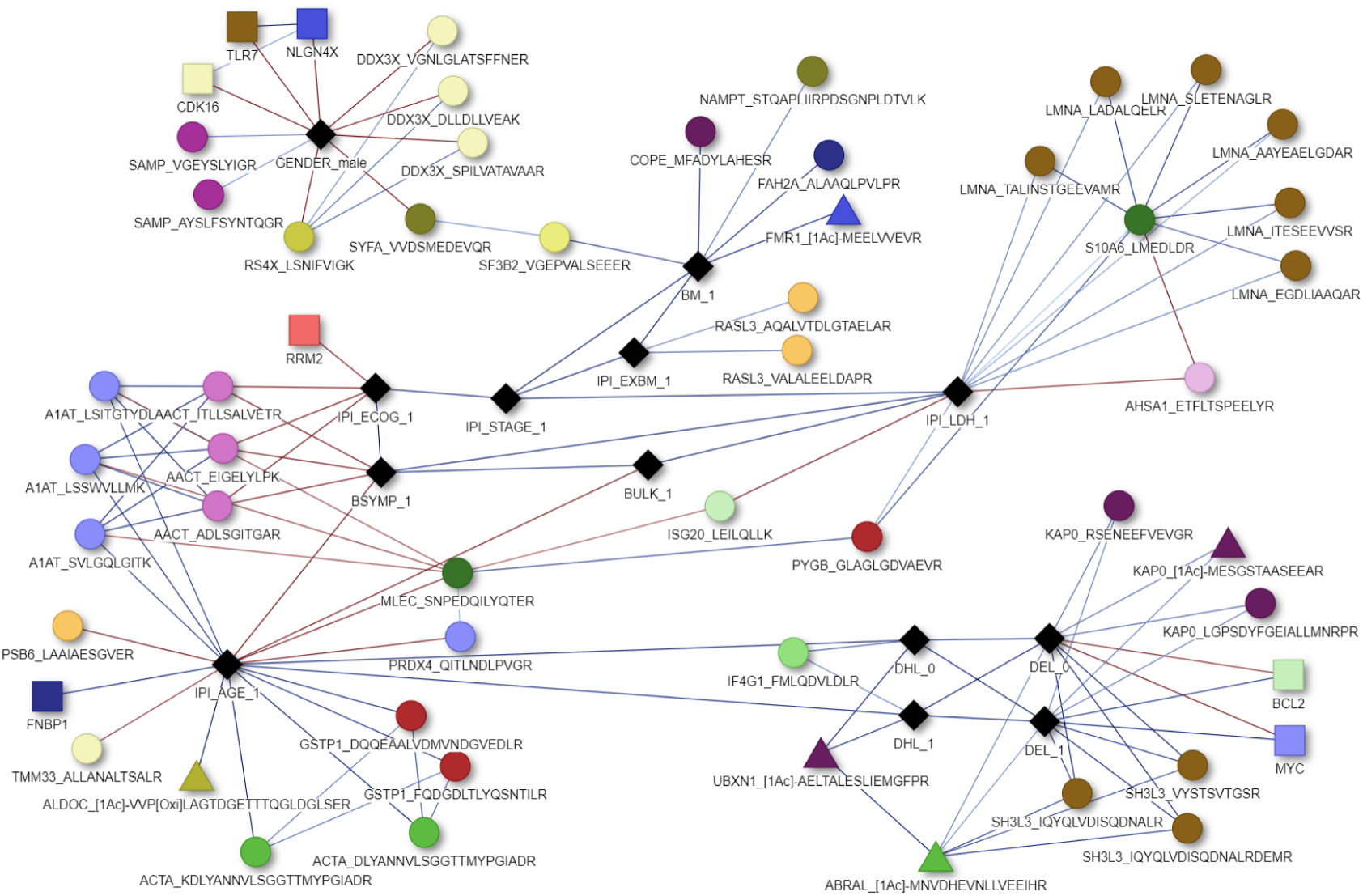
Association network from/to phenotypical and clinical variables in DS2. Feature names ending with “ 1” generally refers to the more severe patient status. Double-Hit lymphoma status (DHL) and Double-Hit expression status (DEL) were both compared to “unlabelled” cases as baseline (see details in Table 3).

Other features prognostic for poor outcome, such as high LDH serum levels (IPI LDH 1), presence of bulky disease (BULK 1), and B symptoms (BSYMP 1), were positively associated with each other. Moreover, multiple genes and proteins were associated with features distinctive for DLBCL. NAMPT (Nicotinamide phosphoribosyltransferase), for example, shows a positive relationship with bone marrow involvement [81, 82]. The strong relationship between the peptides of the proteins A1AT (*SERPINA1* ; Alpha-1-antitrypsin) and AACT (*SERPINA3* ; Alpha-1-antichymotrypsin) can be explained by their similar functional role as serine protease inhibitors. Older patients (*>* 60 years; IPI AGE 1) revealed negative associations to, e.g., presence of B symptoms (B SYMP 1) and presence of bulky disease (BULK 1). As expected, strong positive associations between high expression of the genes *MYC* and *BCL2* and the underlying Double-Hit expression status (DEL 1) were detected. Figure 7 generally outlines a holistic view on the interrelations between genes, proteins, peptides and their CTMs/PTMs, as well as phenotypical features. The specific association coefficients (*i.e.*, edge weights) of the phenotypical variables can be obtained from the heatmap in Suppl. Figure S21.

### PriOmics reveals post-translational modifications as confounding variables in high-throughput proteomics data

PriOmics *Prior A* assumes that the intensities of individual proteotypic peptides serve as a proxy for the underlying protein abundances, while *Prior B* is more appropriate for proteins that are represented by both nonmodified and co- or post-translationally modified peptides, as CTMs and PTMs play important roles in the regulation of the lifetime, structure, activity, location and interaction of proteins [83]. However, quantitative assessment of modified peptides by mass spectrometry may be affected by differences in ionization efficiency and stability in collision-induced dissociation compared to the nonmodified paralog [84]. This has to be taken into consideration when interpreting differences in correlation patterns between nonmodified and post-translationally modified peptides with other proteins. The entire proteome dataset contains only one pair of peptides representing both the nonmodified peptide and its modified paralog, which harbors a known PTM, namely phosphorylation of T160 of lamina-associated polypeptide 2 (LAP2B)(Figure 8). Nonphosphorylated LAP2B is an integral component of the inner nuclear membrane that binds to both lamin B (LMNB1) and chromatin. LAP2B plays an important role in nuclear envelope organization [85]. In metaphase, upon phosphorylation, LAP2B loses both, its LMNB1 and chromosome binding ability, resulting in the disassembly of nuclei. During late anaphase and early telophase, LAP2B becomes again localized to the surface of chromosomes, where it may facilitate reassembly of the nuclear envelope, while LMNB1 colocalizes with LAP2B at detectable levels only upon completion of cytokinesis. There is no clear negative or positive correlation between the intensities of nonphosphorylated and phosphorylated LAP2B and the intensities of the five LMNB1 peptides, which may be due to subtle differences in ionization efficiency or technical variance of quantitation. However, there is a clear negative or at least weaker association with the nuclear ribonucleoproteins HNRPU, ROA3, and SFPQ, the histone H1.5, and the far upstream element binding protein 1 (FUBP1), which is a master transcription regulator of a number of genes including MYC, and whose abundance has been already reported to decrease during mitosis [86]. The decreased levels of HNRPU and H1.5 observed with increased levels of phosphorylated LAP2B reflect most likely their known phosphorylation during mitosis. The scaffold attachment factor A (SAF-A) or HNRPU, a nuclear RNA-binding protein with RNA-to-DNA tethering activity, has been shown to remove, upon phosphorylation by Aurora-B, nuclear RNAs from the surface of prophase chromosomes to facilitate chromosome alignment and segregation in metaphase and anaphase, respectively [87]. Consequently, one would expect that the abundance of nonphosphorylated HNRPU to correlate negatively with phosphorylated LAP2B. This is particularly obvious for nonphosphorylated HN-RPU S*S*GPT*S*LFAVTVAPPGAR, which contains four reported phosphorylation sites, marked by an asterisk, at amino acid positions S187, S188, T191, and S192. As these sites become phosphorylated, the abundance of nonmodified HNRPU will decrease. Correspondingly, increasing phosphorylation of T10 of H1.5, which is contained in H15 [1Ac]-SETAPAETATPAPVEK, has been observed to start during prophase, and to increase to its highest level in metaphase [88]. Another known threonine phosphorylation site is T39, which is found in H15 KATGPPVSELITK and H15 ATGPPVSELITK [89]. However, its cell cycle-dependent phosphorylation has not been investigated yet. The same holds true for heterogeneous nuclear ribonucleoprotein A3 (ROA3), which has been implicated in the regulation of chromosome segregation [90], and whose serine residues at positions 355, 356, and 358, which are contained in the pep-tide ROA3 SSGSPYGGGYGSGGGSGGYGSR, have been reported to be phosphorylated under cellular stress and transformation-promoting conditions. But whether these sites will also be phosphorylated during mitosis, as described for HNRPU, remains to be elucidated. In summary, it appears that the PriOmics *Prior B* is capable of identifying PTMs that are of biological relevance.

**Figure 8:**
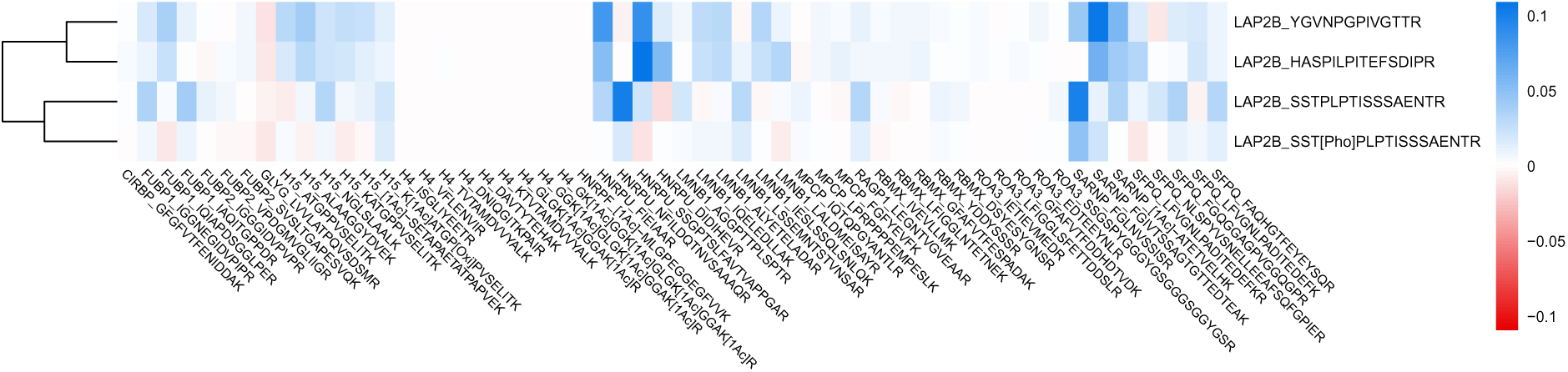
Association patterns of four peptides affiliated with protein LAP2B (Lamina-associated polypeptide 2, isoforms beta/gamma) in a PriOmics – *Prior B* model. Phosphorylation of the peptide SSTPLPTISSSAENTR (bottom row) results in alternating association scheme. Abbreviations: [1Ac], acetylation; [Pho], phosphorylation; [Oxi], oxidation.

## DISCUSSION

In recent years, the complexity of biomedical data analysis increased substantially as a consequence of new experimental possibilities to generate (multi-)omics data. These data are usually high dimensional comprising 10,000s and more molecular variables, generating both an increasingly detailed picture of biological systems but also challenges for data analysis. One major challenge are so-called spurious or erroneous associations, which are indirect associations as a consequence of one or multiple mediating variables. As seen in our analysis and in those of many others [9, 25], applying ordinary multiple testing strategies is not sufficient to control these false positive findings. Instead, variables have to be considered in their multivariate context to identify potential mediators of associations. PriOmics is designed to do so, while taking into account the specific nature of individual omics such as for untargeted high-throughput proteomics data. Moreover, PriOmics can model both continuous and categorical variables, which is a prerequisite to account for the increasing complexity of (multi-)omics data. We exemplified this by using PriOmics to integrate continuous variables such as peptide and RNA abundances with categorical variables, such as genomic alterations and phenotypical variables, in DLBCL specimens. In our analysis, we could demonstrate that PriOmics established rather sparse models where only 1.76 *−* 1.86% of all possible edges were inferred. Moreover, we could demonstrate that the joint analysis of multiple tiers of omics is a prerequisite to better resolve the underlying ground truth, as exemplified by edges established to the COO classification of DLBCLs. Here, we found that incorporating gene transcript levels in addition to peptide abundances, shielded COO associated peptides in favor of their respective RNA expression (seen for gene-protein pairs *IRAG2* -LRMP and *RAB29* -RAB7L, Figure 6). This may reflect the higher quantitative accuracy of measurements of gene expression over data independent acquisition-based tandem mass spectrometry of tryptic digests of proteomes, though it can not be excluded that COO is determined more strongly by gene rather than protein expression levels. Finally, we were able to retain highly relevant information about individual peptides without requiring further aggregation to protein abundances. This allowed us to reveal PTMs as potential confounding variables of respective protein abundances. One should also note that there is an increasing interest in PTMs as biomarkers and gateways for novel therapies in cancer treatment. PTMs occur by adding functional groups to the amino acid sequence of peptides, *i.e.*, phosphorylation, glycosylation, ubiquitination, and several others. Hence, they increase the complexity of the human proteome eminently. In cancer development, their importance has been stated repeatedly [91, 92, 93], further highlighting the need for robust and accurate methods for both measuring and analyzing large-scale proteomics data.

Although PriOmics incorporates many aspects relevant for state-of-the-art omics analysis, we want to recognize some limitations. PriOmics networks are undirected, meaning that PriOmics can not infer causal relationships. The latter can be clarified by prior knowledge. Moreover, there is ongoing research on establishing causal relationships from observational data, which could be also promising in combination with PriOmics. Further, we want to emphasize that although PriOmics can properly deal with both categorical and continuous data, it makes assumptions about the data, namely that the variables follow probability density Eq. (1). This might be inappropriate for continuous variables, which strongly deviate from normality. Here, data transformations could be an option. Moreover, technical copies of variables were incorporated by *a priori* removing specific edges and by using optimized regularization strategies, and not directly in terms of a joint probability density. Despite this limitation, it worked well in practice and offers a reasonable trade-off in analyses where it is not entirely clear if variables are copies or if they carry distinct biological information.

In summary, PriOmics facilitates the joint analysis of complex multi-omics data, especially proteomics data, while making *a priori* peptide to protein aggregation obsolete. The latter allows for retention of potentially relevant biological information such as those related to CTMs and PTMs. Moreover, PriOmics ability to model both continuous and categorical variables might open numerous possibilities to explore the complex interplay between the different tiers of regulation captured by nowadays omics.

## Supporting information

Supplement peptide list

## CODE AVAILABILITY

The algorithm is publicly available as user-friendly R ‘PriOmics’ in the GitHub repository ’https://github.com/roko4/PriOmics’ or as Python Code at ’https://github.com/roko4/PriOmics-Py’.

## ACKNOWLEDGEMENTS

We thank all people involved in the study ”German High Grade Lymphoma Study Group (DSHNHL)”, particularly Lorenz Trümper, the organizers and all participants for providing tumor samples. Further, we thank Anke Bauer and Sina Hillebrecht for performing HTG expression analyses.

