## Supplement for "PriOmics: integration of high-throughput proteomic data with complementary omics layers using mixed graphical modeling with group priors"

[1] Department of Medical Bioinformatics, University Medical Center Göttingen, Göttingen, Germany
[2] Peter L. Reichertz Institute for Medical Informatics of TU Braunschweig and Hannover Medical School, Hannover, Germany
[3] Chair and Institute of Functional Genomics, University of Regensburg, Regensburg, Germany
[4] Department of Clinical Pathology, Robert-Bosch-Krankenhaus, Stuttgart, Germany
[5] Dr. Margarete Fischer-Bosch Institute of Clinical Pharmacology, Stuttgart, and University of Tübingen, Germany
[6] Institute of Theoretical Physics, University of Regensburg, Regensburg, Germany
[7] Institute for Medical Informatics, Statistics and Epidemiology, University Leipzig, Leipzig, Germany
[8] Institute of Human Genetics, Ulm University & Ulm University Medical Center, Ulm, Germany
[9] Department of Statistical Bioinformatics, University of Regensburg, Regensburg, Germany
[†] shared senior authors

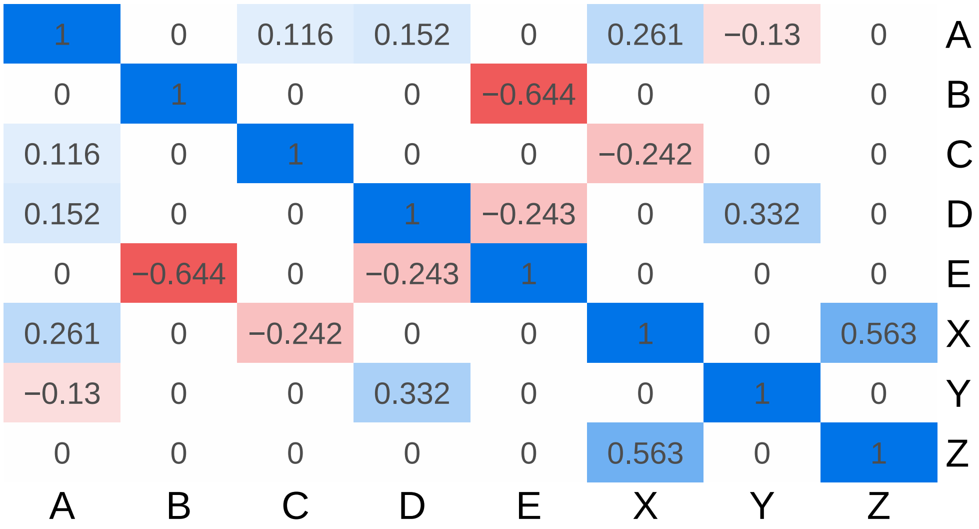


Figure S1: Simulated data ground truth. A to E represent five continuous features and X, Y, and Z denote discrete features with binary outcome.

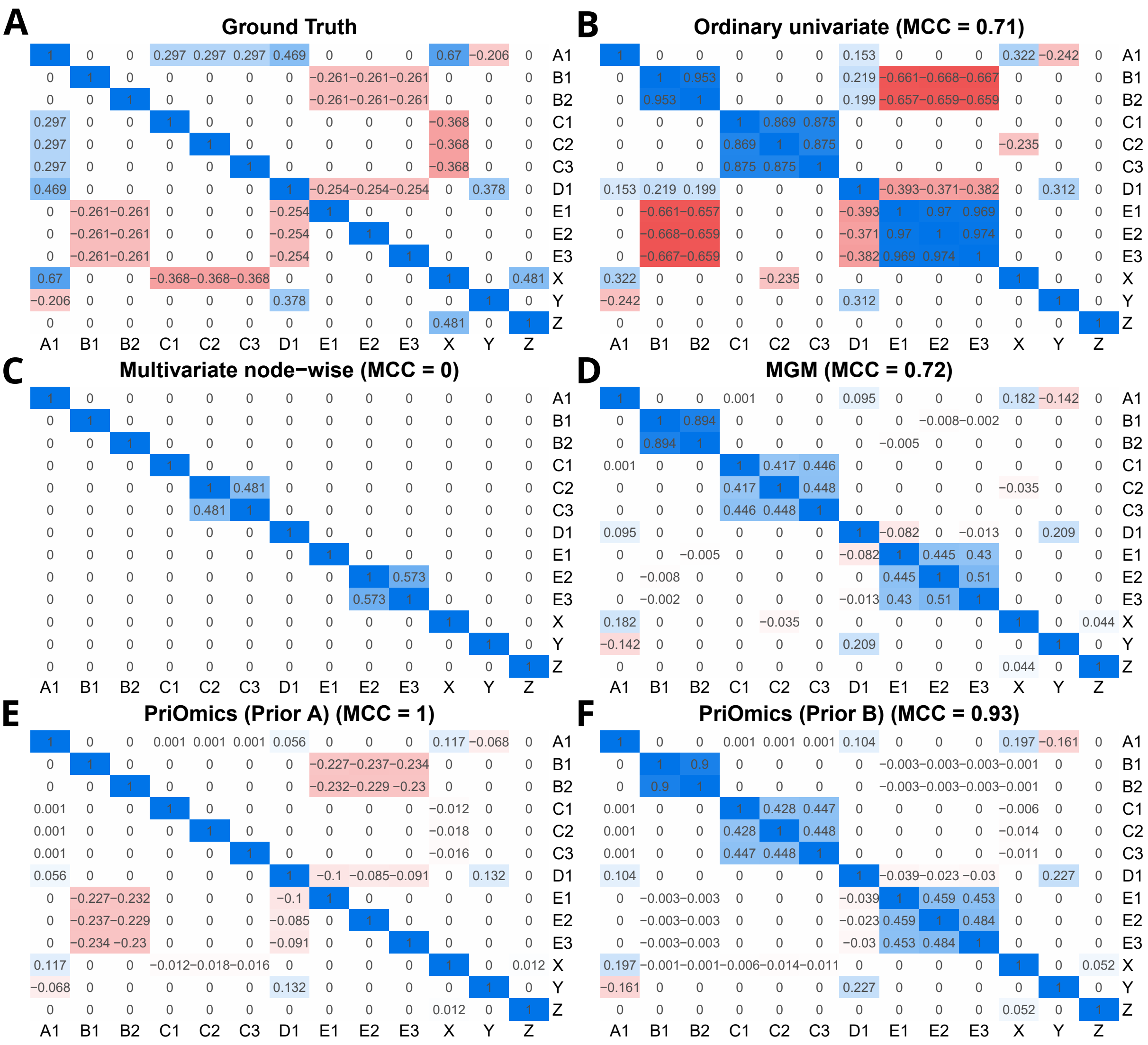


Figure S2: Illustration of network inference approaches on a simulated dataset. A1 to E3 represent five continuous features with a grouping factor indicated by the respective letters, resembling artificial peptides. X, Y, and Z denote artificial discrete features with binary outcome. The different methods were tested for their capability to reconstruct the network of a given dataset (A, ground truth). The association strength (parameter values of probability density Eq. (1)) is indicated by the numbers in the matrix. The sign of the association is displayed in blue for positive values and red for negative values. The different methodological approaches were compared by calculating the Matthews correlation coefficient (MCC), considering only edges between peptides of different proteins. Abbr.: MCC; Matthews correlation coefficient; MGM, Mixed graphical model.

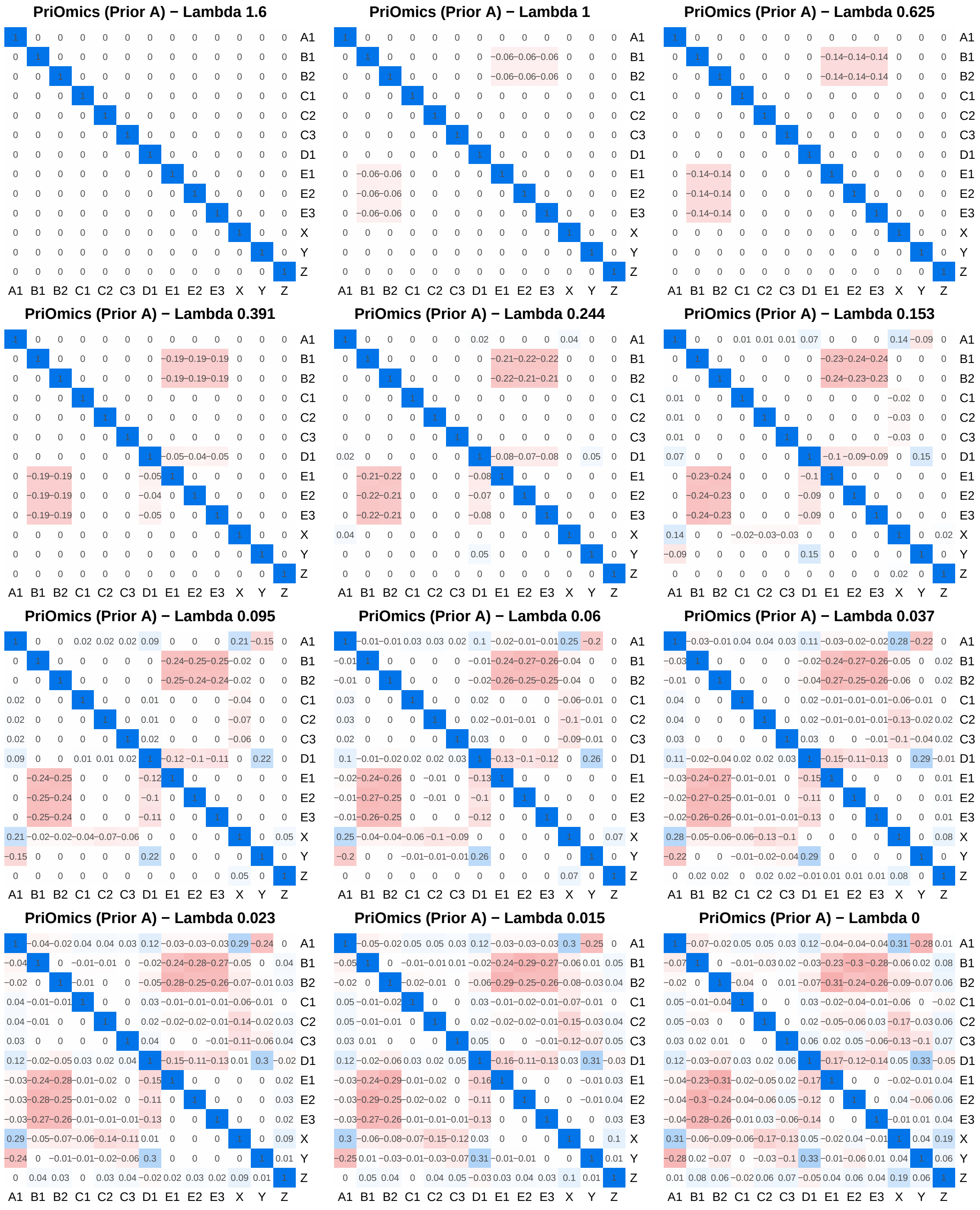


Figure S3: Regularization effects of a PriOmics - Prior A model with decreasing $\lambda$ on simulated data as described in Supplementary Figure S2.

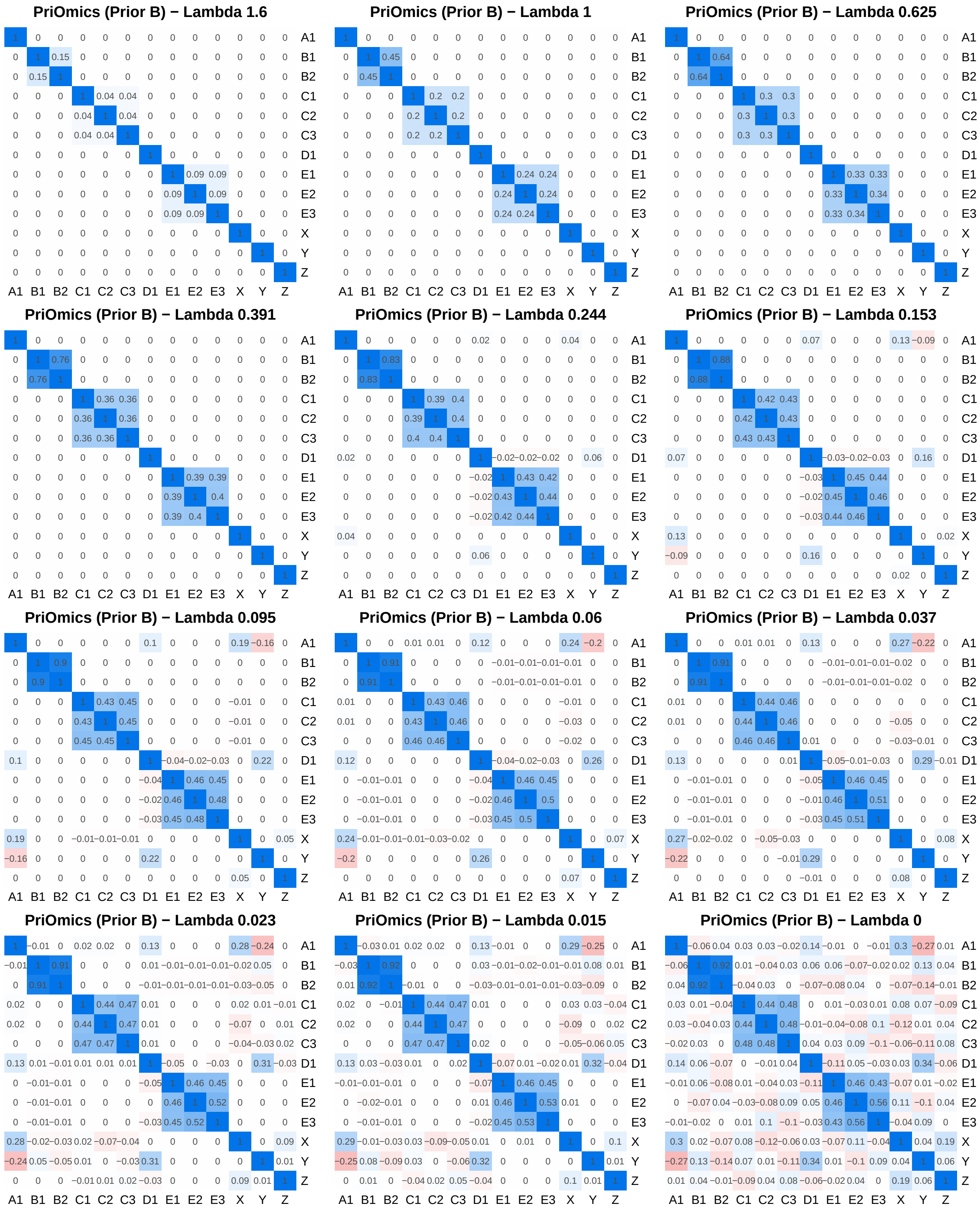


Figure S4: Regularization effects of a PriOmics - Prior B model with decreasing $\lambda$ on simulated data as described in Supplementary Figure S2.

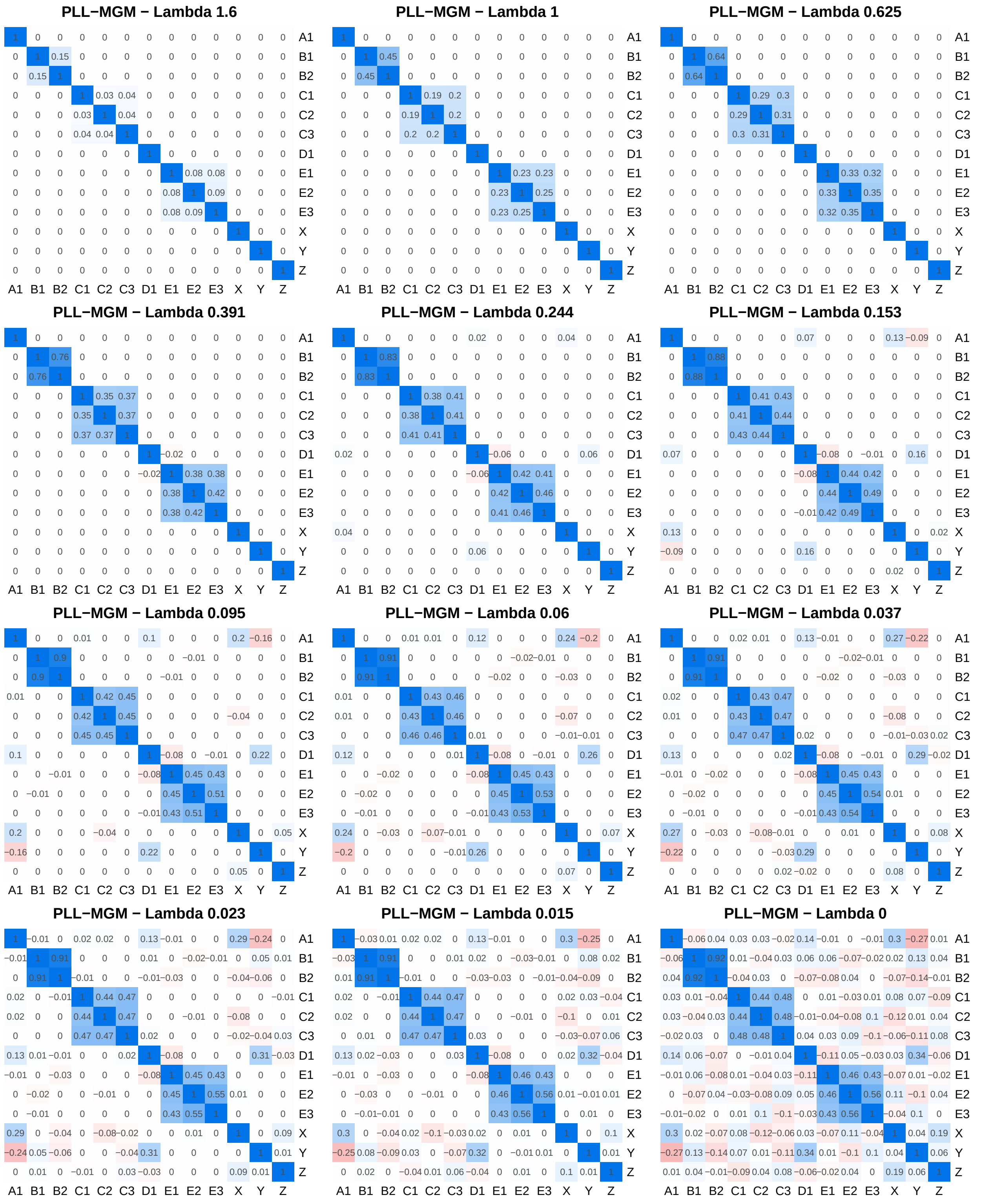


Figure S5: Regularization effects of a PLL-MGM with decreasing $\lambda$ on simulated data as described in Supplementary Figure S2.

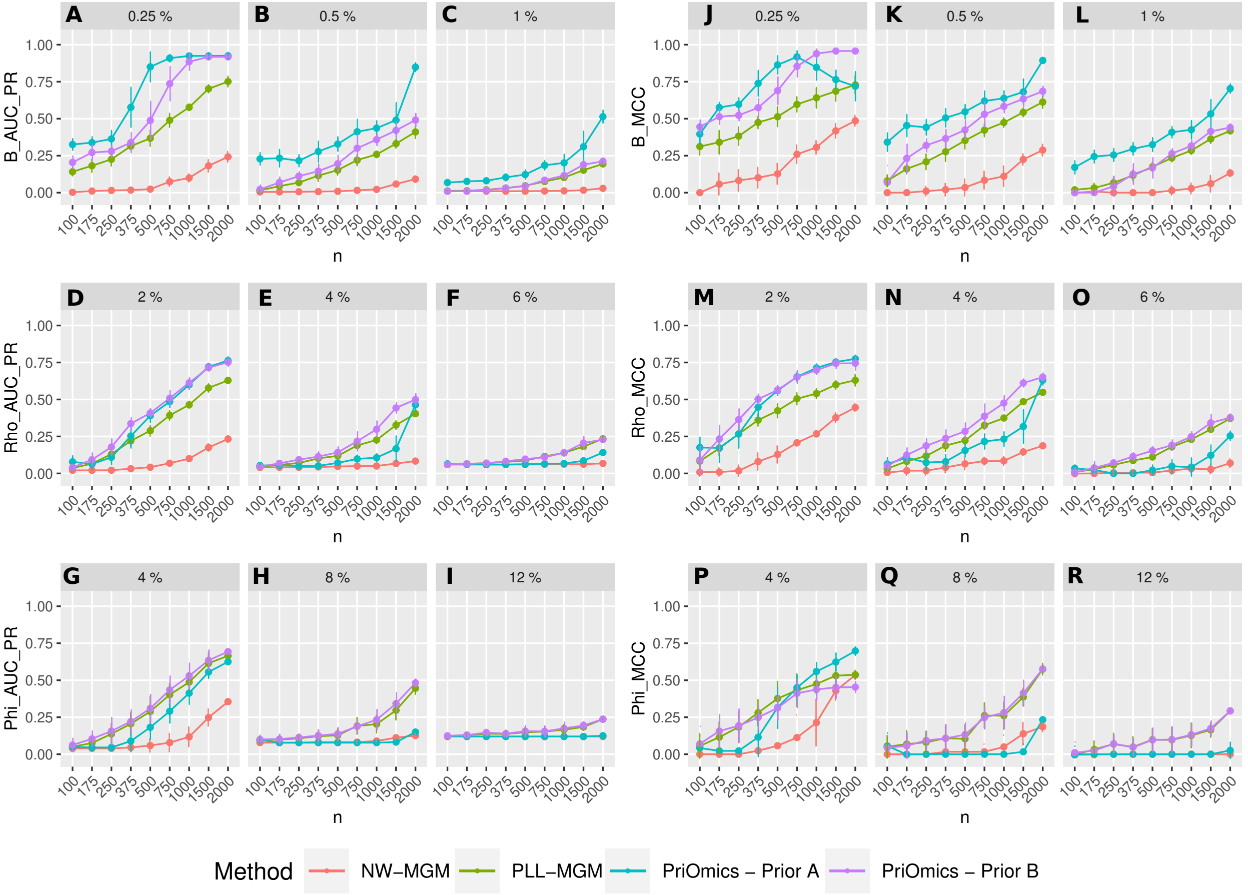


Figure S6: Performance comparison of a node-wise lasso regression MGM implementation ('NW-MGM'), a pseudo-log-likelihood MGM implementation ('PLL-MGM'), and the PriOmics implementations with different priors ('*Prior A*' and '*Prior B*') on simulated data. The plots on the left show the average AUC of the precision-recall curve (AUC_PR), while the right plots display the averaged MCCs. Each metric was calculated for an increasing amount of samples ($n$). The upper, middle and lower plots show the continuous-continuous (i.e., B), continuous-discrete (i.e. Rho), and discrete-discrete couplings (*i.e.*, Phi), respectively. Each scenario was performed with an increasing proportion of predefined edges ($\eta$). The columns correspond to three different simulation studies. The error bars indicate $\pm 1$ standard deviation.

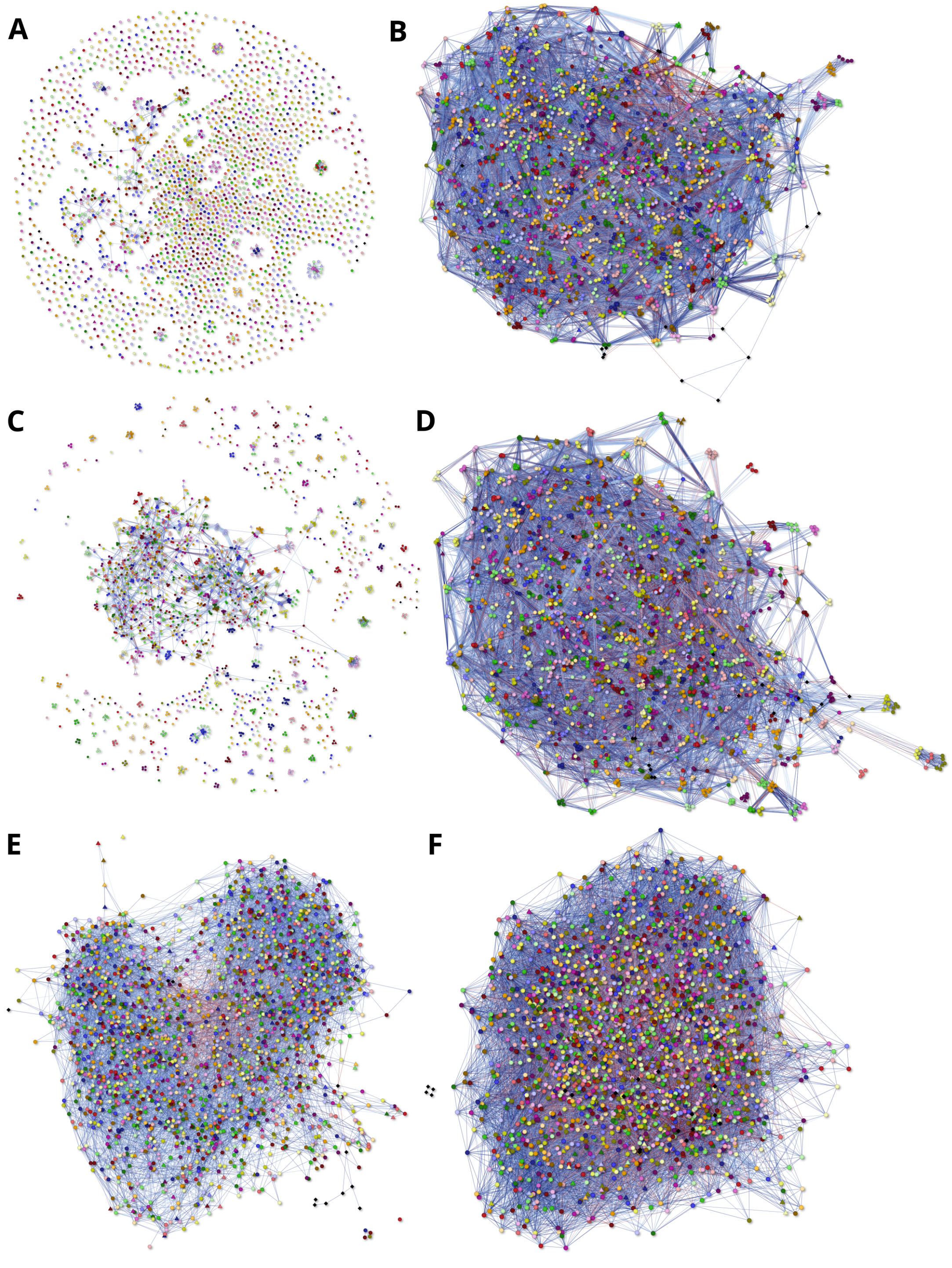


Figure S7: Network plots of DLBCL dataset DS1. Models were calculated with *Prior A* (Figures A & B), with *Prior B* (Figures C & D), and without priors (Figures E & F). Left: models selected by smallest EBIC. Right: models selected by smallest BIC.

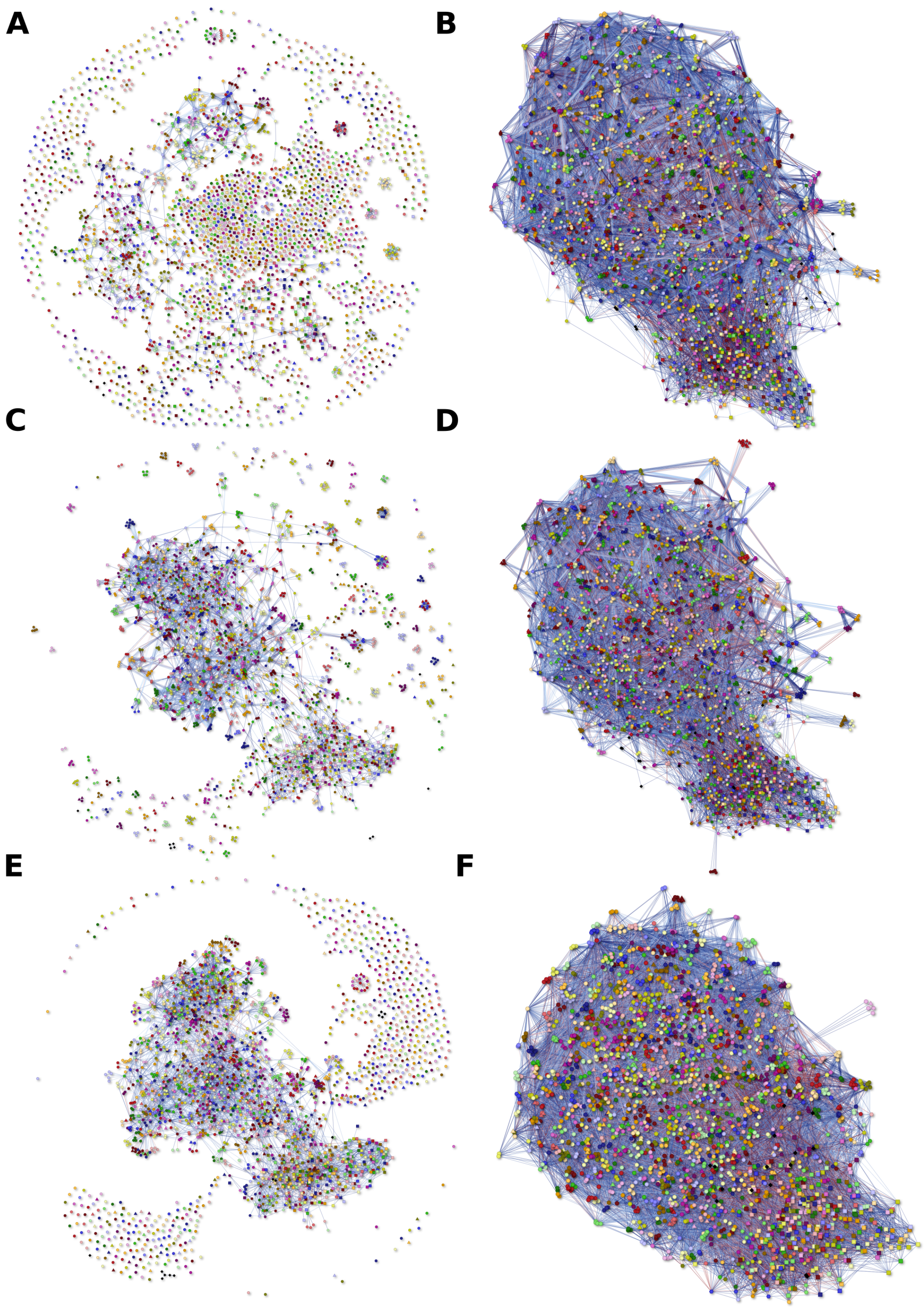


Figure S8: Network plots of DLBCL dataset DS2. Models were calculated with *Prior A* (Figures A & B), with *Prior B* (Figures C & D), and without priors (Figures E & F). Left: models selected by smallest EBIC. Right: models selected by smallest BIC.

Table S1: Number of edges in DLBCL data (DS1) for different $\lambda$ and different model priors.

| total | rate | B | Rho | Phi | COO | BIC | EBIC | Lambda | Method |
|---|---|---|---|---|---|---|---|---|---|
| 0 | 0.00 | 0 | 0 | 0 | 0 | 770742.58 | 770742.58 | 2.66 | Prior A |
| 0 | 0.00 | 0 | 0 | 0 | 0 | 770742.58 | 770742.58 | 2.35 | Prior A |
| 0 | 0.00 | 0 | 0 | 0 | 0 | 770742.58 | 770742.58 | 2.08 | Prior A |
| 0 | 0.00 | 0 | 0 | 0 | 0 | 770742.58 | 770742.58 | 1.84 | Prior A |
| 81 | 0.00 | 81 | 0 | 0 | 0 | 769980.55 | 771231.55 | 1.63 | Prior A |
| 216 | 0.01 | 216 | 0 | 0 | 0 | 766269.10 | 769605.11 | 1.44 | Prior A |
| 503 | 0.02 | 503 | 0 | 0 | 0 | 758900.86 | 766669.43 | 1.28 | Prior A |
| 1039 | 0.04 | 1039 | 0 | 0 | 0 | 747415.45 | 763462.26 | 1.13 | Prior A |
| 2180 | 0.09 | 2180 | 0 | 0 | 0 | 731921.53 | 765590.47 | 1.00 | Prior A |
| 4286 | 0.17 | 4285 | 0 | 4 | 0 | 711727.38 | 777968.71 | 0.89 | Prior A |
| 7261 | 0.29 | 7260 | 0 | 4 | 0 | 686697.63 | 798886.25 | 0.78 | Prior A |
| 11137 | 0.44 | 11132 | 0 | 13 | 0 | 659964.02 | 832092.63 | 0.69 | Prior A |
| 16032 | 0.64 | 16020 | 9 | 17 | 3 | 635870.34 | 883692.29 | 0.61 | Prior A |
| 21107 | 0.84 | 21084 | 24 | 25 | 6 | 612644.57 | 939032.54 | 0.54 | Prior A |
| 27293 | 1.08 | 27252 | 49 | 33 | 13 | 597381.42 | 1019540.55 | 0.48 | Prior A |
| 33214 | 1.32 | 33158 | 75 | 37 | 15 | 582932.64 | 1096770.14 | 0.42 | Prior A |
| 39469 | 1.57 | 39396 | 102 | 41 | 16 | 572825.49 | 1183484.37 | 0.38 | Prior A |
| 46987 | 1.86 | 46901 | 117 | 55 | 19 | 572113.29 | 1299130.81 | 0.33 | Prior A |
| 55338 | 2.19 | 55223 | 170 | 63 | 19 | 578057.46 | 1434545.96 | 0.29 | Prior A |
| 63860 | 2.53 | 63709 | 231 | 63 | 19 | 586257.29 | 1574749.67 | 0.26 | Prior A |
| 72605 | 2.88 | 72380 | 360 | 71 | 26 | 597050.50 | 1721577.76 | 0.23 | Prior A |
| 83374 | 3.31 | 83053 | 520 | 79 | 28 | 620246.94 | 1912207.70 | 0.20 | Prior A |
| 96040 | 3.81 | 95574 | 769 | 95 | 43 | 654651.92 | 2144085.67 | 0.18 | Prior A |
| 109835 | 4.36 | 109152 | 1137 | 95 | 66 | 695257.98 | 2400080.30 | 0.16 | Prior A |
| 127162 | 5.04 | 126307 | 1437 | 118 | 77 | 755350.18 | 2730110.94 | 0.14 | Prior A |
| 147100 | 5.83 | 146025 | 1839 | 122 | 85 | 829465.49 | 3115030.76 | 0.12 | Prior A |
| 169538 | 6.72 | 168136 | 2426 | 134 | 94 | 916654.88 | 3552964.05 | 0.11 | Prior A |
| 196612 | 7.80 | 194845 | 3062 | 132 | 115 | 1028642.28 | 4087249.57 | 0.10 | Prior A |
| 227024 | 9.00 | 224848 | 3793 | 136 | 144 | 1157924.88 | 4691264.27 | 0.09 | Prior A |
| 261071 | 10.36 | 258425 | 4650 | 136 | 172 | 1306089.44 | 5371243.70 | 0.08 | Prior A |
| 0 | 0.00 | 0 | 0 | 0 | 0 | 770742.58 | 770742.58 | 2.66 | Prior B |
| 0 | 0.00 | 0 | 0 | 0 | 0 | 770742.58 | 770742.58 | 2.35 | Prior B |
| 0 | 0.00 | 0 | 0 | 0 | 0 | 770742.58 | 770742.58 | 2.08 | Prior B |
| 20 | 0.00 | 20 | 0 | 0 | 0 | 770790.89 | 771099.78 | 1.84 | Prior B |
| 140 | 0.01 | 140 | 0 | 0 | 0 | 768610.87 | 770773.10 | 1.63 | Prior B |
| 351 | 0.01 | 351 | 0 | 0 | 0 | 760986.46 | 766407.46 | 1.44 | Prior B |
| 702 | 0.03 | 702 | 0 | 0 | 0 | 746595.61 | 757437.63 | 1.28 | Prior B |
| 1305 | 0.05 | 1305 | 0 | 0 | 0 | 726455.52 | 746610.55 | 1.13 | Prior B |
| 2251 | 0.09 | 2251 | 0 | 0 | 0 | 700022.17 | 734787.67 | 1.00 | Prior B |
| 3774 | 0.15 | 3773 | 0 | 4 | 0 | 668068.32 | 726402.08 | 0.89 | Prior B |
| 6027 | 0.24 | 6026 | 0 | 4 | 0 | 632755.97 | 725886.13 | 0.78 | Prior B |
| 8518 | 0.34 | 8511 | 3 | 13 | 1 | 594229.65 | 725924.64 | 0.69 | Prior B |
| 11707 | 0.46 | 11699 | 3 | 17 | 1 | 559235.70 | 740229.44 | 0.61 | Prior B |
| 15680 | 0.62 | 15666 | 9 | 25 | 3 | 529915.69 | 772393.86 | 0.54 | Prior B |
| 20566 | 0.82 | 20539 | 28 | 33 | 6 | 507673.33 | 825829.41 | 0.48 | Prior B |
| 25330 | 1.00 | 25295 | 41 | 37 | 7 | 486549.81 | 878422.34 | 0.42 | Prior B |
| 30796 | 1.22 | 30751 | 56 | 41 | 9 | 471317.54 | 947748.54 | 0.38 | Prior B |
| 37645 | 1.49 | 37568 | 99 | 63 | 14 | 465983.89 | 1048703.73 | 0.33 | Prior B |
| 44354 | 1.76 | 44258 | 134 | 63 | 16 | 461075.61 | 1147659.50 | 0.29 | Prior B |
| 52613 | 2.09 | 52491 | 181 | 63 | 17 | 466317.97 | 1280782.07 | 0.26 | Prior B |
| 61251 | 2.43 | 61065 | 288 | 71 | 22 | 474751.33 | 1423412.42 | 0.23 | Prior B |

| | | | | | | | | | |
|---|---|---|---|---|---|---|---|---|---|
| 71195 | 2.82 | 70947 | 396 | 75 | 25 | 491204.13 | 1594217.25 | 0.20 | Prior B |
| 82070 | 3.26 | 81707 | 585 | 95 | 29 | 513286.47 | 1785709.98 | 0.18 | Prior B |
| 94682 | 3.76 | 94193 | 814 | 95 | 38 | 545048.62 | 2013848.56 | 0.16 | Prior B |
| 109804 | 4.36 | 109127 | 1138 | 118 | 50 | 590858.12 | 2295665.00 | 0.14 | Prior B |
| 127321 | 5.05 | 126453 | 1480 | 124 | 57 | 649337.54 | 2627109.97 | 0.12 | Prior B |
| 147219 | 5.84 | 146060 | 1994 | 140 | 68 | 720602.75 | 3009380.46 | 0.11 | Prior B |
| 170210 | 6.75 | 168672 | 2660 | 140 | 82 | 808174.61 | 3456468.68 | 0.10 | Prior B |
| 197221 | 7.82 | 195288 | 3378 | 140 | 106 | 917245.62 | 3987698.83 | 0.09 | Prior B |
| 227376 | 9.02 | 224987 | 4175 | 144 | 134 | 1042437.00 | 4583946.52 | 0.08 | Prior B |
| 0 | 0.00 | 0 | 0 | 0 | 0 | 770742.58 | 770742.58 | 2.66 | PLL-MGM |
| 0 | 0.00 | 0 | 0 | 0 | 0 | 770742.58 | 770742.58 | 2.35 | PLL-MGM |
| 0 | 0.00 | 0 | 0 | 0 | 0 | 770742.58 | 770742.58 | 2.08 | PLL-MGM |
| 44 | 0.00 | 44 | 0 | 0 | 0 | 769771.22 | 770450.77 | 1.84 | PLL-MGM |
| 306 | 0.01 | 306 | 0 | 0 | 0 | 757836.60 | 762562.61 | 1.63 | PLL-MGM |
| 810 | 0.03 | 810 | 0 | 0 | 0 | 727905.77 | 740415.79 | 1.44 | PLL-MGM |
| 1502 | 0.06 | 1502 | 0 | 0 | 0 | 685037.30 | 708234.90 | 1.28 | PLL-MGM |
| 2440 | 0.10 | 2440 | 0 | 0 | 0 | 634774.27 | 672458.78 | 1.13 | PLL-MGM |
| 3609 | 0.14 | 3609 | 0 | 0 | 0 | 580811.45 | 636550.54 | 1.00 | PLL-MGM |
| 4975 | 0.20 | 4974 | 0 | 4 | 0 | 526498.36 | 603380.93 | 0.88 | PLL-MGM |
| 6496 | 0.26 | 6495 | 0 | 4 | 0 | 473526.75 | 573900.36 | 0.78 | PLL-MGM |
| 8167 | 0.32 | 8159 | 5 | 13 | 1 | 423216.23 | 549505.65 | 0.69 | PLL-MGM |
| 9959 | 0.40 | 9950 | 5 | 17 | 1 | 376020.21 | 530032.46 | 0.61 | PLL-MGM |
| 11771 | 0.47 | 11756 | 12 | 25 | 2 | 331749.07 | 513885.70 | 0.54 | PLL-MGM |
| 13779 | 0.55 | 13751 | 30 | 33 | 6 | 291416.57 | 504766.48 | 0.48 | PLL-MGM |
| 15900 | 0.63 | 15863 | 42 | 37 | 10 | 254313.44 | 500529.17 | 0.42 | PLL-MGM |
| 18157 | 0.72 | 18111 | 56 | 41 | 10 | 220316.32 | 501529.22 | 0.38 | PLL-MGM |
| 20608 | 0.82 | 20541 | 85 | 59 | 11 | 189515.87 | 508984.72 | 0.33 | PLL-MGM |
| 23368 | 0.93 | 23264 | 151 | 59 | 11 | 161883.14 | 524426.61 | 0.29 | PLL-MGM |
| 26416 | 1.05 | 26275 | 217 | 63 | 12 | 136833.48 | 546961.37 | 0.26 | PLL-MGM |
| 30095 | 1.19 | 29891 | 325 | 71 | 17 | 115776.40 | 583543.05 | 0.23 | PLL-MGM |
| 34458 | 1.37 | 34153 | 501 | 71 | 23 | 98366.55 | 634675.76 | 0.20 | PLL-MGM |
| 39563 | 1.57 | 39155 | 681 | 87 | 29 | 84410.56 | 701000.11 | 0.18 | PLL-MGM |
| 45817 | 1.82 | 45273 | 928 | 99 | 33 | 75858.72 | 790937.66 | 0.16 | PLL-MGM |
| 53104 | 2.11 | 52405 | 1209 | 99 | 41 | 71409.84 | 900978.63 | 0.14 | PLL-MGM |
| 61876 | 2.45 | 60983 | 1558 | 118 | 50 | 73614.25 | 1041349.26 | 0.12 | PLL-MGM |
| 82838 | 3.29 | 81504 | 2352 | 118 | 80 | 90837.77 | 1387771.65 | 0.10 | PLL-MGM |
| 82838 | 3.29 | 81504 | 2352 | 118 | 80 | 90837.77 | 1387771.65 | 0.10 | PLL-MGM |
| 95811 | 3.80 | 94248 | 2753 | 126 | 97 | 110103.79 | 1610178.77 | 0.09 | PLL-MGM |
| 109945 | 4.36 | 108128 | 3210 | 126 | 113 | 133771.04 | 1855273.39 | 0.08 | PLL-MGM |

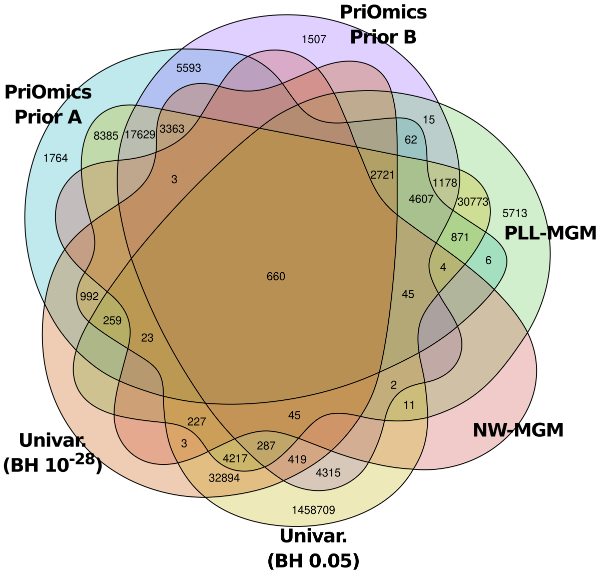


Figure S9: The Venn diagram displays the overlap of inferred associations in DLBCL dataset DS1 analyzed by all methods.

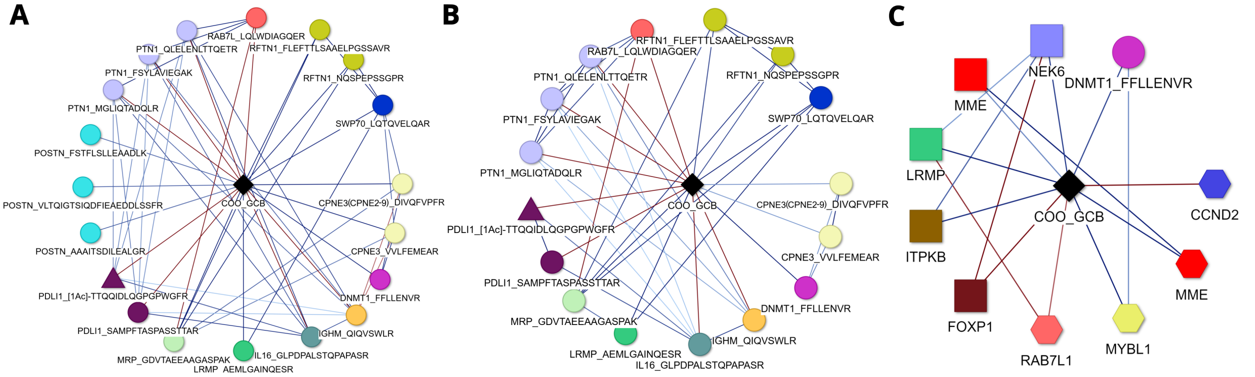


Figure S10: First order neighborhood networks of "GCB"-labeled DLBCLs compared to 'unclassified' ones. Figures A & B were based on PriOmics models calculated on the proteomics dataset (DS1) with *Prior A* and *Prior B*, respectively. The model in Figure C additionally included transcriptomics data (DS2) with *Prior A* for proteomics data and no Prior for RNA data. Genes from DS1 are depicted as squares, genes from DS2 as hexagons, unmodified peptides as circles, peptides with a co- or post-translational modification as triangles, and discrete variables as diamonds. Adjacent nodes of the same color represent the protein affiliation. Edge colors indicate positive (blue) or negative (red) associations. Edge color intensity indicates the association strength. The node labels refer to the gene or protein name, with the amino acid sequence of the associated peptide separated by an underscore. Co- and post-translational modifications are displayed in square brackets. Edge weights are understood relative to the baseline COO="unclassified".

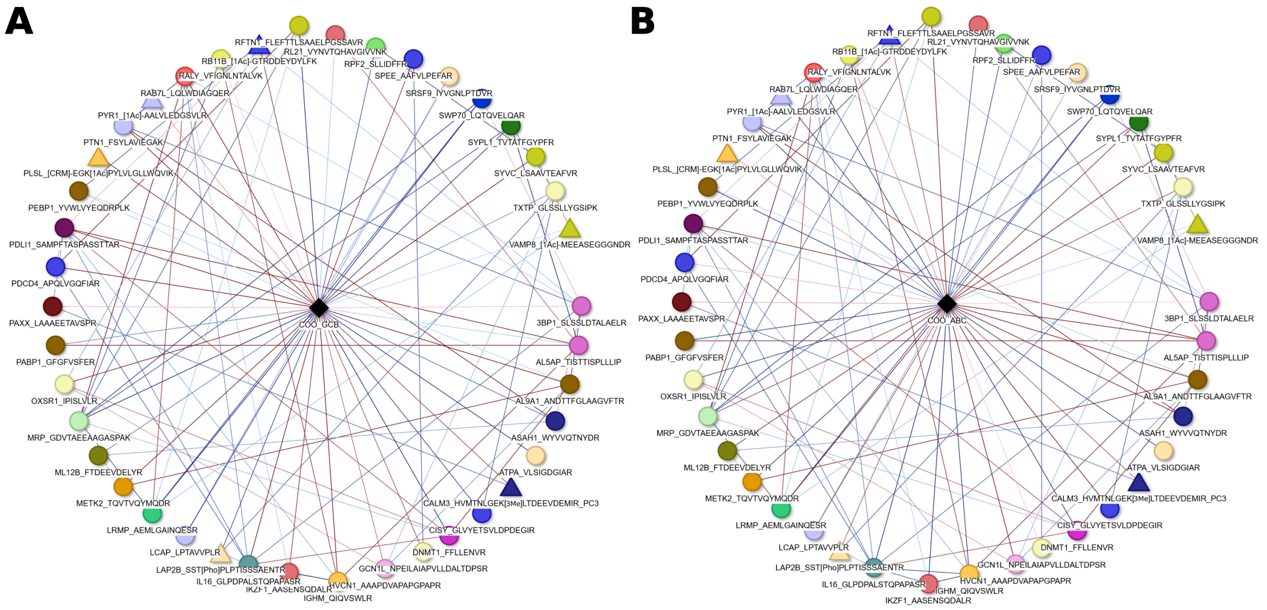


Figure S11: First order neighborhood networks of (A) "GCB" and (B) "ABC" labeled DLBCLs compared to 'unclassified' DLBCLs. Both networks are based on a PLL-MGM calculated on the proteomics dataset (DS1). Unmodified and modified peptides are depicted as circles and triangles, respectively, discrete variables as diamonds. Edge colors indicate positive (blue) or negative (red) associations. Edge color intensity indicates the association strength. The node labels refer to the protein name and the amino acid sequence of the associated peptide, separated by an underscore. The network contains a total of 6 peptides that harbor four different modifications given in square brackets: [1Ac] Acetylation, [CRM] Carbamylation, [3Me] tri-Methylation, and [Pho] Phosphorylation. VAMP8_[1Ac]-MEEASEGGGNDR, PYR1_[1Ac]-AALVLEDGSVLR, and RR11B_[1Ac]-GTRDDEYDYLFK are representatives of the most common co-translational modification found in the human proteome, namely the irreversible acetylation of the protein N-terminus without (VAMP8) or with prior cleavage (PYR1 and RB11B) of the N-terminal methionine by methionine aminopeptidases. PLSL_[CRM]-EGK[1Ac]PYLVLGLLWQVIK features a not yet reported acetylation of the epsilon-amino group of the lysine residue K220 and a carbamylation of the free alpha-amino group of N-terminal glutamic acid due to a non-enzymatic reaction with isocyanic acid derived from urea used for protein extraction. CALM3_HVMTNLGEK[3Me]LTDEEVDEMIR_PC3 features a previously reported N6,N6,N6-trimethylation of the epsilon-amino group of K116, while PC3 indicates that the peptide was triply charged [M+3H] in contrast to most other peptides that were only doubly charged [M+2H]. Finally, LAP2B_SST[Pho]PLPTISSSAENTR is phosphorylated at T160, a known phospho-threonine site of lamina-associated polypeptide 2. Edge weights are understood relative to the baseline COO="unclassified".

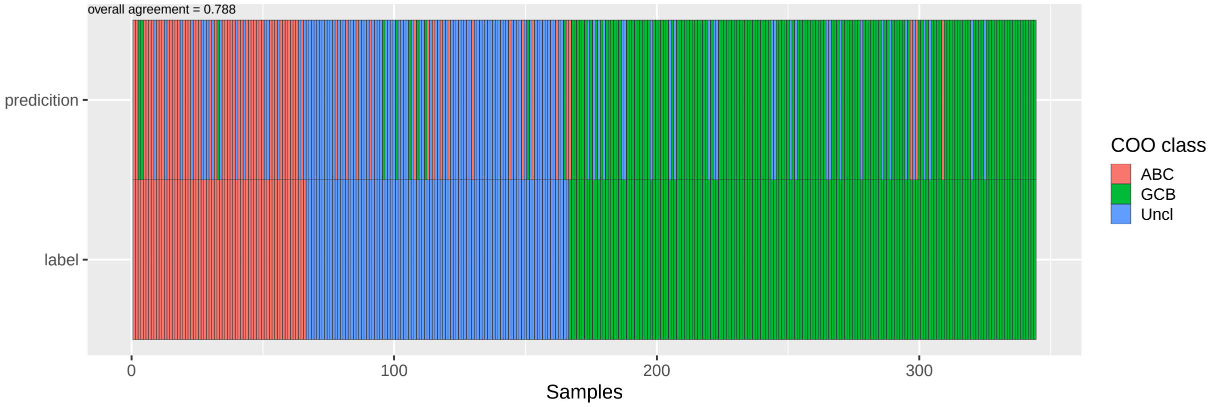


Figure S12: Performance of a PriOmics *Prior A* model for predicting the COO class, learned on the proteomics dataset DS1. The lower row displays the true COO label of the 344 DLBCL specimens as determined by nCounter (NanoString Technologies) gene expression analysis; the upper row displays the predicted COO label.

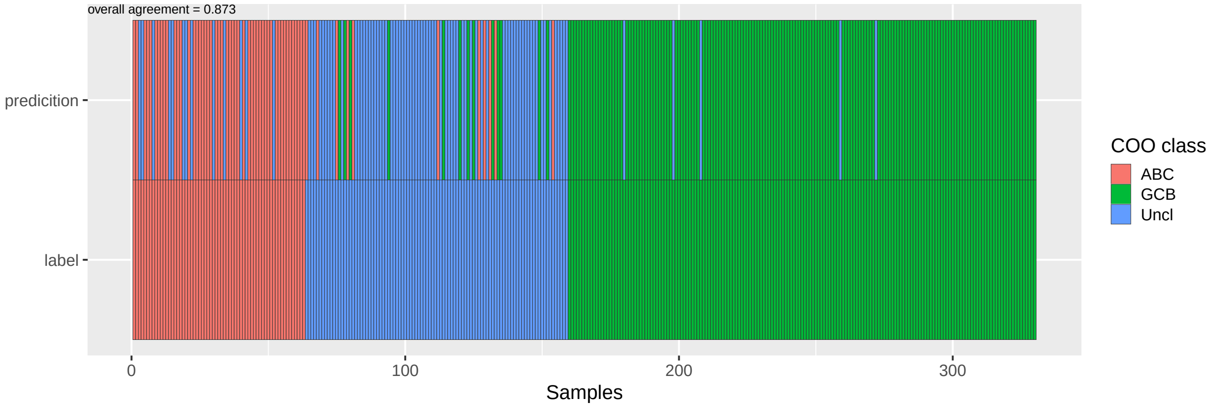


Figure S13: Performance of a PriOmics *Prior A* model for predicting the COO class, learned on the proteomics dataset DS2. The lower row displays the true COO label of the 344 DLBCL specimens as determined by nCounter (NanoString Technologies) gene expression analysis; the upper row displays the predicted COO label.

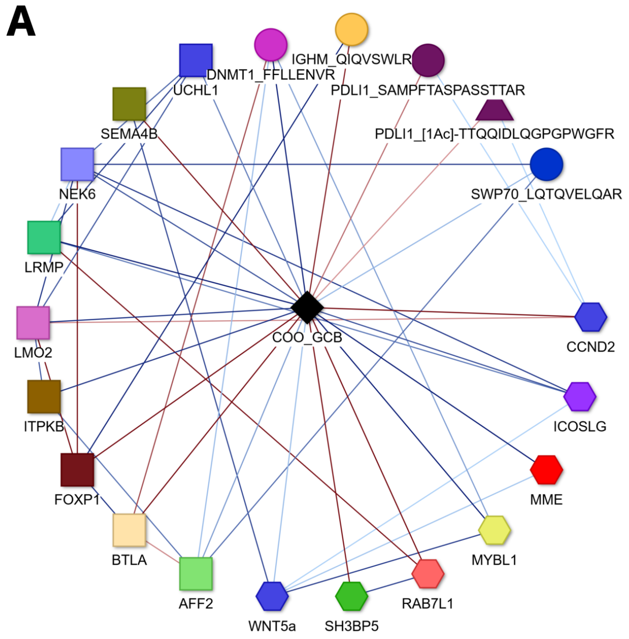


Figure S14: First order neighborhood networks of (A) "GCB" and (B) "ABC" labeled DLBCLs compared to 'unclassified' DLBCLs. Both networks are based on a PLL-MGM calculated on dataset (DS2). Genes from the first source are depicted as squares, genes from the second source as hexagons, unmodified peptides as circles, peptides with a co- or post-translational modification as triangles, and discrete variables as diamonds. Adjacent nodes of the same color represent the protein affiliation. Edge colors indicate positive (blue) or negative (red) associations. Edge color intensity indicates the association strength. The node labels refer to the protein name and the amino acid sequence of the associated peptide, separated by an underscore. Co- and post-translational modifications are displayed in square brackets. Edge weights are understood relative to the baseline COO="unclassified".

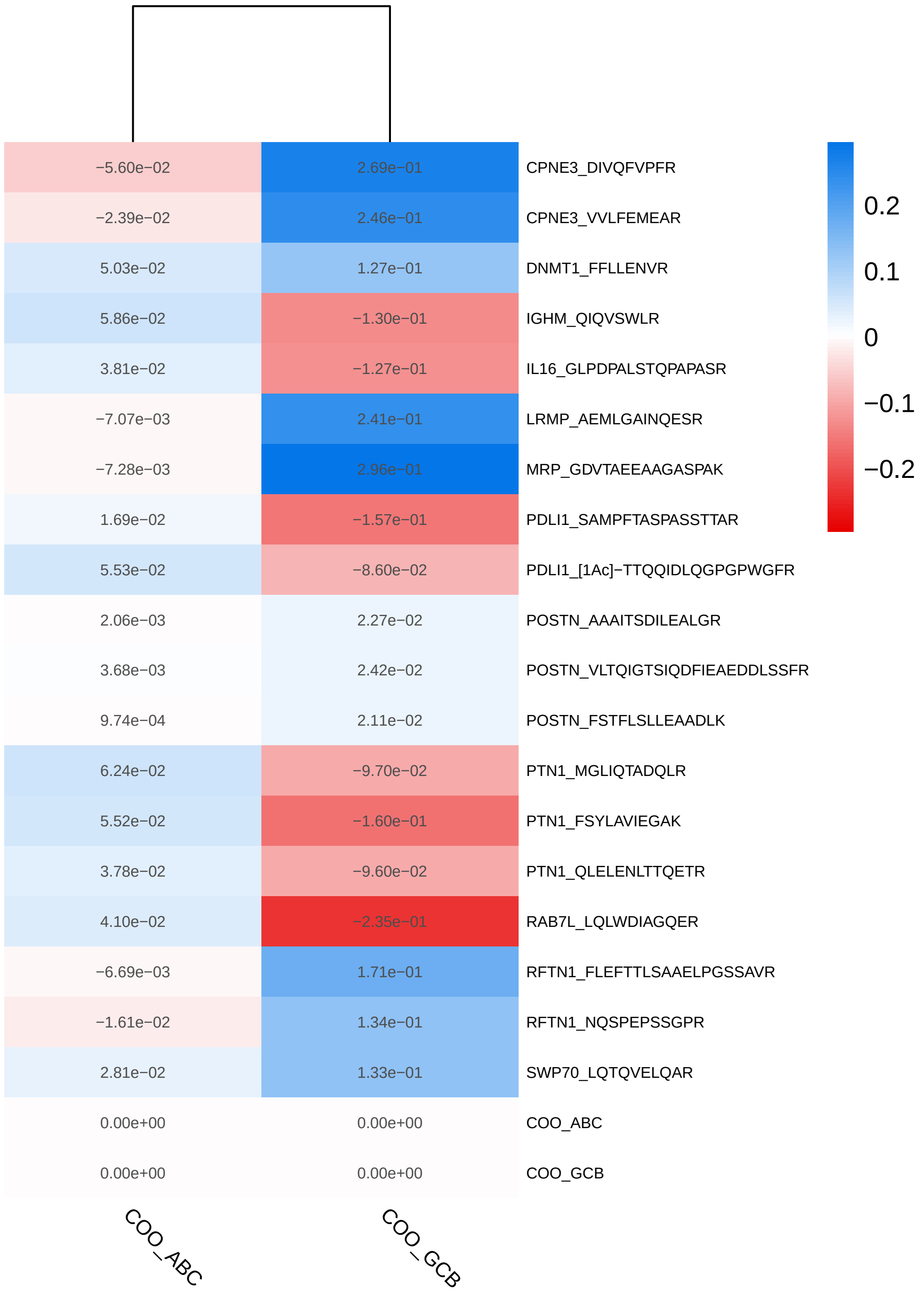


Figure S15: First order neighborhoods of the COO variables ABC (left) and GCB (right) with unclassified cases used as baseline, calculated with PriOmics *Prior A* based on the proteomics dataset DS1. The values refer to the respective edge strengths.

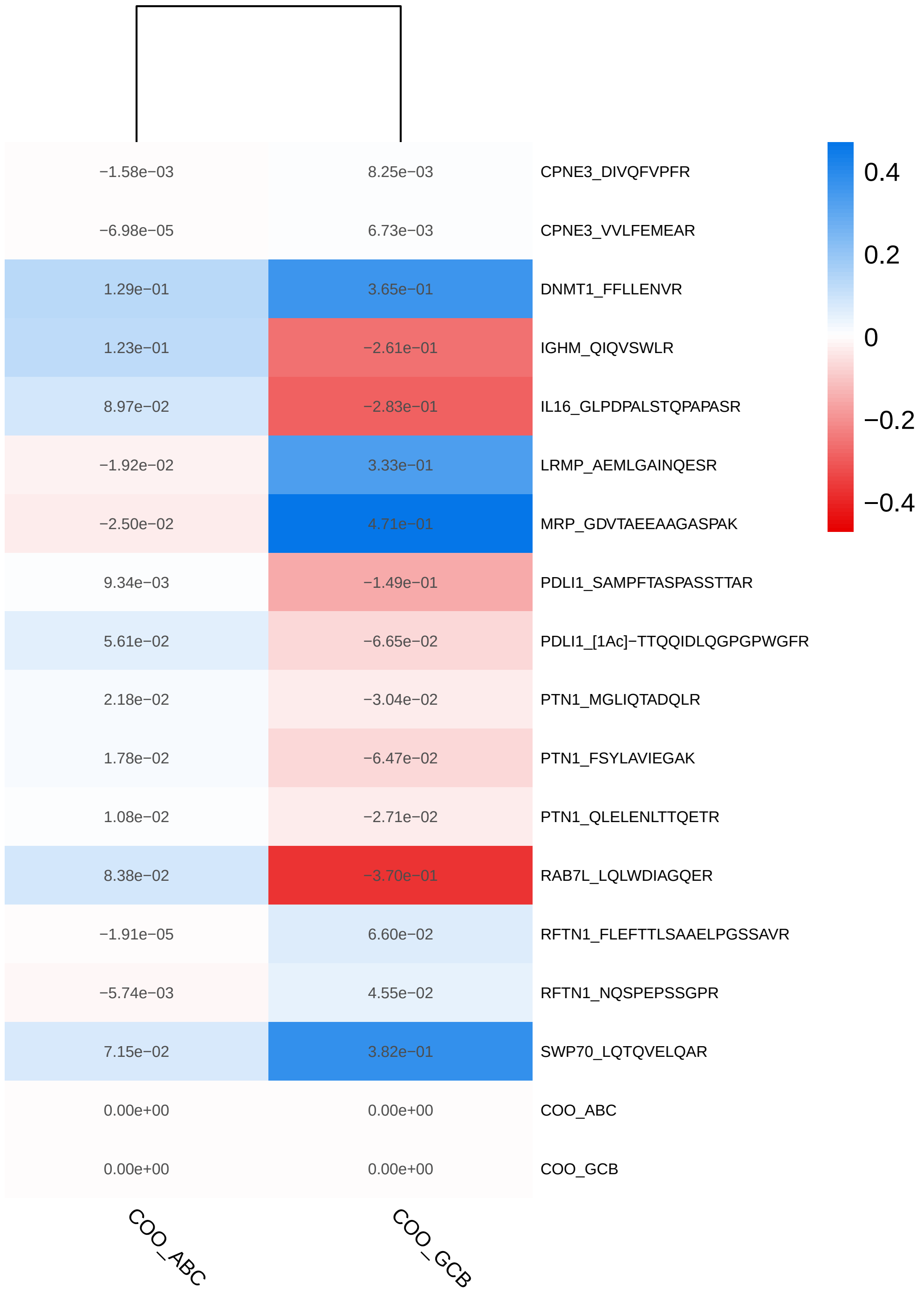


Figure S16: First order neighborhoods of the COO variables ABC (left) and GCB (right) with unclassified cases used as baseline, calculated with PriOmics *Prior B* based on the proteomics dataset DS1. The values refer to the respective edge strengths.

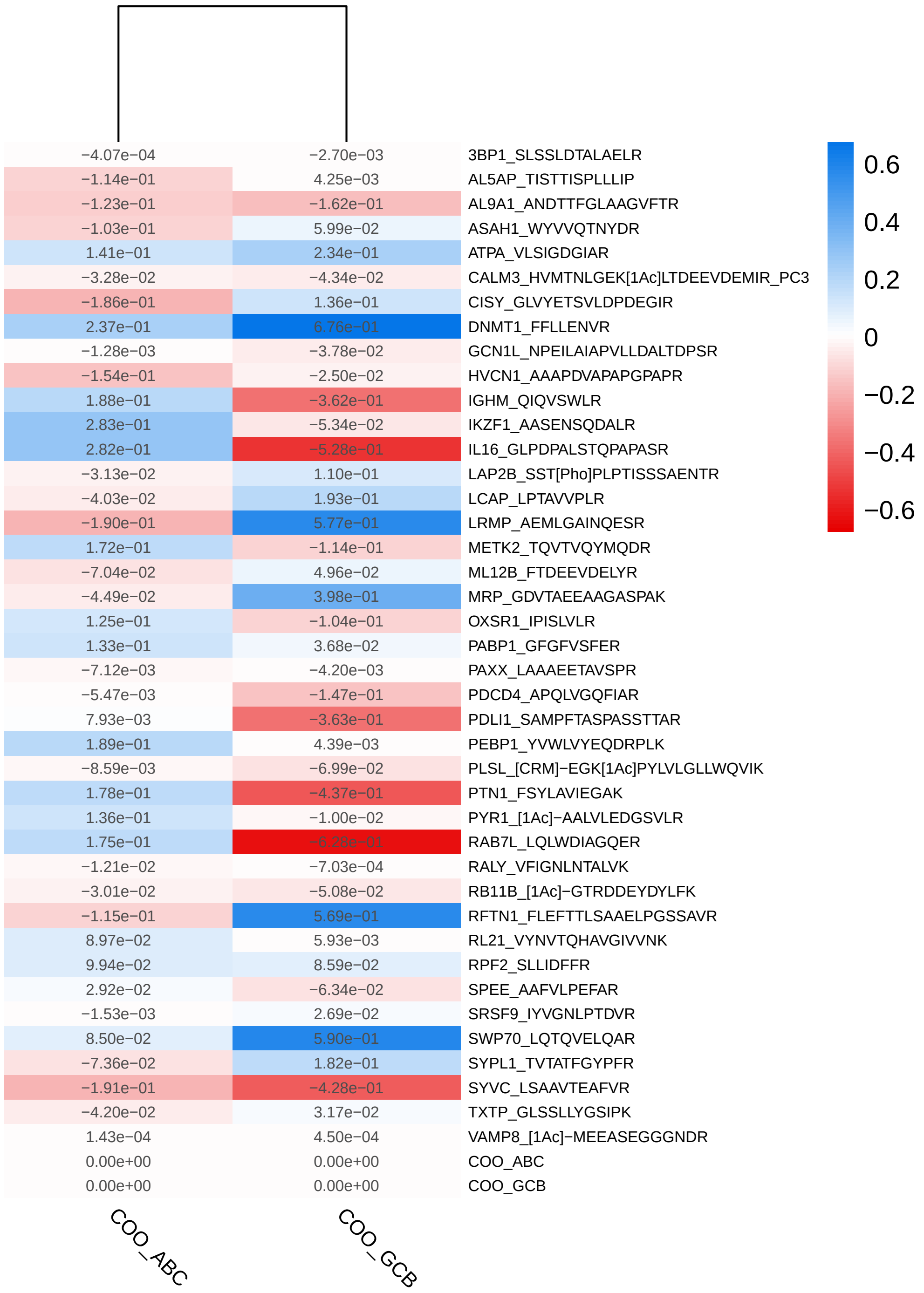


Figure S17: First order neighborhoods of the COO variables ABC (left) and GCB (right) with unclassified cases used as baseline, calculated with PLL-MGM based on the proteomics dataset DS1. The values refer to the respective edge strengths.

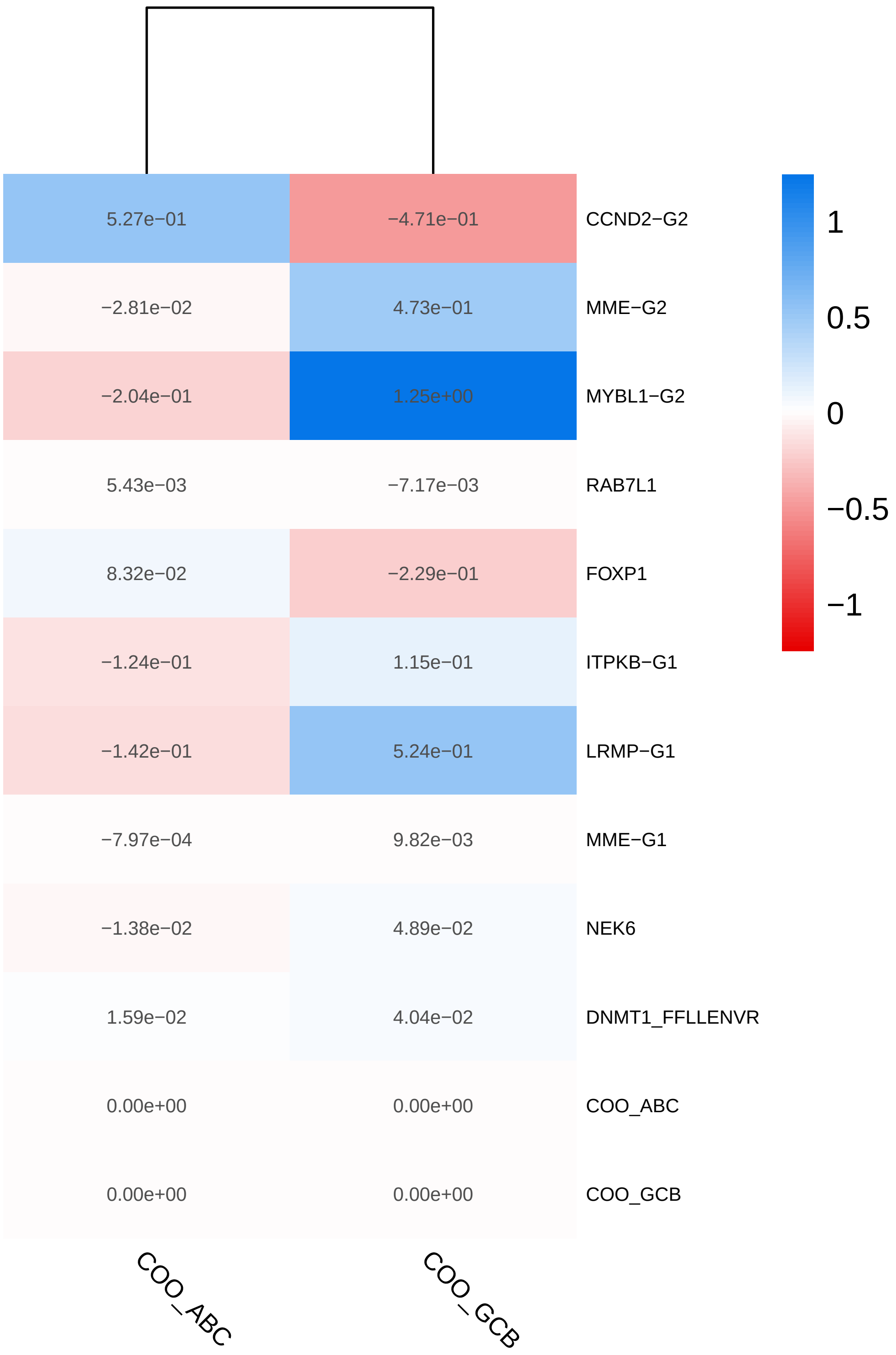


Figure S18: First order neighborhoods of the COO variables ABC (left) and GCB (right) with unclassified cases used as baseline, calculated with PriOmics *Prior A* based on the combined transcriptomics and proteomics dataset DS2. The values refer to the respective edge strengths. Gene names ending with "-G1" and "-G2" indicate the origin of the two transcriptomics datasets, i.e. measured either with NanoString nCounter (G1) or HTG (G2).

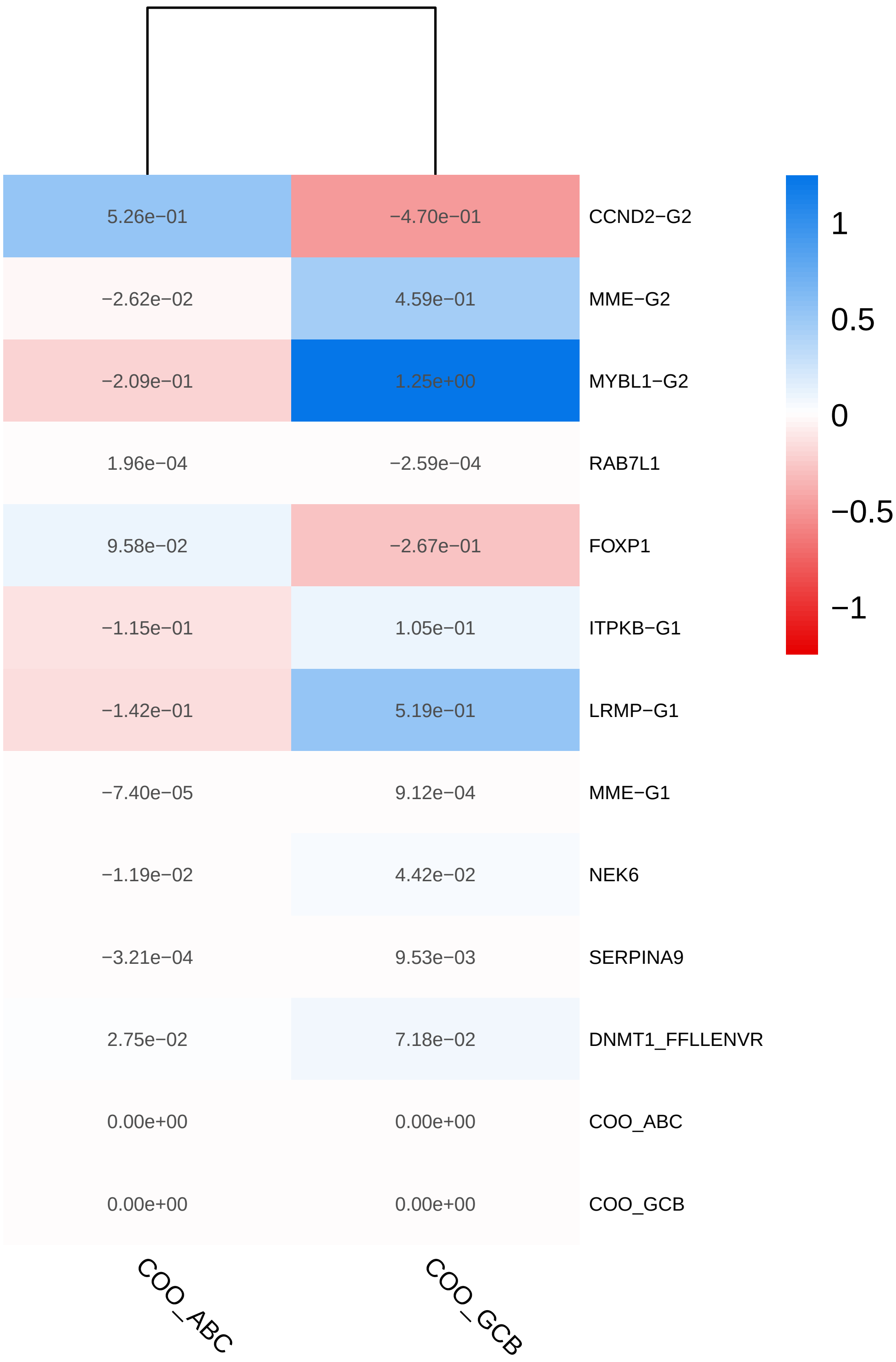


Figure S19: First order neighborhoods of the COO variables ABC (left) and GCB (right) with unclassified cases used as baseline, calculated with PriOmics *Prior B* based on the combined transcriptomics and proteomics dataset DS2. The values refer to the respective edge strengths. Gene names ending with "-G1" and "-G2" indicate the origin of the two transcriptomics datasets, i.e. measured either with NanoString nCounter (G1) or HTG (G2).

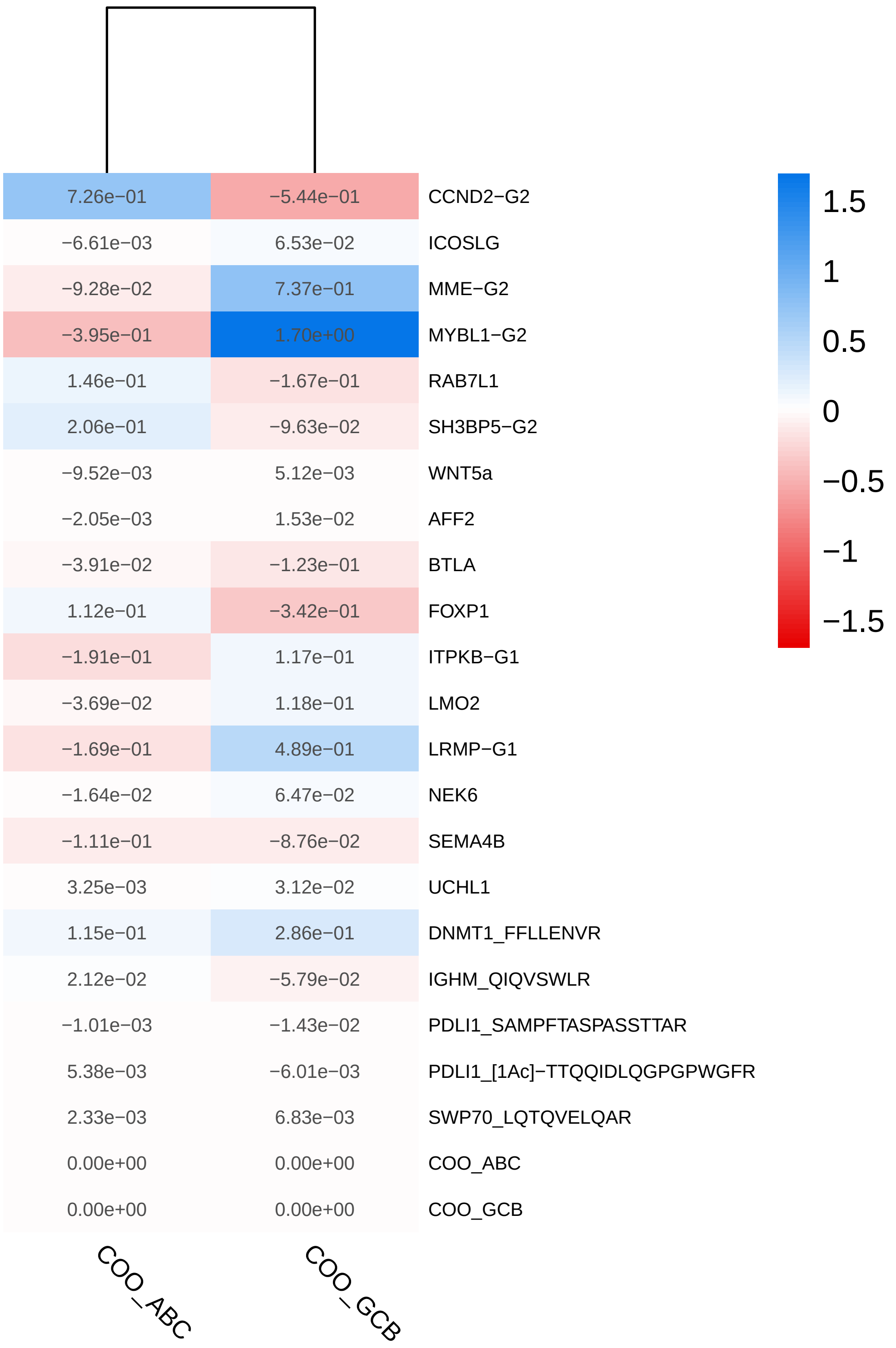


Figure S20: First order neighborhoods of the COO variables ABC (left) and GCB (right) with unclassified cases used as baseline, calculated with PLL-MGM based on the combined transcriptomics and proteomics dataset DS2. The values refer to the respective edge strengths. Gene names ending with "-G1" and "-G2" indicate the origin of the two transcriptomics datasets, i.e. measured either with NanoString nCounter (G1) or HTG (G2).

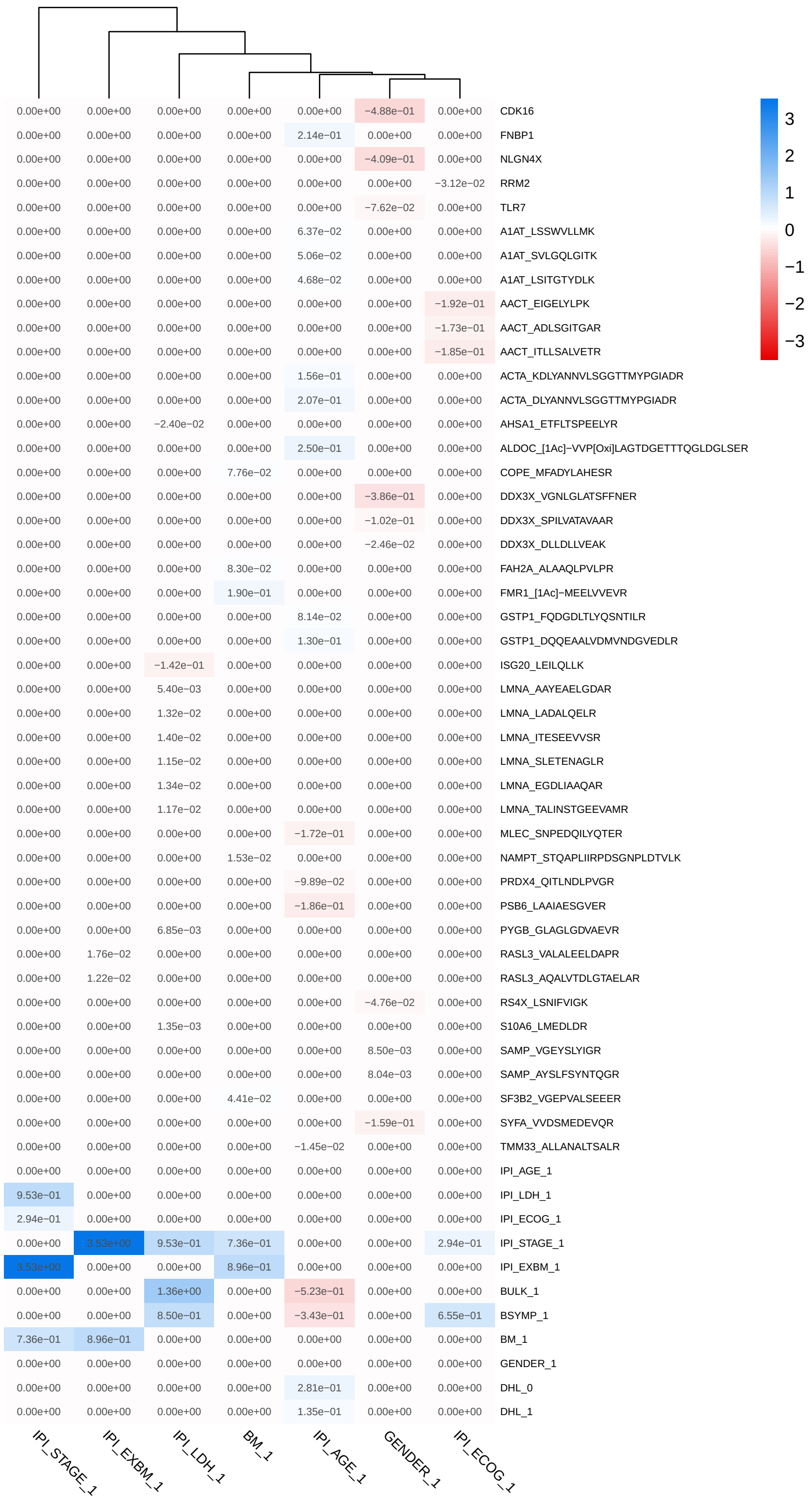


Figure S21: First order neighborhoods of the categorical variables calculated with PriOmics *Prior A* based on the combined transcriptomics and proteomics dataset DS2. The values refer to the respective edge strengths. The variable GENDER_1 represents the male patients.

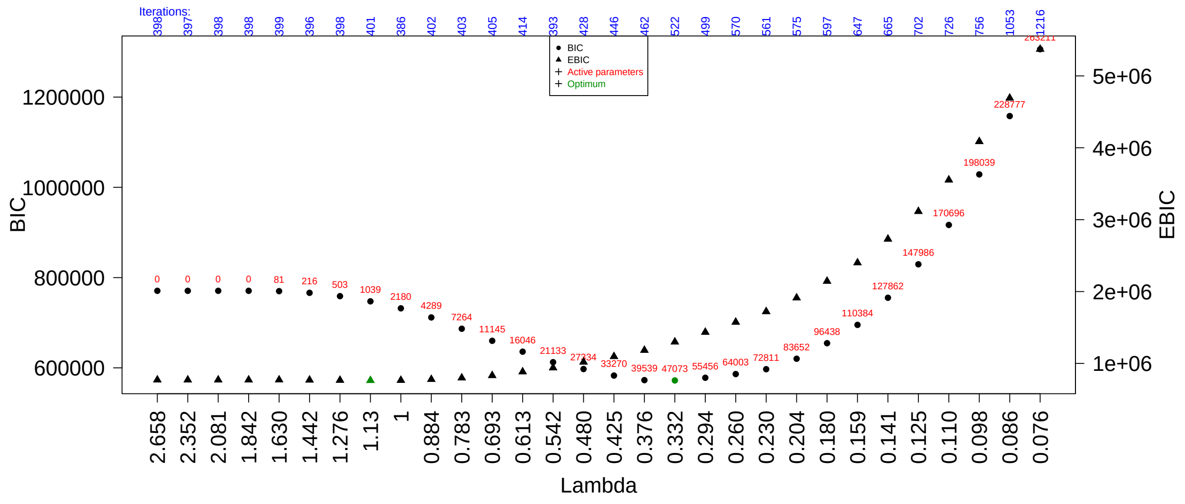

Figure S22: Model selection for dataset DS1: PriOmics - Prior A.

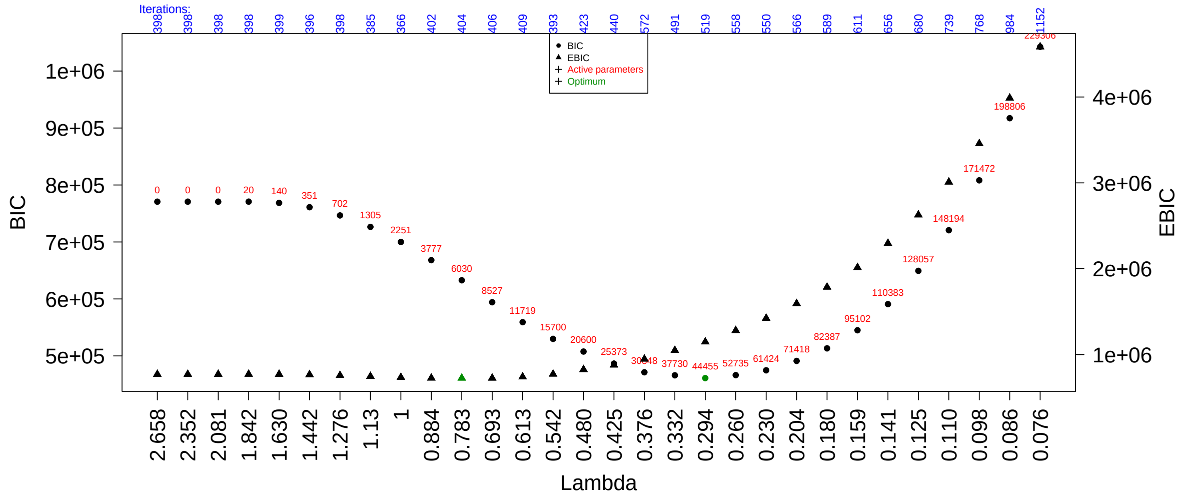

Figure S23: Model selection for dataset DS1: PriOmics - Prior B.

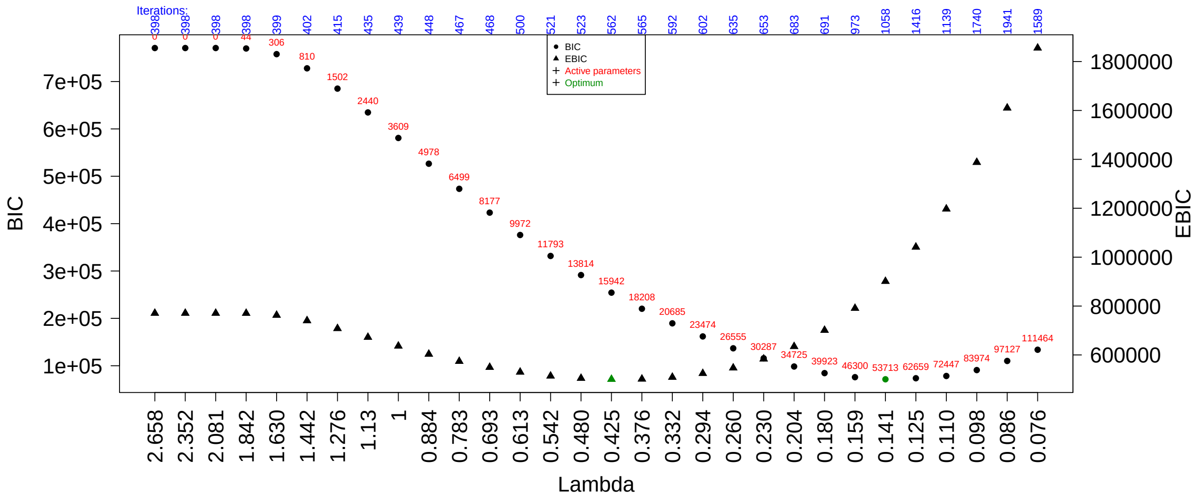


Figure S24: Model selection for dataset DS1: PLL-MGM.

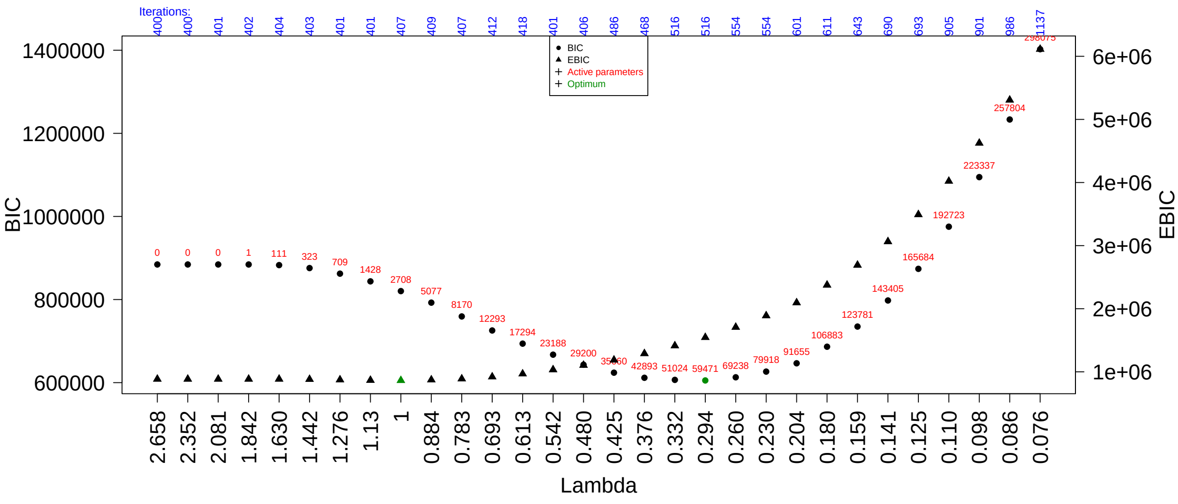


Figure S25: Model selection for dataset DS2: RNA - no prior; Proteomics - Prior A.

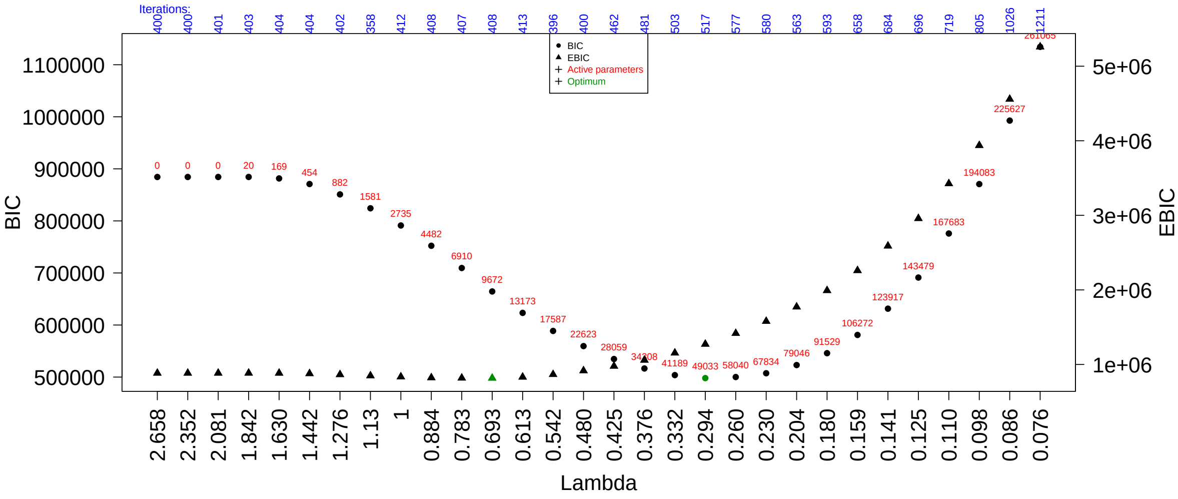


Figure S26: Model selection for dataset DS2: RNA - no prior; Proteomics - Prior B.

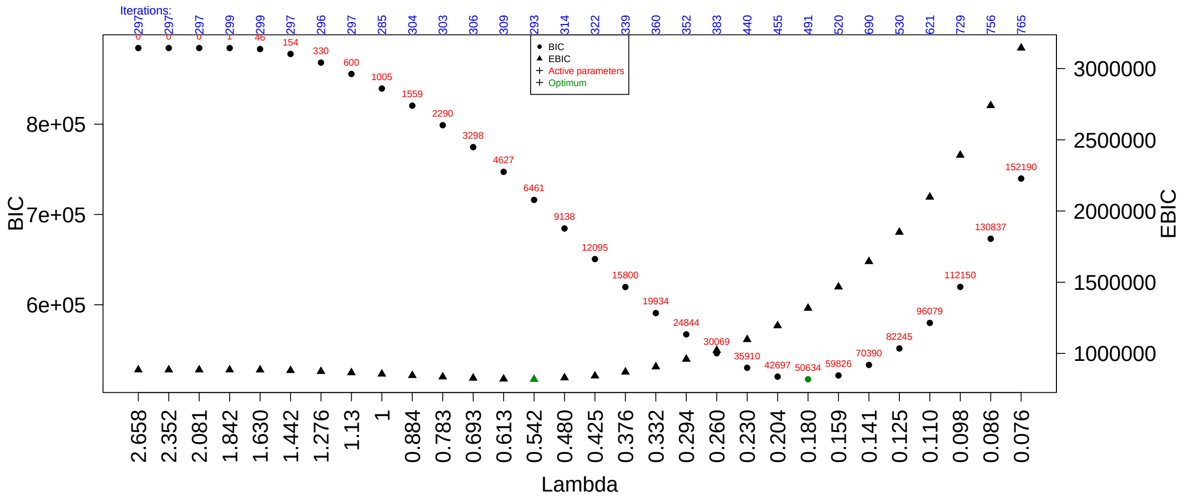


Figure S27: Model selection for dataset DS2: RNA - no prior; Proteomics - no prior; i.e. PLL-MGM.
