## Supplement peptide list for "PriOmics: integration of high-throughput proteomic data with complementary omics layers using mixed graphical modeling with group priors"

| Protein | Peptide | UniProt | Precursor_MZ | Precursor_Charge |
|---|---|---|---|---|
| 1433B | YLSEVASGDNK | P31946 | 591.79 | 2 |
| 1433E | YLAEFATGNDR | P62258 | 628.80 | 2 |
| 1433F | AVTELNEPLSNEDR | Q04917 | 793.89 | 2 |
| 1433G | YLAEVATGEK | P61981 | 540.78 | 2 |
| 1433G | NVTELNEPLSNEER | P61981 | 822.40 | 2 |
| 1433G | ATVVESSEK | P61981 | 475.25 | 2 |
| 1433T | AVTEQGAELSNEER | P27348 | 766.86 | 2 |
| 1433Z | TAFDEAIAELDTLSEESYK | P63104 | 711.34 | 3 |
| 1433Z | SVTEQGAELSNEER | P63104 | 774.86 | 2 |
| 1433Z | YLAEVAAGDDK | P63104 | 576.28 | 2 |
| 2AAA | LAGGDWFTSR | P30153 | 555.27 | 2 |
| 2AAA | MAGDPVANVR | P30153 | 515.26 | 2 |
| 2AAA | [1Ac]-AAADGDDSLYPIAVLIDELR | P30153 | 720.37 | 3 |
| 3BP1 | SLSSLDTALAELR | Q9Y3L3 | 688.38 | 2 |
| 4F2 | LLTSFLPAQLLR | P08195 | 686.42 | 2 |
| 6PGD | WTAISALEYGVPVTLIGEAVFAR | P52209 | 821.78 | 3 |
| 6PGD | HEMLPASLIQAQR | P52209 | 498.60 | 3 |
| 6PGD | AGQAVDDFIEK | P52209 | 596.80 | 2 |
| 6PGD | NPELQNLLLDDFFK | P52209 | 853.44 | 2 |
| 6PGL | ELPAAVAPAGPASLAR | O95336 | 745.92 | 2 |
| A16A1 | LLWTLESLVTGR | Q8IZ83 | 694.40 | 2 |
| A16A1 | AALLWALAAALER | Q8IZ83 | 684.90 | 2 |
| A1AT | LSSWVLLMK | P01009 | 538.81 | 2 |
| A1AT | SVLGQLGITK | P01009 | 508.31 | 2 |
| A1AT | LSITGTYDLK | P01009 | 555.81 | 2 |
| A2MG | VGFYESDVMGR | P01023 | 630.29 | 2 |
| A2MG | LLIYAVLPTGDVIGDSAK | P01023 | 923.02 | 2 |
| AACT | EIGELYLPK | P01011 | 531.30 | 2 |
| AACT | ADLSGITGAR | P01011 | 480.76 | 2 |
| AACT | ITLLSALVETR | P01011 | 608.37 | 2 |
| AATM | IAAAILNTPDLR | P00505 | 634.37 | 2 |
| AATM | FVTVQTISGTGALR | P00505 | 725.41 | 2 |
| AATM | IGASFLQR | P00505 | 446.26 | 2 |
| AATM | DAGMQLQGYR | P00505 | 569.77 | 2 |
| ABCD3 | VLGELWPLFGGR | P28288 | 672.38 | 2 |
| ABCE1 | NVEDLSGGELQR | P61221 | 658.83 | 2 |
| ABCF1 | MEETPTEYLQR | Q8NE71 | 698.82 | 2 |
| ABCF1 | LQGQLEQGDDTAAER | Q8NE71 | 815.89 | 2 |
| ABCF1 | STLLLLLTGK | Q8NE71 | 529.84 | 2 |
| ABHDA | VLSTDVDVILR | Q9NUJ1 | 615.36 | 2 |
| ABHEB | FSSETWQNLGTLHR | Q96IU4 | 559.28 | 3 |
| ABRAL | [1Ac]-MNVDHEVNLLVEEIHR | Q9P1F3 | 663.67 | 3 |
| ACADM | TRPVVAAGAVGLAQR | P11310 | 489.29 | 3 |
| ACADM | ANWYFLLAR | P11310 | 577.31 | 2 |
| ACADV | AGLGSGLSLSGLVHPELSR | P49748 | 617.34 | 3 |
| ACADV | FGMAAALAGTMR | P49748 | 598.80 | 2 |
| ACLY | TIAIIAEGIPEALTR | P53396 | 784.46 | 2 |
| ACLY | DLVSSLTSGLLTIGDR | P53396 | 823.95 | 2 |
| ACLY | [CRM]-AGK[1Ac]DLVSSLTSGLLTIGDR | P53396 | 663.36 | 3 |
| ACON | NAVTQEFGPVPDTAR | Q99798 | 801.40 | 2 |
| ACON | WVVIGDENYGEGSSR | Q99798 | 834.39 | 2 |
| ACON | SQFTITPGSEQIR | Q99798 | 732.38 | 2 |
| ACTA | KDLYANNVLSGGTTMYPGIADR | P62736 | 786.06 | 3 |
| ACTA | DLYANNVLSGGTTMYPGIADR | P62736 | 743.36 | 3 |
| ACTG | GYSFTTTAER | P63261 | 566.77 | 2 |
| ACTG | VAPEEHPVLLTEAPLNPK | P63261 | 652.03 | 3 |
| ACTG | TTGIVMDSGDGVTHTVPIYEGYALPHAILR | P63261 | 796.66 | 4 |
| ACTG | DLYANTVLSGGTTMYPGIADR | P63261 | 739.03 | 3 |
| ACTG | KDLYANTVLSGGTTMYPGIADR | P63261 | 781.73 | 3 |
| ACTN1 | LSNRPAFMPSEGR | P12814 | 487.91 | 3 |
| ACTN4 | NFITAEELR | O43707 | 546.79 | 2 |
| ACTN4 | GISQEQMQEFR | O43707 | 676.82 | 2 |
| ACTS | K[1Ac]DLYANNVM[CRM]SGGTTMYPGIADR | P68133 | 820.38 | 3 |
| ACTZ | TLFSNIVLSGGSTLFK | P61163 | 842.47 | 2 |
| ADRM1 | LFFWMQEPK | Q16186 | 613.31 | 2 |
| ADRM1 | SQSAAVTPSSTTSSTR | Q16186 | 784.38 | 2 |
| ADT2 | DFLAGGVAAAISK | P05141 | 610.34 | 2 |
| ADT2 | AAYFGIYDTAK | P05141 | 610.30 | 2 |
| ADT2 | [1Ac]-TDAAVSFAK | P05141 | 476.24 | 2 |
| ADT2 | [1Ac]-TDAAVSFAK[Frm]DFLAGGVAAAISK | P05141 | 727.38 | 3 |
| ADT2 | [1Ac]-TDAAVSFAK[AAR]DFLAGGVAAAISK | P05141 | 727.39 | 3 |
| ADT3 | DFLAGGIAAAISK | P12236 | 617.35 | 2 |

| | | | | |
|---|---|---|---|---|
| ADT3 | [1Ac]-TEQAISFAK | P12236 | 518.77 | 2 |
| AGM1 | MGLLAVLR | O95394 | 436.77 | 2 |
| AHSA1 | ETFLTSPEELYR | O95433 | 742.87 | 2 |
| AIFM1 | ALGTEVIQLFPEK | O95831 | 722.91 | 2 |
| AIP | VLELDPALAPVVSR | O00170 | 739.93 | 2 |
| AK1A1 | SPAQILLR | P14550 | 449.28 | 2 |
| AL1A1 | IFVEESIYDEFVR | P00352 | 823.41 | 2 |
| AL1A1 | SSSGT[Pho]PDLPVLLTDLK | P00352 | 574.96 | 3 |
| AL5AP | TISTTISPLLLIP | P20292 | 684.92 | 2 |
| AL9A1 | ANDTTFGLAAGVFTR | P49189 | 770.89 | 2 |
| ALBU | RHPDYSVVLLLR | P02768 | 489.95 | 3 |
| ALBU | VPQVSTPTLVEVSR | P02768 | 756.43 | 2 |
| ALBU | HPDYSVVLLLR | P02768 | 656.37 | 2 |
| ALBU | YLYEIAR | P02768 | 464.25 | 2 |
| ALBU | FQNALLVR | P02768 | 480.78 | 2 |
| ALBU | RY[Amn]TKK[1Ac]VPQVSTPTLVEVSR | P02768 | 562.07 | 4 |
| ALBU | [+1K]-YTK[1Ac]K[1Ac]VPQ[Dea]VSTPTLVEVSR | P02768 | 562.07 | 4 |
| ALBU | RHPYFYAPELLFFAK[1Ac]R[CRM] | P02768 | 535.78 | 4 |
| ALDH2 | AAFQLGSPWR | P05091 | 566.80 | 2 |
| ALDH2 | TFVQEDIYDEFVER | P05091 | 895.42 | 2 |
| ALDH2 | VIQVAAGSSNLK | P05091 | 593.84 | 2 |
| ALDH2 | VAFTGSTEIGR | P05091 | 569.30 | 2 |
| ALDOA | QLLLTADDR | P04075 | 522.79 | 2 |
| ALDOA | IGEHTPSALAIMENANVLAR | P04075 | 703.04 | 3 |
| ALDOA | RLQSIGTENTEENR | P04075 | 549.61 | 3 |
| ALDOA | FSHEEIAMATVTALR | P04075 | 559.29 | 3 |
| ALDOA | GILAADESTGSIAK | P04075 | 666.85 | 2 |
| ALDOC | [1Ac]-VVP[Oxi]LAGTDGETTTQGLDGLSER | P09972 | 758.71 | 3 |
| ALDR | TTAQVLIR | P15121 | 451.28 | 2 |
| ALDR | VAIDVGYR | P15121 | 446.75 | 2 |
| AMPB | GLSGTAVLDLR | Q9H4A4 | 551.32 | 2 |
| AMPL | TLIEFLLR | P28838 | 502.81 | 2 |
| AMPL | QLMETPANEMTPTR | P28838 | 809.88 | 2 |
| AMPL | GVLFASGQNLAR | P28838 | 616.84 | 2 |
| AMPL | LFEASIETGDR | P28838 | 619.31 | 2 |
| AMPL | ETLNISGPPLK | P28838 | 584.83 | 2 |
| AMPL | TK[1Ac]GLVLGIYSKEK[1Ac] | P28838 | 760.44 | 2 |
| AN32A | VSGGLEVLAEK | P39687 | 551.31 | 2 |
| AN32B | IFGGLDMLAEK | Q92688 | 597.32 | 2 |
| ANM1 | ATLYVTAIEDR | Q99873 | 626.33 | 2 |
| ANM5 | VPLVAPEDLR | O14744 | 554.82 | 2 |
| ANT3 | EVPLNTIIFMGR | P01008 | 695.38 | 2 |
| ANX11 | SETDLLDIR | P50995 | 531.28 | 2 |
| ANX11 | AHLVAVFNEYQR | P50995 | 482.92 | 3 |
| ANX11 | DAQELYAAGENR | P50995 | 668.81 | 2 |
| ANX11 | DESTNVDMSLAQR | P50995 | 733.33 | 2 |
| ANXA1 | GLGTDEDTLIEILASR | P04083 | 851.95 | 2 |
| ANXA1 | GVDEATIIDILTK | P04083 | 694.39 | 2 |
| ANXA1 | GTDVNVFNTILTTR | P04083 | 775.91 | 2 |
| ANXA1 | SEDFGVNEDLADSDAR | P04083 | 870.37 | 2 |
| ANXA1 | TPAQFDADELR | P04083 | 631.80 | 2 |
| ANXA1 | [1Ac]-AMVSEFLK | P04083 | 483.75 | 2 |
| ANXA2 | TNQELQEINR | P07355 | 622.82 | 2 |
| ANXA2 | QDIAFAYQR | P07355 | 556.28 | 2 |
| ANXA2 | SALSGHLETVILGLLK | P07355 | 551.00 | 3 |
| ANXA2 | GVDEVTIVNILTNR | P07355 | 771.93 | 2 |
| ANXA2 | DALNIETAIK | P07355 | 544.30 | 2 |
| ANXA2 | SYSPYDMLESIR | P07355 | 730.84 | 2 |
| ANXA3 | [1Ac]-ASIWVGHR | P12429 | 484.26 | 2 |
| ANXA4 | AEIDMLDIR | P09525 | 538.28 | 2 |
| ANXA4 | DEGNYLDDALVR | P09525 | 690.33 | 2 |
| ANXA4 | ISQTYQQQYGR | P09525 | 686.34 | 2 |
| ANXA5 | SEIDLFNIR | P08758 | 553.80 | 2 |
| ANXA5 | GTVTDFPGFDER | P08758 | 670.81 | 2 |
| ANXA5 | VLTEIIASR | P08758 | 501.30 | 2 |
| ANXA5 | GLGTDEESILTLLTSR | P08758 | 852.95 | 2 |
| ANXA5 | FITIFGTR | P08758 | 477.77 | 2 |
| ANXA5 | ETSGNLEQLLLAVVK | P08758 | 807.46 | 2 |
| ANXA6 | SEIDLLNIR | P08133 | 536.80 | 2 |
| ANXA6 | GLGTDEDTIIDIITHR | P08133 | 590.31 | 3 |
| ANXA6 | DAFVAIVQSVK | P08133 | 588.83 | 2 |
| ANXA6 | EAILDIITSR | P08133 | 565.82 | 2 |
| ANXA6 | EEGGENLDQAR | P08133 | 609.27 | 2 |
| AP1B1 | LAPPLVTLLSAEPELQYVALR | Q10567 | 765.11 | 3 |

| | | | | |
|---|---|---|---|---|
| AP1B1 | GLEISGTFTR | Q10567 | 540.79 | 2 |
| AP1G1 | VLAINILGR | O43747 | 484.82 | 2 |
| AP1M1 | VFLSGMPELR | Q9BXS5 | 574.81 | 2 |
| AP2A1 | DFLTPPLLSVR | O95782 | 629.36 | 2 |
| APEX1 | QGFGELLQAVPLADSFR | P27695 | 924.49 | 2 |
| APMAP | LLEYDTVTR | Q9HDC9 | 555.30 | 2 |
| APOA1 | QGLLPVLESFK | P02647 | 615.86 | 2 |
| APOA1 | LSPLGEEMR | P02647 | 516.26 | 2 |
| APOA1 | DYVSQFEGSALGK | P02647 | 700.84 | 2 |
| APOA1 | DLATVYVDVLK | P02647 | 618.35 | 2 |
| APOE | LGPLVEQGR | P02649 | 484.78 | 2 |
| APOE | LAVYQAGAR | P02649 | 474.77 | 2 |
| APOE | FWDYLR | P02649 | 450.22 | 2 |
| APOE | AATVGSLAGQPLQER | P02649 | 749.40 | 2 |
| APOL2 | ISAEGGEQVER | Q9BQE5 | 587.79 | 2 |
| APOL3 | ISAGSGGQAER | O95236 | 516.76 | 2 |
| APT | SFPDFPTPGVVFR | P07741 | 733.38 | 2 |
| APT | IDYIAGLDSR | P07741 | 561.79 | 2 |
| APT | [1Ac]-ADSELQLVEQR | P07741 | 665.34 | 2 |
| ARBK1 | SLLEGLLQR | P25098 | 514.81 | 2 |
| ARC1B | TWKPTLVILR | O15143 | 409.59 | 3 |
| ARC1B | NAYVWTLK | O15143 | 497.77 | 2 |
| ARF4 | IQEVADELQK | P18085 | 586.81 | 2 |
| ARF4 | DAVLLLFANK | P18085 | 552.33 | 2 |
| ARF5 | VQESADELQK | P84085 | 573.79 | 2 |
| ARF6 | ILMLGLDAAGK | P62330 | 551.32 | 2 |
| ARF6 | DAIILIFANK | P62330 | 559.33 | 2 |
| ARHG1 | QESGYLIEEIGDVLLAR | Q92888 | 635.67 | 3 |
| ARK72 | FYAYNPLAGGLLTGK | O43488 | 792.92 | 2 |
| ARK72 | VASVLGTMEMGR | O43488 | 625.82 | 2 |
| ARL8B | GVNAIVYMIDAADR | Q9NVJ2 | 754.38 | 2 |
| ARL8B | MNLSAIQDR | Q9NVJ2 | 524.27 | 2 |
| ARP2 | ILLTEPPMNPTK | P61160 | 677.38 | 2 |
| ARP2 | DLMVGDEASELR | P61160 | 667.82 | 2 |
| ARP2 | HIVLSGGSTMYPGLPSR | P61160 | 591.31 | 3 |
| ARP2 | HMVFLGGAVLADIMK | P61160 | 534.62 | 3 |
| ARP3 | DITYFIQQLLR | P61158 | 705.39 | 2 |
| ARP3 | NIVLSGGSTMFR | P61158 | 641.33 | 2 |
| ARP3 | AEPEDHYFLLTEPPLNTPENR | P61158 | 828.07 | 3 |
| ARP3 | LSEELSGGR | P61158 | 474.24 | 2 |
| ARP3 | FMEQVIFK | P61158 | 521.28 | 2 |
| ARP3 | HGIVEDWDLMER | P61158 | 750.35 | 2 |
| ARP5L | ALAVGGLGSIIR | Q9BPX5 | 563.85 | 2 |
| ARPC2 | ASHTAPQVLFSHR | O15144 | 484.26 | 3 |
| ARPC2 | DNTINLIHTFR | O15144 | 672.36 | 2 |
| ARPC2 | MILLEVNNR | O15144 | 551.31 | 2 |
| ARPC2 | DDETMYVESK | O15144 | 608.76 | 2 |
| ARPC2 | DTDAAVGDNIGYITFVLFPR | O15144 | 728.70 | 3 |
| ARPC3 | LIGNMALLPIR | O15145 | 605.87 | 2 |
| ARPC4 | AENFFILR | P59998 | 505.28 | 2 |
| ARPC4 | ELLLQPVTISR | P59998 | 634.88 | 2 |
| ARPC4 | VLIEGSINSVR | P59998 | 593.84 | 2 |
| ARPC5 | ALAAGGVGSIVR | O15511 | 535.82 | 2 |
| ASAH1 | WYVVQTNYDR | Q13510 | 672.32 | 2 |
| ASNA | LLNFPTIVER | O43681 | 601.35 | 2 |
| ASNA | LEETLPVIR | O43681 | 535.32 | 2 |
| ASNS | ELYLFDVLR | P08243 | 584.32 | 2 |
| AT1A1 | LNIPVSQVNPR | P05023 | 618.86 | 2 |
| AT1A1 | AVAGDASESALLK | P05023 | 616.33 | 2 |
| AT2A2 | EFDELNPSAQR | P16615 | 653.31 | 2 |
| AT2A2 | IGIFGQDEDVTSK | P16615 | 704.85 | 2 |
| AT2A3 | EFDDLSPEQQR | Q93084 | 682.31 | 2 |
| AT2A3 | [1Ac]-MEAAHLLPAADVLR | Q93084 | 774.91 | 2 |
| AT2B1 | GQILWFR | P20020 | 460.26 | 2 |
| AT5F1 | NNIAMALEVTYR | P24539 | 697.86 | 2 |
| ATD3A | GLLLFVDEADAFLR | Q9NVI7 | 789.93 | 2 |
| ATLA3 | SFILDFMLR | Q6DD88 | 571.31 | 2 |
| ATP5H | LAALPENPPAIDWAYYK | O75947 | 966.50 | 2 |
| ATP5I | YSALFLGVAYGATR | P56385 | 744.90 | 2 |
| ATPA | TGAIVDVPVGEELLGR | P25705 | 812.95 | 2 |
| ATPA | AVDSLVPIGR | P25705 | 513.80 | 2 |
| ATPA | VLSIGDGIAR | P25705 | 500.79 | 2 |
| ATPA | EIVTNFLAGFEA | P25705 | 655.83 | 2 |
| ATPA | LELAQYR | P25705 | 446.75 | 2 |

| | | | | |
|---|---|---|---|---|
| ATPA | ILGADTSVDLEETGR | P25705 | 788.40 | 2 |
| ATPB | VALTGLTVAEYFR | P06576 | 720.40 | 2 |
| ATPB | FTQAGSEVSALLGR | P06576 | 718.38 | 2 |
| ATPB | IMNVIGEPIDER | P06576 | 693.36 | 2 |
| ATPB | LVLEVAQHLGESTVR | P06576 | 550.98 | 3 |
| ATPB | IMDPNIVGSEHYDVAR | P06576 | 605.96 | 3 |
| ATPB | AIAELGIYPAVDPLDSTSR | P06576 | 994.52 | 2 |
| ATPD | AQAELVGTADEATR | P30049 | 716.36 | 2 |
| ATPK | LGELPSWILMR | P56134 | 657.87 | 2 |
| ATPK | DFSPSGIFGAFQR | P56134 | 714.85 | 2 |
| ATPO | TDPSILGGMIVR | P48047 | 629.84 | 2 |
| ATPO | YATALYSAASK | P48047 | 573.30 | 2 |
| ATPO | LVRPPVQVYGIEGR | P48047 | 528.31 | 3 |
| B3AT | VLLPLIFR | P02730 | 485.83 | 2 |
| B3AT | ADFLEQPVLGFVR | P02730 | 745.90 | 2 |
| BACH | LMDEVAGIVAAR | O00154 | 622.84 | 2 |
| BAF | AYVVLGQFLVLK | O75531 | 675.41 | 2 |
| BASI | FFVSSSQGR | P35613 | 507.75 | 2 |
| BAX | MIAAVDTDSPR | Q07812 | 588.29 | 2 |
| BAX | IGDELDSNMELQR | Q07812 | 760.36 | 2 |
| BCAP | VATEAEFSPEDSPSVR | Q6ZUJ8 | 860.90 | 2 |
| BGH3 | EGVYTVFAPTNEAFR | Q15582 | 850.92 | 2 |
| BGH3 | YGTLFTMDR | Q15582 | 552.26 | 2 |
| BGH3 | LTLLAPLNSVFK | Q15582 | 658.40 | 2 |
| BGH3 | VLTPPMGTVMDVLK | Q15582 | 750.91 | 2 |
| BGH3 | FSMLVAAIQSAGLTETLNR | Q15582 | 674.69 | 3 |
| BID | DLATALEQLLQAYPR | P55957 | 567.98 | 3 |
| BID | IEADSESQEDIIR | P55957 | 752.86 | 2 |
| BIEA | FGFPAFSGISR | P53004 | 593.31 | 2 |
| BIEA | NPHPSSAFLNLIGFVSR | P53004 | 619.33 | 3 |
| BLMH | IGPITPLEFYR | Q13867 | 653.36 | 2 |
| BLVRB | NDLSPTTVMSEGAR | P30043 | 739.35 | 2 |
| BLVRB | TVAGQDAVIVLLGTR | P30043 | 756.94 | 2 |
| BOLA2 | DLEAEHVEVEDTTLNR | Q9H3K6 | 623.97 | 3 |
| BOLA2 | [1Ac]-MELSAEYLR | Q9H3K6 | 577.28 | 2 |
| BTK | [1Ac]-AAVILESIFLK | Q06187 | 623.38 | 2 |
| BUB3 | VLVWDLR | O43684 | 450.77 | 2 |
| BUB3 | VYTLSVSGDR | O43684 | 548.79 | 2 |
| C1QC | FNAVLTNPQGDYDTSTGK | P02747 | 964.46 | 2 |
| C1TC | LAILQVGNR | P11586 | 492.30 | 2 |
| C1TC | LDIDPETITWQR | P11586 | 743.88 | 2 |
| C1TC | MFGIPVVVAVNAFK | P11586 | 746.42 | 2 |
| CAB39 | LLGELLLDR | Q9Y376 | 521.32 | 2 |
| CALD1 | NDDDEEEAAR | Q05682 | 582.23 | 2 |
| CALM | DTDSEEEIR | P62158 | 547.24 | 2 |
| CALM3 | [1Ac]-ADQLTEEQIAEFK | P0DP25 | 782.38 | 2 |
| CALM3 | HVMTNLGEK[1Ac]LTDEEVDEMIR | P0DP25 | 801.05 | 3 |
| CALM3 | HVMTNLGEK[1Ac]LTDEEVDEMIR | P0DP25 | 601.04 | 4 |
| CALR | FYALSASFEPFSNK | P27797 | 804.39 | 2 |
| CALR | GLQTSQDAR | P27797 | 488.25 | 2 |
| CALR | QIDNPDYK | P27797 | 496.74 | 2 |
| CALR | EQFLDGDGWTSR | P27797 | 705.82 | 2 |
| CALX | GTLSGWILSK | P27824 | 531.30 | 2 |
| CALX | APVPTGEVYFADSFDR | P27824 | 885.92 | 2 |
| CALX | AEEDEILNR | P27824 | 544.76 | 2 |
| CALX | KIPNPDFFEDLEPFR | P27824 | 621.98 | 3 |
| CAN1 | DFFLANASR | P07384 | 520.76 | 2 |
| CAN1 | LYELIITR | P07384 | 510.81 | 2 |
| CAND1 | IDLRPVLGEGVPILASFLR | Q86VP6 | 689.08 | 3 |
| CAND1 | ISGSILNELIGLVR | Q86VP6 | 742.45 | 2 |
| CAND1 | AVAALLTIPEAEK | Q86VP6 | 663.39 | 2 |
| CAND1 | LTLIDPETLLPR | Q86VP6 | 690.91 | 2 |
| CAND1 | LGTLSALDILIK | Q86VP6 | 628.89 | 2 |
| CAP1 | LSDLLAPISEQIK | Q01518 | 713.91 | 2 |
| CAP1 | LEAVSHTSDMHR | Q01518 | 461.55 | 3 |
| CAP1 | AGAAPYVQAFDSLLAGPVAEYLK | Q01518 | 784.41 | 3 |
| CAP1 | EMNDAAMFYTNR | Q01518 | 731.81 | 2 |
| CAPG | QAALQVAEGFISR | P40121 | 695.38 | 2 |
| CAPZB | STLNEIYFGK | P47756 | 586.30 | 2 |
| CAPZB | SGSGTMNLGGSLTR | P47756 | 669.33 | 2 |
| CASP3 | SGTDVDAANLR | P42574 | 559.78 | 2 |
| CATA | GPLLVQDVVFTDEMAHFDR | P04040 | 730.36 | 3 |
| CATA | AFYVNVLNEEQR | P04040 | 741.37 | 2 |
| CATA | LSQEDPDYGIR | P04040 | 646.81 | 2 |

| | | | | |
|---|---|---|---|---|
| CATA | FSTVAGESGSADTVR | P04040 | 742.35 | 2 |
| CATB | LPASFDAR | P07858 | 438.73 | 2 |
| CATC | NVHGINFVSPVR | P53634 | 446.91 | 3 |
| CATD | VGFAEAAR | P07339 | 410.72 | 2 |
| CATD | LVDQNIFSFYLSR | P07339 | 801.42 | 2 |
| CATD | VSTLPAITLK | P07339 | 521.83 | 2 |
| CATD | FDGILGMAYPR | P07339 | 620.31 | 2 |
| CATD | ISVNNVLPVFDNLMQQK | P07339 | 980.02 | 2 |
| CATG | VSSFLPWIR | P08311 | 552.81 | 2 |
| CATZ | VGDYGSLSGR | Q9UBR2 | 505.75 | 2 |
| CATZ | NVDGVNYASITR | Q9UBR2 | 654.83 | 2 |
| CAZA1 | FITHAPPGEFNEVFNDVR | P52907 | 697.01 | 3 |
| CBR1 | VVNVSSIMSVR | P16152 | 595.83 | 2 |
| CBR1 | LFSGDVVLTAR | P16152 | 589.33 | 2 |
| CBR1 | FHQLDIDDLQSIR | P16152 | 533.94 | 3 |
| CBR1 | GQAAVQQLQAEGLSPR | P16152 | 826.94 | 2 |
| CBR1 | [1Ac]-SSGIHVALVTGGNK | P16152 | 691.38 | 2 |
| CBX3 | IIGATDSSGELMFLMK | Q13185 | 856.93 | 2 |
| CC124 | VLEEGSVEAR | Q96CT7 | 544.78 | 2 |
| CCAR2 | VLLLSSPGLEELYR | Q8N163 | 794.95 | 2 |
| CCAR2 | INPLPGGR | Q8N163 | 412.24 | 2 |
| CCAR2 | LAEAEETAR | Q8N163 | 495.25 | 2 |
| CD14 | LTVGAAQVPAQLLVGALR | P08571 | 889.04 | 2 |
| CD20 | MESLNFIR | P11836 | 505.26 | 2 |
| CD44 | YGFIEGHVVIPR | P16070 | 462.92 | 3 |
| CD99 | NANAEPAVQR | P14209 | 535.27 | 2 |
| CDC42 | TPFLLVGTQIDLR | P60953 | 736.93 | 2 |
| CDIPT | FGAMLDMLTDR | O14735 | 635.30 | 2 |
| CDK1 | LESEEEGVPSTAIR | P06493 | 758.88 | 2 |
| CECR5 | IEGVLLLGEPVR | Q9BXW7 | 647.89 | 2 |
| CERU | GPEEEHLGILGPVIWAEVGDTIR | P00450 | 829.77 | 3 |
| CFAB | DFHINLFQVLPWLK | P00751 | 590.66 | 3 |
| CH10 | FLPLFDR | P61604 | 454.26 | 2 |
| CH10 | VLQATVVAVGSGSK | P61604 | 658.38 | 2 |
| CH10 | VVLDDKDYFLFR | P61604 | 510.60 | 3 |
| CH60 | VTDALNATR | P10809 | 480.76 | 2 |
| CH60 | ALMLQGVDLLADAVAVTMGPK | P10809 | 705.05 | 3 |
| CH60 | GVMLAVDAVIAELK | P10809 | 714.91 | 2 |
| CH60 | VGLQVVAVK | P10809 | 456.80 | 2 |
| CH60 | ISSIQSIVPALEIANAHR | P10809 | 640.36 | 3 |
| CH60 | ALMLQGVDLLADAVAVTMGPK[1Ac]GRT-[Ami] | P10809 | 823.45 | 3 |
| CH60 | ALMLQGVDLLADAVAVTMGPK[1Ac]GRT-[Ami] | P10809 | 617.84 | 4 |
| CHD4 | IGVMSLIR | Q14839 | 444.77 | 2 |
| CHTOP | ASMQQQQQLASAR | Q9Y3Y2 | 723.86 | 2 |
| CIRBP | GFGFVTFENIDDAK | Q14011 | 780.37 | 2 |
| CISY | ALGVLAQLIWSR | O75390 | 663.90 | 2 |
| CISY | GLVYETSVLDPDEGIR | O75390 | 881.95 | 2 |
| CISY | DYIWNTLNSGR | O75390 | 669.83 | 2 |
| CISY | DILADLIPK | O75390 | 499.30 | 2 |
| CK098 | VVGAVIDQGLITR | E9PRG8 | 670.90 | 2 |
| CKLF6 | SGLAAYFFMGR | Q9NX76 | 610.30 | 2 |
| CLH1 | LLLPWLEAR | Q00610 | 555.84 | 2 |
| CLH1 | ISGETIFVTAPHEATAGIIGVNR | Q00610 | 785.09 | 3 |
| CLH1 | VVGAMQLYSVDR | Q00610 | 669.35 | 2 |
| CLH1 | NNLAGAEELFAR | Q00610 | 652.83 | 2 |
| CLH1 | AFMTADLPNELIELLEK | Q00610 | 974.01 | 2 |
| CLH1 | NLQNLLILTAIK | Q00610 | 677.43 | 2 |
| CLIC1 | GVTFNVTTVDTK | O00299 | 641.34 | 2 |
| CLIC1 | LFMVLWLK | O00299 | 525.31 | 2 |
| CLUS | ASSIIDELFQDR | P10909 | 697.35 | 2 |
| CLUS | ELDESLQVAER | P10909 | 644.82 | 2 |
| CLUS | IDSLLENDR | P10909 | 537.77 | 2 |
| CLUS | VTTVASHTSDSDVPSGVTEVVVK | P10909 | 772.06 | 3 |
| CN37 | GEEVGELSR | P09543 | 488.24 | 2 |
| CNDP2 | TVFGVEPDLTR | Q96KP4 | 617.33 | 2 |
| CNDP2 | GNILIPGINEAVAAVTEEEHK | Q96KP4 | 735.39 | 3 |
| CNDP2 | TGQEIPVNVR | Q96KP4 | 556.81 | 2 |
| CNDP2 | [1Ac]-AALTTLFK | Q96KP4 | 453.77 | 2 |
| CO1A1 | GSEGPQGVR | P02452 | 443.72 | 2 |
| CO1A1 | GESGPSGPAGPTGAR | P02452 | 649.31 | 2 |
| CO1A1 | DGEAGAQGPPGPAGPAGER | P02452 | 845.89 | 2 |
| CO1A1 | [1Ac]-FS[Pho]GLQGPPGPPGSPGEQGPSGASGPAGPR | P02452 | 908.08 | 3 |
| CO1A1 | [1Ac]-GFS[Pho]GLQGPPGPPGS[Dhy]PGE[Dhy]QGPSGASGPAGPR | P02452 | 686.56 | 4 |
| CO1A1 | GFS[Pho]GLQGPPGPPGS[Dhy]P[Oxi]GEQGPSGASGPAGPR | P02452 | 912.41 | 3 |

| | | | | |
|---|---|---|---|---|
| CO1A1 | GFS[Pho]GLQGPPGP[Oxi]PGSPGE[Dhy]Q[Dea]GPSGASGPAGPR | P02452 | 912.74 | 3 |
| CO1A2 | GETGPSGPVGPAGAVGPR | P08123 | 781.90 | 2 |
| CO1A2 | GEAGAAGPAGPAGPR | P08123 | 618.31 | 2 |
| CO1A2 | GVVGPQGAR | P08123 | 420.74 | 2 |
| CO1A2 | GPSGPQGIR | P08123 | 434.74 | 2 |
| CO3 | TELRPGETLNVNFLLR | P01024 | 624.68 | 3 |
| CO3 | LVAYYTLIGASGQR | P01024 | 756.41 | 2 |
| CO3 | ILLQGTPVAQMTEDAVDAER | P01024 | 719.70 | 3 |
| CO3 | IPIEDGSGEVVLSR | P01024 | 735.89 | 2 |
| CO3 | NEQVEIR | P01024 | 444.23 | 2 |
| CO3 | SNLDEDIIAEENIVSR | P01024 | 908.95 | 2 |
| CO4B | VGDTLNLNLR | P0C0L5 | 557.81 | 2 |
| CO4B | TTNIQGINLLFSSR | P0C0L5 | 782.43 | 2 |
| CO6A1 | VPSYQALLR | P12109 | 523.80 | 2 |
| CO6A1 | VAVVQYSGTGQQRPER | P12109 | 592.31 | 3 |
| CO6A1 | TAEYDVAYGESHLFR | P12109 | 586.61 | 3 |
| CO6A1 | NLVWNAGALHYSDEVEIIQGLTR | P12109 | 866.78 | 3 |
| CO6A1 | IALVITDGR | P12109 | 479.29 | 2 |
| CO6A1 | VFSVAITPDHLEPR | P12109 | 527.62 | 3 |
| CO6A1 | [CRM]-VLVTGK[1Ac]TAEYDVAYGESHLFR | P12109 | 610.81 | 4 |
| CO6A1 | [CRM]-PGLSLVK[1Ac]ENYAELLEDAFLK | P12109 | 778.75 | 3 |
| CO6A2 | VFAVVITDGR | P12110 | 538.81 | 2 |
| CO6A2 | NFVINVVNR | P12110 | 537.81 | 2 |
| CO6A2 | DDDLNLR | P12110 | 430.71 | 2 |
| CO6A2 | DIASTPHELYR | P12110 | 651.33 | 2 |
| CO6A2 | YGGLHFSDQVEVFSPPGSDR | P12110 | 732.01 | 3 |
| CO6A3 | AAPLQGMLPGLLAPLR | P12111 | 809.48 | 2 |
| CO6A3 | ALILVGLER | P12111 | 492.32 | 2 |
| CO6A3 | SQAPVLDAIR | P12111 | 535.30 | 2 |
| CO6A3 | LSDAGITPLFLTR | P12111 | 702.40 | 2 |
| CO6A3 | VVESLDVGQDR | P12111 | 608.81 | 2 |
| COASY | SGLLVLTTPLASLAPR | Q13057 | 804.99 | 2 |
| COEA1 | NLVVGDETTSSLR | Q05707 | 695.86 | 2 |
| COF1 | LGGSAVISLEGKPL | P23528 | 670.89 | 2 |
| COF1 | YALYDATYETK | P23528 | 669.32 | 2 |
| COF1 | [1Ac]-ASGVAVSDGVIK | P23528 | 572.81 | 2 |
| COIA1 | LQDLYSIVR | P39060 | 553.81 | 2 |
| COIA1 | TEAPSATGQASSLLGGR | P39060 | 801.91 | 2 |
| COPA | TLDLPIYVTR | P53621 | 595.84 | 2 |
| COPA | VLTIDPTEFK | P53621 | 581.82 | 2 |
| COPB | VLQDLVMDILR | P53618 | 657.88 | 2 |
| COPB | VLSTPDLEVR | P53618 | 564.82 | 2 |
| COPB | NAVLAIYTIYR | P53618 | 648.87 | 2 |
| COPB | EAELLEPLMPAIR | P53618 | 741.41 | 2 |
| COPB2 | LPEAAFLAR | P35606 | 494.28 | 2 |
| COPB2 | LESTLNYGMER | P35606 | 656.81 | 2 |
| COPD | ENVNLAQIR | P48444 | 528.79 | 2 |
| COPD | IEGLLAAFPK | P48444 | 529.82 | 2 |
| COPD | NSNILEDLETLR | P48444 | 708.87 | 2 |
| COPE | MFADYLAHESR | O14579 | 447.21 | 3 |
| COPG1 | SLEELPVDIILASVG | Q9Y678 | 777.93 | 2 |
| COPG1 | VFNETPINPR | Q9Y678 | 593.81 | 2 |
| COPZ1 | [1Ac]-MEALILEPSLYTVK | P61923 | 824.95 | 2 |
| COR1A | DAGPLLISLK | P31146 | 513.81 | 2 |
| COR1A | YFEITSEAPFLHYLSMFSSK | P31146 | 799.72 | 3 |
| COR1A | AAPEASGTPSSDAVSR | P31146 | 751.86 | 2 |
| COR1A | HVFGQPAK | P31146 | 442.24 | 2 |
| COR1A | LQATVQELQK | P31146 | 579.33 | 2 |
| COR1B | DADPILISLR | Q9BR76 | 556.82 | 2 |
| CORO7 | LAVAGEDAR | P57737 | 451.24 | 2 |
| COTL1 | AAYNLVR | Q14019 | 403.73 | 2 |
| COTL1 | FALITWIGENVSGLQR | Q14019 | 902.49 | 2 |
| COTL1 | FTTGDAMSK | Q14019 | 479.22 | 2 |
| COTL1 | DDGSAVIWVTFK | Q14019 | 669.34 | 2 |
| COTL1 | [CRM]-SK[1Ac]FALITWIGENVSGLQR | Q14019 | 702.04 | 3 |
| COX2 | MMITSQDVLHSWAVPTLGLK | P00403 | 743.06 | 3 |
| COX2 | ILYMTDEVNDPSLTIK | P00403 | 926.47 | 2 |
| COX2 | IFEMGPVFTL | P00403 | 577.30 | 2 |
| COX41 | DHPLPEVAHVK | P13073 | 414.56 | 3 |
| COX41 | SEDFSLPAYMDR | P13073 | 715.82 | 2 |
| COX5A | LNDFASTVR | P20674 | 511.77 | 2 |
| CPNE1 | LYGPTNFAPIINHVAR | Q99829 | 594.99 | 3 |
| CPNE1 | DIVQFVPYR | Q99829 | 568.81 | 2 |
| CPNE1 | EALAQTVLAEVPTQLVSYFR | Q99829 | 745.74 | 3 |

| | | | | |
|---|---|---|---|---|
| CPNE1 | SDPFLEFFR | Q99829 | 579.28 | 2 |
| CPNE3 | DIVQFVPFR | O75131 | 560.81 | 2 |
| CPNE3 | VVLFEMEAR | O75131 | 547.29 | 2 |
| CPNS1 | SMVAVMDSDTTGK | P04632 | 671.30 | 2 |
| CPNS1 | THYSNIEANESEEVR | P04632 | 593.27 | 3 |
| CPNS1 | [1Ac]-MFLVNSFLK | P04632 | 570.81 | 2 |
| CPSF3 | GLIPVFALGR | Q9UKF6 | 521.82 | 2 |
| CPSF5 | TINLYPLTNYTFGTK | O43809 | 873.46 | 2 |
| CPSF6 | AVSDASAGDYGSAIETLVTAISLIK | Q16630 | 818.10 | 3 |
| CS043 | EAPGPAGGGGGGSR | Q9BQ61 | 563.77 | 2 |
| CSK | LLYPPETGLFLVR | P41240 | 759.44 | 2 |
| CSK | NVLVSEDNVAK | P41240 | 594.32 | 2 |
| CSK | GEFGDVMLGDYR | P41240 | 679.81 | 2 |
| CSK | VMEGTVAAQDEFYR | P41240 | 808.37 | 2 |
| CSN3 | [1Ac]-ASALEQFVNSVR | Q9UNS2 | 681.85 | 2 |
| CTBL1 | FVDILGLR | Q8WYA6 | 466.78 | 2 |
| CTBP1 | VGQAVALR | Q13363 | 407.25 | 2 |
| CUL2 | LMVEPLQAILIR | Q13617 | 698.42 | 2 |
| CY1 | GLLSSLDHTSIR | P08574 | 433.57 | 3 |
| CY24B | LLGSALALAR | P04839 | 492.81 | 2 |
| CYBP | SFDLLVK | Q9HB71 | 411.24 | 2 |
| CYC | ADLIAYLK | P99999 | 453.77 | 2 |
| CYC | MIFVGIK | P99999 | 404.24 | 2 |
| CYFP1 | NVIQSVLQAIR | Q7L576 | 620.87 | 2 |
| CYFP2 | NVLISVLQAIR | Q96F07 | 613.39 | 2 |
| CYTB | SQVVAGTNYFIK | P04080 | 663.86 | 2 |
| CYTB | VHVGDEDFVHLR | P04080 | 474.91 | 3 |
| DAD1 | FLEEYLSSTPQR | P61803 | 735.37 | 2 |
| DAD1 | [1Ac]-SASVVSVISR | P61803 | 523.80 | 2 |
| DBNL | NGPALQEAYVR | Q9UJU6 | 609.32 | 2 |
| DBNL | TWEQQQEVVSR | Q9UJU6 | 695.34 | 2 |
| DBNL | FQDVGPQAPVGSVYQK | Q9UJU6 | 860.44 | 2 |
| DBNL | AMSTTSISSPQPGK | Q9UJU6 | 696.35 | 2 |
| DC1I2 | ADAEEEAATR | Q13409 | 531.74 | 2 |
| DCPS | DLTPEHLPLLR | Q96C86 | 435.25 | 3 |
| DCTN2 | ENLATVEGNFASIDER | Q13561 | 882.92 | 2 |
| DCXR | AVIQVSQIVAR | Q7Z4W1 | 592.36 | 2 |
| DCXR | [1Ac]-MELFLAGR | Q7Z4W1 | 489.76 | 2 |
| DDB1 | DLLFILTAK | Q16531 | 517.32 | 2 |
| DDX1 | ELLIIGGVAAR | Q92499 | 556.35 | 2 |
| DDX17 | APLPDLYPFGTMR | Q92841 | 739.38 | 2 |
| DDX17 | VLEEANQAINPK | Q92841 | 663.36 | 2 |
| DDX17 | SSQSSSQQFSGIGR | Q92841 | 728.34 | 2 |
| DDX17 | DMVGIAQTGSGK | Q92841 | 582.29 | 2 |
| DDX21 | APQVLVLAPTR | Q9NR30 | 582.86 | 2 |
| DDX21 | GVTFLFPIQAK | Q9NR30 | 610.86 | 2 |
| DDX21 | WQLSVATEQPELEGPR | Q9NR30 | 920.47 | 2 |
| DDX27 | EAIVAALLTR | Q96GQ7 | 528.82 | 2 |
| DDX3X | VGNLGLATSFFNER | O00571 | 762.89 | 2 |
| DDX3X | SPILVATAVAAR | O00571 | 584.86 | 2 |
| DDX3X | DLLDLLVEAK | O00571 | 564.83 | 2 |
| DDX46 | VTYVVLDEADR | Q7L014 | 640.33 | 2 |
| DDX5 | TTYLVLDEADR | P17844 | 648.33 | 2 |
| DDX5 | LMEEIMSEK | P17844 | 555.26 | 2 |
| DDX5 | TAQEVETYR | P17844 | 548.77 | 2 |
| DDX6 | SGAYLIPLLER | P26196 | 616.36 | 2 |
| DDX6 | LDDTVHVVIATPGR | P26196 | 498.28 | 3 |
| DECR | FNVIQPGPIK | Q16698 | 556.83 | 2 |
| DEK | LTMQVSSLQR | P35659 | 581.82 | 2 |
| DEK | SGVNSELVK | P35659 | 466.76 | 2 |
| DEOC | IGASTLLSDIER | Q9Y315 | 637.85 | 2 |
| DERM | YFESVLDR | Q07507 | 514.76 | 2 |
| DHB12 | ISYSLFTALR | Q53GQ0 | 585.83 | 2 |
| DHB12 | TIAVDFASEDIYDK | Q53GQ0 | 793.88 | 2 |
| DHB4 | VVLVTGAGAGLGR | P51659 | 585.35 | 2 |
| DHB4 | AYALAFAER | P51659 | 506.27 | 2 |
| DHB4 | IDVVVNNAGILR | P51659 | 641.88 | 2 |
| DHB4 | VLQQFADNDVSR | P51659 | 696.35 | 2 |
| DHB4 | GALVVVNDLGGDFK | P51659 | 702.38 | 2 |
| DHE3 | TAAYVNAIEK | P00367 | 540.29 | 2 |
| DHE3 | YSTDVSVDEVK | P00367 | 621.30 | 2 |
| DHE3 | DSNYHLLMSVQESLER | P00367 | 640.98 | 3 |
| DHE3 | DDGSWEVIEGYR | P00367 | 713.32 | 2 |
| DHX15 | TLATDILMGVLK | O43143 | 637.87 | 2 |

| | | | | |
|---|---|---|---|---|
| DHX15 | FTDILVR | O43143 | 432.25 | 2 |
| DHX15 | EVDDLGPEVGDIK | O43143 | 693.34 | 2 |
| DHX15 | LQLPVWEYK | O43143 | 588.33 | 2 |
| DHX15 | SNLGSVVLQLK | O43143 | 579.35 | 2 |
| DHX9 | AIEPPPLDAVIEAEHTLR | Q08211 | 657.69 | 3 |
| DHX9 | DVVQAYPEVR | Q08211 | 588.31 | 2 |
| DHX9 | DFVNYLVR | Q08211 | 513.27 | 2 |
| DHX9 | GMTLVTPLQLLLFASK | Q08211 | 866.51 | 2 |
| DJB11 | TLEVEIEPGVR | Q9UBS4 | 621.34 | 2 |
| DKC1 | IMLPGVLR | O60832 | 449.78 | 2 |
| DLDH | NLGLEELGIELDPR | P09622 | 784.42 | 2 |
| DMA | YTAIAYWVPR | P28067 | 620.33 | 2 |
| DNJC7 | EALGDAQQSVR | Q99615 | 587.30 | 2 |
| DNM1L | [1Ac]-MEALIPVINK | O00429 | 585.33 | 2 |
| DNMT1 | FFLLENVR | P26358 | 519.29 | 2 |
| DNPEP | YASNAVSEALIR | Q9ULA0 | 647.34 | 2 |
| DNPH1 | FGTVLTEHVAAAELGAR | O43598 | 581.31 | 3 |
| DOCK2 | SSSSVGGLSVSSR | Q92608 | 605.31 | 2 |
| DOCK2 | RPFGVAVMDITDIIK | Q92608 | 558.98 | 3 |
| DOCK8 | GFVFNLIR | Q8NF50 | 483.28 | 2 |
| DPM3 | ELQSQIQEAR | Q9P2X0 | 601.31 | 2 |
| DPYL2 | ILDLGITGPEGHVLSRPEEVEAEAVNR | Q16555 | 725.88 | 4 |
| DPYL2 | MVIPGGIDVHTR | Q16555 | 432.24 | 3 |
| DPYL2 | SSAEVIAQAR | Q16555 | 516.28 | 2 |
| DQB1 | VEPTVTISPSR | P01920 | 593.33 | 2 |
| DRA | FASFEAQGALANIAVDK | P01903 | 876.45 | 2 |
| DRA | NGKPVTTGVSETVFLPR | P01903 | 601.33 | 3 |
| DRA | [AAR]-NGK[1Ac]P[Oxi]VTTGVSETVFLPR | P01903 | 630.01 | 3 |
| DRG1 | IQLLDLPGIIEGAK | Q9Y295 | 740.44 | 2 |
| DSRAD | MGFTEVTPVTGASLR | P55265 | 783.40 | 2 |
| DX39A | VNIVFNYDMPEDSDTYLHR | O00148 | 776.69 | 3 |
| DX39A | VSVFFGGLSIK | O00148 | 577.33 | 2 |
| DX39A | FEVNVAELPEEIDISTYIEQSR | O00148 | 861.09 | 3 |
| DX39B | VNIAFNYDMPEDSDTYLHR | Q13838 | 767.35 | 3 |
| DX39B | VAVFFGGLSIK | Q13838 | 569.34 | 2 |
| DX39B | FEVNISELPDEIDISSYIEQTR | Q13838 | 866.43 | 3 |
| DYHC1 | VTFVNFTVTR | Q14204 | 592.33 | 2 |
| DYHC1 | GIFEALRPLETLPVEGLIR | Q14204 | 708.41 | 3 |
| DYHC1 | IQGLTVEQAEAVVR | Q14204 | 756.92 | 2 |
| DYHC1 | TFSSIPVSR | Q14204 | 497.27 | 2 |
| DYHC1 | GNEIVLSAGSTPR | Q14204 | 650.85 | 2 |
| DYN2 | AIPNQGEILVIR | P50570 | 661.89 | 2 |
| E2AK2 | [1Ac]-AGDLSAGFFMEELNTYR | P19525 | 981.95 | 2 |
| ECH1 | YQETFNVIER | Q13011 | 649.82 | 2 |
| ECH1 | MMADEALGSGLVSR | Q13011 | 718.85 | 2 |
| ECHA | TVLGTPEVLLGALPGAGGTQR | P40939 | 669.71 | 3 |
| ECHA | FVDLYGAQK | P40939 | 520.77 | 2 |
| ECHA | ILQEGVDPK | P40939 | 499.78 | 2 |
| ECHB | LVMAAANR | P55084 | 423.24 | 2 |
| ECHB | FNFLAPELPAVSEFSTSETMGHSADR | P55084 | 947.44 | 3 |
| ECHB | AALTGLLHR | P55084 | 476.29 | 2 |
| ECHM | SLAMEMVLTGDR | P30084 | 661.83 | 2 |
| ECM29 | TEALSVIELLLK | Q5VYK3 | 664.91 | 2 |
| EDF1 | [1Ac]-AESDWDTVTVLR | O60869 | 717.35 | 2 |
| EF1A1 | VETGVLKPGMVVTFAPVNVTTEVK | P68104 | 839.13 | 3 |
| EF1A1 | YYVTIIDAPGHR | P68104 | 468.91 | 3 |
| EF1A1 | MDSTEPPYSQK | P68104 | 641.78 | 2 |
| EF1D | IASLEVENQSLR | P29692 | 679.87 | 2 |
| EF1D | GVVQELQQAISK | P29692 | 650.37 | 2 |
| EF1D | ATAPQTQHVSPMR | P29692 | 475.24 | 3 |
| EF1D | SLAGSSGPGASSGTSGDHGELVVR | P29692 | 729.02 | 3 |
| EF1D | [1Ac]-ATNFLAHEK | P29692 | 536.77 | 2 |
| EF1G | ILGLLDAYLK | P26641 | 559.84 | 2 |
| EF1G | ALIAAQYSGAQVR | P26641 | 674.37 | 2 |
| EF1G | VLSAPPHFHFGQTNR | P26641 | 427.72 | 4 |
| EF1G | LDPGSEETQTLVR | P26641 | 722.87 | 2 |
| EF1G | [1Ac]-AAGTLYTYPENWR | P26641 | 792.38 | 2 |
| EF2 | TILMMGR | P13639 | 411.22 | 2 |
| EF2 | FSVSPVVR | P13639 | 445.76 | 2 |
| EF2 | AYLPVNESFGFTADLR | P13639 | 900.45 | 2 |
| EFHD2 | LSEIDVSSEGVK | Q96C19 | 631.83 | 2 |
| EFHD2 | ADLNQGIGEPQSPSR | Q96C19 | 784.89 | 2 |
| EFHD2 | DGFIDLMELK | Q96C19 | 590.80 | 2 |
| EFTU | AEAGDNLGALVR | P49411 | 593.31 | 2 |

| | | | | |
|---|---|---|---|---|
| EFTU | GTVVTGTLER | P49411 | 516.79 | 2 |
| EFTU | DLEKPFLLPVEAVYSVPGR | P49411 | 710.39 | 3 |
| EFTU | GITINAAHVEYSTAAR | P49411 | 558.63 | 3 |
| EHD1 | HLIEQDFPGMR | Q9H4M9 | 671.83 | 2 |
| EHD2 | LFELEEQDLFR | Q9NZN4 | 719.86 | 2 |
| EHD4 | LFEAEAQDLFR | Q9H223 | 669.84 | 2 |
| EI2BE | [1Ac]-AAPVVAPPGVVVSR | Q13144 | 680.90 | 2 |
| EIF3A | VLLATLSIPITPER | Q14152 | 761.96 | 2 |
| EIF3A | TLSFGSDLNYATR | Q14152 | 722.86 | 2 |
| EIF3A | EDAPIGPHLQSMPSEQIR | Q14152 | 669.00 | 3 |
| EIF3A | LATLLGLQAPPTR | Q14152 | 675.91 | 2 |
| EIF3B | VTLMQLPTR | P55884 | 529.80 | 2 |
| EIF3B | DQYSVIFESGDR | P55884 | 708.33 | 2 |
| EIF3B | AQAVSEDAGGNEGR | P55884 | 680.81 | 2 |
| EIF3C | FEELTNLIR | Q99613 | 567.81 | 2 |
| EIF3C | ELLGQGLLLR | Q99613 | 556.35 | 2 |
| EIF3D | IFHTVTTTDDPVIR | O15371 | 538.95 | 3 |
| EIF3D | SVYSWDIVVQR | O15371 | 676.35 | 2 |
| EIF3D | LGDDIDLIVR | O15371 | 564.82 | 2 |
| EIF3F | VIGLSSDLQQVGGASAR | O00303 | 829.45 | 2 |
| EIF3F | LHPVILASIVDSYER | O00303 | 571.32 | 3 |
| EIF3G | VTNLSEDTR | O75821 | 517.76 | 2 |
| EIF3I | FFHLAFEEEFGR | Q13347 | 510.25 | 3 |
| EIF3L | LAGFLDLTEQEFR | Q9Y262 | 769.90 | 2 |
| EIF3L | YGDFFIR | Q9Y262 | 459.23 | 2 |
| EIF3L | IDESIHLQLR | Q9Y262 | 408.56 | 3 |
| ELAV1 | VLVDQTTGLSR | Q15717 | 594.83 | 2 |
| ELAV1 | DANLYISGLPR | Q15717 | 609.83 | 2 |
| ELOB | [1Ac]-MDVFLMIR | Q15370 | 533.77 | 2 |
| ELOV1 | [1Ac]-MEAVVNLYQEVMK | Q9BW60 | 798.39 | 2 |
| ELP3 | ILALVPPWTR | Q9H9T3 | 583.36 | 2 |
| EMAL4 | EIEVPDQYGTIR | Q9HC35 | 710.36 | 2 |
| ENOA | YISPDQLADLYK | P06733 | 713.37 | 2 |
| ENOA | VVIGMDVAASEFFR | P06733 | 770.90 | 2 |
| ENOA | IEEELGSK | P06733 | 452.73 | 2 |
| ENOA | LAMQEFMILPVGAANFR | P06733 | 954.50 | 2 |
| ENOA | YNQLLR | P06733 | 403.73 | 2 |
| ENOA | GNPTVEVDLFTSK | P06733 | 703.86 | 2 |
| ENOA | [1Ac]-SILK[Frm]IHAR | P06733 | 504.30 | 2 |
| ENPL | SILFVPTSAPR | P14625 | 594.34 | 2 |
| ENPL | FAFQAEVNR | P14625 | 541.27 | 2 |
| ENPL | GVVDSDDLPLNVSR | P14625 | 743.38 | 2 |
| ENPL | LSLNIDPDAK | P14625 | 543.30 | 2 |
| ENPL | SGYLLPDTK | P14625 | 497.27 | 2 |
| ENTP1 | SLSNYPFDFQGAR | P49961 | 751.36 | 2 |
| ERAP1 | ILASTQFEPTAAR | Q9NZ08 | 702.88 | 2 |
| ERAP1 | NPVGYPLAWQFLR | Q9NZ08 | 780.92 | 2 |
| ERF1 | LSVLGAITSVQQR | P62495 | 686.40 | 2 |
| ERH | ADTQTYQPYNK | P84090 | 664.81 | 2 |
| ERH | [1Ac]-SHTILLVQPTK | P84090 | 639.87 | 2 |
| ERO1A | FDGILTEGEGPR | Q96HE7 | 645.82 | 2 |
| ERP44 | DLAEITTLDR | Q9BS26 | 573.80 | 2 |
| ESTD | MYSYVTEELPQLINANFPVDPQR | P10768 | 908.78 | 3 |
| ESYT1 | ALTLGALTLPLAR | Q9BSJ8 | 655.41 | 2 |
| ESYT1 | LLAETVAPAVR | Q9BSJ8 | 570.34 | 2 |
| ESYT1 | FEWELPLDEAQR | Q9BSJ8 | 766.87 | 2 |
| ESYT2 | ALALLEDEER | A0FGR8 | 579.80 | 2 |
| ETFA | LEVAPISDIIAIK | P13804 | 691.42 | 2 |
| ETFB | EIDGGLETLR | P38117 | 551.79 | 2 |
| ETHE1 | EAVLIDPVLETAPR | O95571 | 761.93 | 2 |
| ETHE1 | GGSGAPILLR | O95571 | 470.78 | 2 |
| EZRI | QLLTLSSELSQAR | P15311 | 723.40 | 2 |
| EZRI | IALLEEAR | P15311 | 457.77 | 2 |
| F10A1 | AIEINPDSAQPYK | P50502 | 723.37 | 2 |
| F16P1 | APVILGSPDDVLEFLK | P09467 | 856.98 | 2 |
| F213A | VNLLSVLEAAK | Q9BRX8 | 578.85 | 2 |
| FA49B | GLLGALTSTPYSPTQHLER | Q9NUQ9 | 681.03 | 3 |
| FA49B | DAEGILEDLQSYR | Q9NUQ9 | 754.86 | 2 |
| FA49B | MTNPAIQNDFSYYR | Q9NUQ9 | 860.39 | 2 |
| FABP5 | ELGVGIALR | Q01469 | 464.28 | 2 |
| FABP5 | FEETTADGR | Q01469 | 513.23 | 2 |
| FABP5 | [1Ac]-ATVQQLEGR | Q01469 | 522.28 | 2 |
| FAH2A | ALAAQLPVLPR | Q96GK7 | 574.86 | 2 |
| FAS | RPTPQDSPIFLPVDDTSFR | P49327 | 730.04 | 3 |

| | | | | |
|---|---|---|---|---|
| FBLN1 | EFTRPEEIIFLR | P23142 | 517.28 | 3 |
| FBN1 | AGYQSTLTR | P35555 | 498.76 | 2 |
| FBRL | IVALNAHTFLR | P22087 | 418.92 | 3 |
| FEN1 | LIADVAPSAIR | P39748 | 563.33 | 2 |
| FIBA | MELERPGGNEITR | P02671 | 501.25 | 3 |
| FIBA | GLIDEVNQDFTNR | P02671 | 760.87 | 2 |
| FIBA | GGSTSYGTGSETESPR | P02671 | 786.84 | 2 |
| FIBA | GDFSSANNR | P02671 | 484.22 | 2 |
| FIBA | GSESGIFTNTK | P02671 | 570.78 | 2 |
| FIBA | [CRM]-MK[1Ac]GLIDEVNQDFTNR | P02671 | 622.30 | 3 |
| FIBB | IRPFFPQQ | P02675 | 516.78 | 2 |
| FIBB | QDGSVDFGR | P02675 | 490.73 | 2 |
| FIBB | EEAPSLRPAPPPISGGGYR | P02675 | 651.01 | 3 |
| FIBB | MGPTELLIEMEDWK | P02675 | 846.40 | 2 |
| FIBG | EGFGHLSPTGTTEFWLGNEK | P02679 | 736.35 | 3 |
| FIBG | IHLISTQSAIPYALR | P02679 | 561.66 | 3 |
| FIBG | YLQEIYNSNNQK | P02679 | 757.37 | 2 |
| FIBG | VELEDWNGR | P02679 | 559.27 | 2 |
| FIBG | TSTADYAMFK | P02679 | 567.76 | 2 |
| FINC | VDVIPVNLPGEHGQR | P02751 | 543.96 | 3 |
| FINC | VPGTSTSATLTGLTR | P02751 | 731.40 | 2 |
| FINC | SSPVVIDASTAIDAPSNLR | P02751 | 957.00 | 2 |
| FINC | FTNIGPDTMR | P02751 | 576.28 | 2 |
| FINC | WSRPQAPITGYR | P02751 | 477.92 | 3 |
| FIS1 | GIVLLEELLPK | Q9Y3D6 | 612.38 | 2 |
| FIS1 | [1Ac]-MEAVLNELVSVEDLLK | Q9Y3D6 | 922.49 | 2 |
| FKB1A | GWEEGVAQMSVGQR | P62942 | 767.36 | 2 |
| FLII | FVFLLDR | Q13045 | 455.26 | 2 |
| FLII | ADLTALFLPR | Q13045 | 558.82 | 2 |
| FLII | [1Ac]-MEATGVLPFVR | Q13045 | 631.33 | 2 |
| FLNA | IANLQTDLSDGLR | P21333 | 708.38 | 2 |
| FLNA | YAPSEAGLHEMDIR | P21333 | 530.25 | 3 |
| FLNA | LPQLPITNFSR | P21333 | 643.37 | 2 |
| FLOT1 | AQQVAVQEQEIAR | O75955 | 735.39 | 2 |
| FMR1 | [1Ac]-MEELVVEVR | Q06787 | 573.30 | 2 |
| FPPS | VLTEDEMGHPEIGDAIAR | P14324 | 651.65 | 3 |
| FPPS | GLTVVVAFR | P14324 | 481.30 | 2 |
| FRIH | [1Ac]-TTASTSQVR | P02794 | 496.75 | 2 |
| FRIL | LGGPEAGLGEYLFER | P02792 | 804.41 | 2 |
| FRIL | LNQALLDLHALGSAR | P02792 | 531.30 | 3 |
| FSCN1 | LVARPEPATGYTLEFR | Q16658 | 607.33 | 3 |
| FUBP1 | IGGNEGIDVPIPR | Q96AE4 | 668.86 | 2 |
| FUBP1 | IQIAPDSGGLPER | Q96AE4 | 676.86 | 2 |
| FUBP1 | IAQITGPPDR | Q96AE4 | 534.30 | 2 |
| FUBP2 | IGGGIDVPVPR | Q92945 | 540.31 | 2 |
| FUBP2 | VPDGMVGLIIGR | Q92945 | 613.85 | 2 |
| FUBP2 | SVSLTGAPESVQK | Q92945 | 651.85 | 2 |
| FUCO | DLVGELGTALR | P04066 | 572.32 | 2 |
| FUMH | IYELAAGGTAVGTGLNTR | P07954 | 882.47 | 2 |
| G3BP1 | EAGEQGDIEPR | Q13283 | 600.78 | 2 |
| G3P | VIHDNFGIVEGLMTTVHAITATQK | P04406 | 649.60 | 4 |
| G3P | VIISAPSADAPMFVMGVNHEK | P04406 | 738.37 | 3 |
| G3P | LISWYDNEFGYSNR | P04406 | 882.40 | 2 |
| G3P | LVINGNPITIFQER | P04406 | 807.45 | 2 |
| G3P | [1Ac]-DP[Oxi]SK[1Ac]IK[1Ac]WGDAGAEYVVESTGVFTTMEK | P04406 | 772.62 | 4 |
| G6PD | EMVQNLMVLR | P11413 | 616.83 | 2 |
| G6PD | LFYLALPPTVYEAVTK | P11413 | 913.01 | 2 |
| G6PD | [1Ac]-AEQVALSR | P11413 | 458.25 | 2 |
| G6PI | ILLANFLAQTEALMR | P06744 | 852.48 | 2 |
| G6PI | NLVTEDVMR | P06744 | 538.77 | 2 |
| G6PI | TLAQLNPESSLFIIASK | P06744 | 916.51 | 2 |
| G6PI | TFTTQETITNAETAK | P06744 | 828.41 | 2 |
| GANAB | LSFQHDPETSVLVLR | Q14697 | 580.98 | 3 |
| GANAB | YRVPDVLVADPPIAR | Q14697 | 560.99 | 3 |
| GANAB | YSLLPFWYTLLYQAHR | Q14697 | 691.03 | 3 |
| GANAB | IDELEPR | Q14697 | 436.23 | 2 |
| GANAB | YFTWDPSR | Q14697 | 536.25 | 2 |
| GAPR1 | EAQQYSEALASTR | Q9H4G4 | 727.35 | 2 |
| GBLP | LWDLTTGTTTR | P63244 | 632.83 | 2 |
| GBLP | DETNYGIPQR | P63244 | 596.78 | 2 |
| GBLP | DVLSVAFSSDNR | P63244 | 655.32 | 2 |
| GBP1 | EAIEVFIR | P32455 | 488.78 | 2 |
| GCN1L | NPEILAIAPVLLDALTDPSR | Q92616 | 706.73 | 3 |
| GDIA | FQLLEGPPESMGR | P31150 | 730.86 | 2 |

| | | | | |
|---|---|---|---|---|
| GDIB | FVSISDLLVPK | P50395 | 609.36 | 2 |
| GDIB | DLGTESQIFISR | P50395 | 683.35 | 2 |
| GDIB | VPSTEAEALASSLMGLFEK | P50395 | 660.67 | 3 |
| GDIB | MTGSEFDFEEMKR | P50395 | 536.23 | 3 |
| GDIR1 | AEEYEFLTPVEEAPK | P52565 | 876.42 | 2 |
| GDIR1 | TDYMVGSYGPR | P52565 | 623.28 | 2 |
| GDIR2 | APNVVVTR | P52566 | 428.26 | 2 |
| GDIR2 | TLLGDGPVVTDPK | P52566 | 656.36 | 2 |
| GELS | TGAQELLR | P06396 | 444.25 | 2 |
| GELS | EVQGFESATFLGYFK | P06396 | 861.92 | 2 |
| GELS | YIETDPANR | P06396 | 539.76 | 2 |
| GGACT | [1Ac]-ALVFVYGTLK | Q9BVM4 | 576.84 | 2 |
| GGT5 | DLLGETLAQLIR | P36269 | 671.39 | 2 |
| GIMA1 | FTAQDQQAVR | Q8WWP7 | 582.29 | 2 |
| GLRX1 | AQEILSQLPIK | P35754 | 620.37 | 2 |
| GLRX3 | [1Ac]-AAGAAEAAVAAVEEVGSAGQFEELLR | O76003 | 853.43 | 3 |
| GLRX3 | [1Ac]-AAGAAEAAVAAVE[NaX]EVGSAGQFEELLR | O76003 | 860.76 | 3 |
| GLRX3 | [1Ac]-AAGAAEAAVAAVEE[NaX]VGSAGQFEELLR | O76003 | 860.76 | 3 |
| GLU2B | ESLQQMAEVTR | P14314 | 646.32 | 2 |
| GLU2B | ETMVTSTTEPSR | P14314 | 669.81 | 2 |
| GLYG | LVVLATPQVSDSMR | P46976 | 758.41 | 2 |
| GLYM | LGAPALTSR | P34897 | 443.26 | 2 |
| GLYM | LIIAGTSAYAR | P34897 | 568.33 | 2 |
| GLYM | AMADALLER | P34897 | 495.26 | 2 |
| GMIP | TPLFGVDFLQLPR | Q9P107 | 751.92 | 2 |
| GMIP | VTLSLFGLR | Q9P107 | 503.31 | 2 |
| GNA13 | VFLQYLPAIR | Q14344 | 610.36 | 2 |
| GNAI2 | IAQSDYIPTQQDVLR | P04899 | 873.95 | 2 |
| GNAI2 | AVVYSNTIQSIMAIVK | P04899 | 868.98 | 2 |
| GNAI2 | AMGNLQIDFADPSR | P04899 | 767.87 | 2 |
| GNAI3 | ISQSNYIPTQQDVLR | P08754 | 881.46 | 2 |
| GOT1B | VPVLGSLLNLPGIR | Q9Y3E0 | 724.45 | 2 |
| GPDM | SEISLLPSDIDR | P43304 | 672.85 | 2 |
| GPX1 | DYTQMNELQR | P07203 | 649.30 | 2 |
| GRHPR | NTMSLLAANNLLAGLR | Q9UBQ7 | 836.46 | 2 |
| GRHPR | ILDAAGANLK | Q9UBQ7 | 493.29 | 2 |
| GRHPR | ETAVFINISR | Q9UBQ7 | 575.32 | 2 |
| GRP75 | AQFEGIVTDLIR | P38646 | 681.37 | 2 |
| GRP75 | VQQTVQDLFGR | P38646 | 645.84 | 2 |
| GRP75 | DAGQISGLNVLR | P38646 | 621.84 | 2 |
| GRP75 | VINEPTAAALAYGLDK | P38646 | 823.44 | 2 |
| GRP75 | TTPSVVAFTADGER | P38646 | 725.86 | 2 |
| GRP75 | SDIGEVILVGGMTR | P38646 | 723.88 | 2 |
| GRP78 | ITPSYVAFTPEGER | P11021 | 783.89 | 2 |
| GRP78 | DAGTIAGLNVMR | P11021 | 609.32 | 2 |
| GRP78 | ITITNDQNR | P11021 | 537.78 | 2 |
| GRP78 | NQLTSNPENTVFDAK | P11021 | 839.41 | 2 |
| GRP78 | TFAPEEISAMVLTK | P11021 | 768.90 | 2 |
| GSHR | ALLTPVAIAAGR | P00390 | 576.86 | 2 |
| GSLG1 | LLELQYFISR | Q92896 | 641.36 | 2 |
| GSTO1 | VPSLVGSFIR | P78417 | 537.82 | 2 |
| GSTP1 | FQDGDLTLYQSNTILR | P09211 | 942.48 | 2 |
| GSTP1 | DQQEAALVDMVNDGVEDLR | P09211 | 706.33 | 3 |
| GT251 | VLIALLAR | Q8NBJ5 | 434.80 | 2 |
| GTF2I | MSVDAVEIETLR | P78347 | 681.85 | 2 |
| H12 | ASGPPVSELITK | P16403 | 599.84 | 2 |
| H12 | ALAAAGYDVEK | P16403 | 554.29 | 2 |
| H12 | KASGPPVSELITK | P16403 | 442.93 | 3 |
| H12 | RKASGPPVSELITK | P16403 | 494.96 | 3 |
| H13 | [1Ac]-SETAPLAPTIPAPAEK | P16402 | 817.94 | 2 |
| H13 | [1Ac]-SETAPLAPTIPAPAEKTP[PGP]VKKK | P16402 | 777.44 | 3 |
| H13 | [1Ac]-SETAPLAPTIPAPAEKTPVKKK[Frm] | P16402 | 782.11 | 3 |
| H15 | ATGPPVSELITK | P16401 | 606.85 | 2 |
| H15 | ALAAGGYDVEK | P16401 | 547.28 | 2 |
| H15 | NGLSLAALK | P16401 | 443.77 | 2 |
| H15 | KATGPPVSELITK | P16401 | 447.60 | 3 |
| H15 | [1Ac]-SETAPAETATPAPVEK | P16401 | 820.90 | 2 |
| H15 | K[1Ac]ATGP[Oxi]PVSELITK | P16401 | 466.93 | 3 |
| H1X | YSQLVVETIR | Q92522 | 604.34 | 2 |
| H1X | [1Ac]-SVELEEALPVTTAEGMAK | Q92522 | 958.98 | 2 |
| H2A2A | VGAGAPVYMAAVLEYLTAEILELAGNAAR | Q6FI13 | 978.52 | 3 |
| H2A2B | HLQLAVR | Q8IUE6 | 418.76 | 2 |
| H2A2C | [1Ac]-VGAGAPVY[Amn]MAAVLEYLTAEILELAGNAAR | Q16777 | 997.53 | 3 |
| H2AW | SARAGVIFPVGR | Q9P0M6 | 615.36 | 2 |

| | | | | |
|---|---|---|---|---|
| H2AY | SIAFPSIGSGR | O75367 | 546.30 | 2 |
| H2AY | GVTIASGGVLPNIHPELLAK | O75367 | 662.72 | 3 |
| H2AY | EFVEAVLELR | O75367 | 602.83 | 2 |
| H2AY | [AAR]-GVGAPVYMAAVLEYLT[Pho]AEILELAGNAAR | O75367 | 743.38 | 4 |
| H2AY | [1Ac]-IGVGAPVYM[DTM]AAVLEYLTAEILELAGNAAR | O75367 | 990.54 | 3 |
| H2AZ | VGATAAVYSAAILEYLTAEVLELAGNASK | P0C0S5 | 965.85 | 3 |
| H2AZ | GDEELDSLIK | P0C0S5 | 559.78 | 2 |
| H2B1M | AMGIMNSFVNDIFER | Q99879 | 872.41 | 2 |
| H2B1M | QVHPDTGISSK | Q99879 | 584.80 | 2 |
| H2B1M | KESYSVYVYK | Q99879 | 633.32 | 2 |
| H2B2C | KESYSIYVYK | Q6DN03 | 640.33 | 2 |
| H4 | ISGLIYEETR | P62805 | 590.81 | 2 |
| H4 | VFLENVIR | P62805 | 495.29 | 2 |
| H4 | TVTAMDVVYALK | P62805 | 655.85 | 2 |
| H4 | DNIQGITKPAIR | P62805 | 442.59 | 3 |
| H4 | DAVTYTEHAK | P62805 | 567.77 | 2 |
| H4 | KTVTAMDVVYALK | P62805 | 480.27 | 3 |
| H4 | GLGK[1Ac]GGAK[1Ac]R | P62805 | 464.27 | 2 |
| H4 | GGK[1Ac]GLGK[1Ac]GGAK[1Ac]R | P62805 | 606.35 | 2 |
| H4 | GK[1Ac]GGK[1Ac]GLGK[1Ac]GGAK[1Ac]R | P62805 | 719.91 | 2 |
| HACD3 | LESEGSPETLTNLR | Q9P035 | 773.39 | 2 |
| HAT1 | LLVTDMSDAEQYR | O14929 | 770.87 | 2 |
| HBA | MFLSFPTTK | P69905 | 536.28 | 2 |
| HBA | FLASVSTVLTSK | P69905 | 626.86 | 2 |
| HBA | VGAHAGEYGAEALER | P69905 | 510.58 | 3 |
| HBA | VADALTNAVAHVDDMPNALSALSDLHAHK | P69905 | 600.10 | 5 |
| HBA | TYFPHFDLSHGSAQVK | P69905 | 459.23 | 4 |
| HBB | VNVDEVGGEALGR | P68871 | 657.84 | 2 |
| HBB | SAVTALWGK | P68871 | 466.76 | 2 |
| HBB | EFTPPVQAAYQK | P68871 | 689.85 | 2 |
| HBB | FFESFGDLSTPDAVMGNPK | P68871 | 686.99 | 3 |
| HBB | [CRM]-SAVTALWGK[1Ac]VNVDEVGGEALGR | P68871 | 771.73 | 3 |
| HBD | VNVDAVGGEALGR | P02042 | 628.83 | 2 |
| HCD2 | GGIVGMTLPIAR | Q99714 | 592.84 | 2 |
| HCD2 | VLDVNLMGTFNVIR | Q99714 | 795.94 | 2 |
| HCD2 | VMTIAPGLFGTPLLTSLPEK | Q99714 | 695.73 | 3 |
| HCD2 | GLVAVITGGASGLGLATAER | Q99714 | 605.01 | 3 |
| HCD2 | [CRM]-SVK[1Ac]GLVAVITGGASGLGLATAER | Q99714 | 738.08 | 3 |
| HCFC1 | [1Ac]-ASAVSPANLPAVLLQPR | P51610 | 873.50 | 2 |
| HEMO | LWWLDLK | P02790 | 487.28 | 2 |
| HEMO | EWFWDLATGTMK | P02790 | 742.85 | 2 |
| HG2A | DLISNNEQLPMLGR | P04233 | 800.41 | 2 |
| HG2A | LTVTSQNLQLENLR | P04233 | 814.95 | 2 |
| HG2A | [+1R]-DK[1Ac]LTVTSQNLQLENLR | P04233 | 690.71 | 3 |
| HNRH1 | EGRPSGEAFVELESEDEVK | P31943 | 703.00 | 3 |
| HNRH1 | [1Ac]-MMLGTEGGEGFVVK | P31943 | 748.86 | 2 |
| HNRH1 | [1Ac]-MLGTEGGEGFVVK | P31943 | 683.34 | 2 |
| HNRH2 | ATENDIYNFFSPLNPMR | P55795 | 676.99 | 3 |
| HNRH3 | YIEIFR | P31942 | 420.73 | 2 |
| HNRH3 | ATENDIANFFSPLNPIR | P31942 | 959.99 | 2 |
| HNRH3 | STGEAFVQFASK | P31942 | 636.32 | 2 |
| HNRL1 | NYILDQTNVYGSAQR | Q9BUJ2 | 871.43 | 2 |
| HNRL2 | YNVLGAETVLNQMR | Q1KMD3 | 804.41 | 2 |
| HNRL2 | FPTLWSGAR | Q1KMD3 | 517.77 | 2 |
| HNRPC | VFIGNLNTLVVK | P07910 | 658.90 | 2 |
| HNRPC | GFAFVQYVNER | P07910 | 665.33 | 2 |
| HNRPC | MIAGQVLDINLAAEPK | P07910 | 841.96 | 2 |
| HNRPC | AAVAGEDGR | P07910 | 423.21 | 2 |
| HNRPC | GDDLQAIK | P07910 | 430.23 | 2 |
| HNRPD | IFVGGLSPDTPEEK | Q14103 | 744.88 | 2 |
| HNRPD | MFIGGLSWDTTK | Q14103 | 678.34 | 2 |
| HNRPF | [1Ac]-MLGPEGGEGFVVK | P52597 | 681.34 | 2 |
| HNRPK | IILDLISESPIK | P61978 | 670.91 | 2 |
| HNRPK | LLIHQSLAGGIIGVK | P61978 | 506.98 | 3 |
| HNRPK | ILSISADIETIGEILK | P61978 | 858.00 | 2 |
| HNRPK | GSYGDLGGPIITTQVTIPK | P61978 | 959.02 | 2 |
| HNRPK | IITITGTQDQIQNAQYLLQNSVK | P61978 | 863.80 | 3 |
| HNRPL | SSSGLLEWESK | P14866 | 611.80 | 2 |
| HNRPM | AFITNIPFDVK | P52272 | 632.85 | 2 |
| HNRPM | MGPAMGPALGAGIER | P52272 | 714.36 | 2 |
| HNRPM | GNFGGSFAGSFGGAGGHAPGVAR | P52272 | 678.99 | 3 |
| HNRPM | MGPLGLDHMASSIER | P52272 | 538.60 | 3 |
| HNRPM | MGAGMGFGLER | P52272 | 563.26 | 2 |
| HNRPM | LGSTVFVANLDYK | P52272 | 713.88 | 2 |

| | | | | |
|---|---|---|---|---|
| HNRPM | [1Ac]-AAGVEAAAEVAATEIK | P52272 | 771.90 | 2 |
| HNRPQ | LMMDPLTGLNR | O60506 | 630.83 | 2 |
| HNRPQ | NLANTVTEEILEK | O60506 | 737.39 | 2 |
| HNRPQ | DLFEDELVPLFEK | O60506 | 797.41 | 2 |
| HNRPR | NLATTVTEEILEK | O43390 | 730.90 | 2 |
| HNRPR | LMMDPLSGQNR | O43390 | 631.30 | 2 |
| HNRPR | DLYEDELVPLFEK | O43390 | 805.40 | 2 |
| HNRPU | FIEIAAR | Q00839 | 410.24 | 2 |
| HNRPU | NFILDQTNVSAAAQR | Q00839 | 824.43 | 2 |
| HNRPU | SSGPTSLFAVTVAPPGAR | Q00839 | 857.96 | 2 |
| HNRPU | DIDIHEVR | Q00839 | 498.76 | 2 |
| HP1B3 | [1Ac]-ATDTSQGELVHPK | Q5SSJ5 | 712.85 | 2 |
| HPRT | SIPMTVDFIR | P00492 | 589.82 | 2 |
| HPT | VGYVSGWGR | P00738 | 490.75 | 2 |
| HPT | DIAPTLTLYVGK | P00738 | 645.87 | 2 |
| HPT | HYEGSTVPEK | P00738 | 573.77 | 2 |
| HS105 | AGGIETIANEFSDR | Q92598 | 740.36 | 2 |
| HS71B | IINEPTAAAIAYGLDR | P0DMV9 | 844.45 | 2 |
| HS71B | DAGVIAGLNVLR | P0DMV9 | 599.35 | 2 |
| HS90A | APFDLFENR | P07900 | 554.77 | 2 |
| HS90A | RAPFDLFENR | P07900 | 632.83 | 2 |
| HS90A | TDTGEPMGR | P07900 | 482.21 | 2 |
| HS90A | ELISNSSDALDK | P07900 | 646.32 | 2 |
| HS90B | YHTSQSGDEMTSLSEYVSR | P08238 | 726.32 | 3 |
| HS90B | EQVANSAFVER | P08238 | 625.31 | 2 |
| HS90B | HLEINPDHPIVETLR | P08238 | 594.99 | 3 |
| HS90B | NPDDITQEEYGEFYK | P08238 | 924.40 | 2 |
| HS90B | [Frm]-VFIM[Oxi]DS[Pho]CDELIPEYLNFIR | P08238 | 611.03 | 4 |
| HSP74 | EFSITDVVPYPISLR | P34932 | 868.47 | 2 |
| HSP74 | AGGIETIANEYSDR | P34932 | 748.35 | 2 |
| HSP74 | NFTTEQVTAMLLSK | P34932 | 791.91 | 2 |
| HSP7C | FEELNADLFR | P11142 | 627.31 | 2 |
| HSP7C | DAGTIAGLNVLR | P11142 | 600.34 | 2 |
| HSP7C | GTLDPVEK | P11142 | 429.73 | 2 |
| HSP7C | SFYPEEVSSMVLTK | P11142 | 808.90 | 2 |
| HSP7C | SQIHDIVLVGGSTR | P11142 | 494.61 | 3 |
| HSP7C | ARFEELNADLFR | P11142 | 494.26 | 3 |
| HSPB1 | LFDQAFGLPR | P04792 | 582.31 | 2 |
| HSPB1 | GPSWDPFR | P04792 | 481.23 | 2 |
| HSPB1 | QLSSGVSEIR | P04792 | 538.29 | 2 |
| HSPB1 | LATQSNEITIPVTFESR | P04792 | 953.50 | 2 |
| HSPB1 | QLSS[Pho]GVSEIR | P04792 | 578.27 | 2 |
| HUWE1 | LLGPSAAADILQLSSSLPLQSR | Q7Z6Z7 | 746.42 | 3 |
| HUWE1 | SLLSILQR | Q7Z6Z7 | 465.29 | 2 |
| HVCN1 | AAAPDVAPAPGPAPR | Q96D96 | 679.36 | 2 |
| HXK1 | LSDETLIDIMTR | P19367 | 703.86 | 2 |
| HXK1 | LALLQVR | P19367 | 406.77 | 2 |
| HXK1 | SANLVAATLGAILNR | P19367 | 742.43 | 2 |
| HXK1 | LVPDSDVR | P19367 | 450.74 | 2 |
| HXK1 | MVSGMYLGELVR | P19367 | 677.85 | 2 |
| HYEP | GGHFAAFEEPELLAQDIR | P07099 | 667.33 | 3 |
| HYOU1 | LAGLFNEQR | Q9Y4L1 | 524.28 | 2 |
| HYOU1 | DAVVYPILVEFTR | Q9Y4L1 | 761.42 | 2 |
| IC1 | LLDSLPSDTR | P05155 | 558.80 | 2 |
| ICAM1 | EPAVGEPAEVTTTVLVR | P05362 | 884.48 | 2 |
| IDH3A | IAEFAFEYAR | P50213 | 608.80 | 2 |
| IDH3A | DMANPTALLLSAVMMLR | P50213 | 616.32 | 3 |
| IDHC | LVSGWVKPIIIGR | O75874 | 479.97 | 3 |
| IDHC | SDYLNTFEFMDK | O75874 | 755.33 | 2 |
| IDHP | LIDDMVAQVLK | P48735 | 622.85 | 2 |
| IDHP | TIEAEAAHGTVTR | P48735 | 452.57 | 3 |
| IDHP | YFDLGLPNR | P48735 | 547.79 | 2 |
| IF2A | YVMTTTTLER | P05198 | 607.81 | 2 |
| IF2A | VVTDTDETELAR | P05198 | 674.83 | 2 |
| IF2A | TEGLSVLSQAMAVIK | P05198 | 773.93 | 2 |
| IF2B | DYTYEELLNR | P20042 | 658.31 | 2 |
| IF2G | [1Ac]-AGGEAGVTLGQPHLSR | P41091 | 796.41 | 2 |
| IF4A1 | ATQALVLAPTR | P60842 | 570.84 | 2 |
| IF4A1 | LQMEAPHIIVGTPGR | P60842 | 540.30 | 3 |
| IF4A2 | ETQALVLAPTR | Q14240 | 599.84 | 2 |
| IF4A3 | DVIAQSQSGTGK | P38919 | 595.80 | 2 |
| IF4A3 | [1Ac]-ATTATMATSGSAR | P38919 | 634.30 | 2 |
| IF4B | TGSESSQTGTSTTSSR | P23588 | 787.35 | 2 |
| IF4G1 | FMLQDVLDLR | Q04637 | 625.33 | 2 |

| | | | | |
|---|---|---|---|---|
| IF4H | EALTYDGALLGDR | Q15056 | 697.35 | 2 |
| IF5 | VNILFDFVK | P55010 | 547.82 | 2 |
| IF5A1 | VHLVGIDIFTGK | P63241 | 433.59 | 3 |
| IF5A1 | NDFQLIGIQDGYLSLLQDSGEVR | P63241 | 860.77 | 3 |
| IGHM | QIQVSWLR | P01871 | 515.30 | 2 |
| IGJ | SSEDPNEDIVER | P01591 | 695.31 | 2 |
| IGKC | TVAAPSVFIFPPSDEQLK | P01834 | 973.52 | 2 |
| IGKC | DSTYSLSSTLTLSK | P01834 | 751.88 | 2 |
| IKZF1 | AASENSQDALR | Q13422 | 581.28 | 2 |
| IL16 | GLPDPALSTQPAPASR | Q14005 | 789.42 | 2 |
| ILEU | [1Ac]-MEQLSSANTR | P30740 | 589.78 | 2 |
| ILF2 | VLQSALAAIR | Q12905 | 521.32 | 2 |
| ILF2 | ILPTLEAVAALGNK | Q12905 | 705.42 | 2 |
| ILF2 | INNVIDNLIVAPGTFEVQIEEVR | Q12905 | 861.47 | 3 |
| ILF3 | VLQDMGLPTGAEGR | Q12906 | 722.37 | 2 |
| ILF3 | VLAGETLSVNDPPDVLDR | Q12906 | 955.50 | 2 |
| ILK | SAVVEMLIMR | Q13418 | 574.81 | 2 |
| ILK | GMAFLHTLEPLIPR | Q13418 | 532.30 | 3 |
| IMB1 | AAVENLPTFLVELSR | Q14974 | 829.96 | 2 |
| IMB1 | VLANPGNSQVAR | Q14974 | 613.34 | 2 |
| IMB1 | SNEILTAIIQGMR | Q14974 | 723.39 | 2 |
| IMB1 | LQQVLQMESHIQSTSDR | Q14974 | 667.33 | 3 |
| IMB1 | [1Ac]-MELITILEK | Q14974 | 566.32 | 2 |
| IMDH2 | LVGIISSR | P12268 | 422.77 | 2 |
| IMDH2 | NLIDAGVDALR | P12268 | 578.82 | 2 |
| IMDH2 | EDLVVAPAGITLK | P12268 | 663.39 | 2 |
| IPO5 | QLALEVIVTLSETAAAMLR | O00410 | 677.05 | 3 |
| IPO5 | ITFLLQAIR | O00410 | 537.84 | 2 |
| IPO5 | [1Ac]-AAAAAEQQQFYLLLGNLLSPDNVVR | O00410 | 915.15 | 3 |
| IPO5 | [1Ac]-AAAAAE[KXX]QQQFYLLLGNLLSPDNVVR | O00410 | 927.80 | 3 |
| IPO7 | ENIVEAIIHSPELIR | O95373 | 578.32 | 3 |
| IPO7 | [1Ac]-MDPNTIIEALR | O95373 | 657.84 | 2 |
| IPO7 | [1Ac]-MDPN[Dea]TIIEALR | O95373 | 658.33 | 2 |
| IPYR | LKPGYLEATVDWFR | Q15181 | 565.63 | 3 |
| IQGA1 | EEVITLIR | P46940 | 486.79 | 2 |
| IQGA1 | EQLWLANEGLITR | P46940 | 771.92 | 2 |
| IQGA1 | LEGVLAEVAQHYQDTLIR | P46940 | 685.70 | 3 |
| IQGA1 | TLINAEDPPMVVVR | P46940 | 777.42 | 2 |
| IQGA1 | [1Ac]-SAADEVDGLGVAR | P46940 | 651.32 | 2 |
| ISG20 | LEILQLLK | Q96AZ6 | 485.32 | 2 |
| IST1 | TNQIGTVNDR | P53990 | 559.28 | 2 |
| ITAM | SLPISLVFLVPVR | P11215 | 720.45 | 2 |
| ITB2 | ALNEITESGR | P05107 | 545.28 | 2 |
| ITB2 | YLIYVDESR | P05107 | 579.30 | 2 |
| ITIH2 | SSALDMENFR | P19823 | 585.27 | 2 |
| ITIH4 | LGVYELLLK | Q14624 | 524.33 | 2 |
| KAD2 | AVLLGPPGAGK | P54819 | 490.30 | 2 |
| KAD2 | LQAYHTQTTPLIEYYR | P54819 | 666.34 | 3 |
| KAP0 | LGPSDYFGEIALLMNRPR | P10644 | 683.69 | 3 |
| KAP0 | RSENEEFVEVGR | P10644 | 484.24 | 3 |
| KAP0 | [1Ac]-MESGSTAASEEAR | P10644 | 684.29 | 2 |
| KAP2 | [1Ac]-SHIQIPPGLTELLQGYTVEVLR | P13861 | 835.80 | 3 |
| KCD12 | DLQLVLPDYFPER | Q96CX2 | 802.92 | 2 |
| KCD12 | YILDYLR | Q96CX2 | 478.27 | 2 |
| KCD12 | MFTQQQPQELAR | Q96CX2 | 738.87 | 2 |
| KCD12 | LGAPQQPGPGPPPSR | Q96CX2 | 728.39 | 2 |
| KCD12 | SPSGGAAGPLLTPSQSLDGSR | Q96CX2 | 978.00 | 2 |
| KPRA | GQDIFIIQTIPR | Q14558 | 700.90 | 2 |
| KPYM | APIIAVTR | P14618 | 420.77 | 2 |
| KPYM | LDIDSPPITAR | P14618 | 599.33 | 2 |
| KPYM | GDLGIEIPAEK | P14618 | 571.31 | 2 |
| KPYM | VNFAMNVGK | P14618 | 490.26 | 2 |
| KPYM | TATESFASDPILYRPVAVALDTK | P14618 | 822.44 | 3 |
| KPYM | TATESFASDPILYRPVAVALDTK[1Ac]GPEIRT-[Ami] | P14618 | 790.67 | 4 |
| KV313 | FSGSGSGTDFTLTISR | P18136 | 816.90 | 2 |
| LA | LTTDFNVIVEALSK | P05455 | 775.43 | 2 |
| LA | GSIFVVFDSIESAK | P05455 | 749.89 | 2 |
| LA | GQVLNIQMR | P05455 | 529.79 | 2 |
| LAC2 | AGVETTTPSK | P0CG05 | 495.76 | 2 |
| LAMC1 | LSAEDLVLEGAGLR | P11047 | 721.90 | 2 |
| LAMC1 | NTIEETGNLAEQAR | P11047 | 773.38 | 2 |
| LAMP2 | IPLNDLFR | P13473 | 494.28 | 2 |
| LAMP2 | GILTVDELLAIR | P13473 | 656.90 | 2 |
| LAP2B | SSTPLPTISSSAENTR | P42167 | 824.41 | 2 |

| | | | | |
|---|---|---|---|---|
| LAP2B | YGVNPGPIVGTTR | P42167 | 665.86 | 2 |
| LAP2B | HASPILPITEFSDIPR | P42167 | 598.32 | 3 |
| LAP2B | SST[Pho]PLPTISSSAENTR | P42167 | 864.40 | 2 |
| LASP1 | GFSVVADTPELQR | Q14847 | 709.87 | 2 |
| LASP1 | MGPSGGEGMEPER | Q14847 | 667.28 | 2 |
| LASP1 | QSFTMVADTPENLR | Q14847 | 804.89 | 2 |
| LBR | VVEGTPLIDGR | Q14739 | 578.32 | 2 |
| LC7L3 | YLQSLLAEVER | O95232 | 660.86 | 2 |
| LC7L3 | [1Ac]-MISAAQLLDELMGR | O95232 | 795.40 | 2 |
| LCAP | LPTAVVPLR | Q9UIQ6 | 483.31 | 2 |
| LDAH | AFLLFPTIER | Q9H6V9 | 603.85 | 2 |
| LDHA | VTLTSEEEAR | P00338 | 567.79 | 2 |
| LDHA | DLADELALVDVIEDK | P00338 | 829.43 | 2 |
| LDHA | FIIPNVVK | P00338 | 465.29 | 2 |
| LDHA | GEMMDLQHGSLFLR | P00338 | 545.27 | 3 |
| LDHA | QVVESAYEVIK | P00338 | 632.84 | 2 |
| LDHA | SADTLWGIQK | P00338 | 559.80 | 2 |
| LDHA | [1Ac]-ATLK[Frm]DQLIYNLLK | P00338 | 801.96 | 2 |
| LDHA | NCK[1Ac]LLIVSNPVDILTYVAWK | P00338 | 777.76 | 3 |
| LDHB | IVVVTAGVR | P07195 | 457.30 | 2 |
| LDHB | SLADELALVDVLEDK | P07195 | 815.43 | 2 |
| LDHB | FIIPQIVK | P07195 | 479.31 | 2 |
| LDHB | GLTSVINQK | P07195 | 480.28 | 2 |
| LDHB | MVVESAYEVIK | P07195 | 634.33 | 2 |
| LEG3 | IALDFQR | P17931 | 431.74 | 2 |
| LEG3 | GNDVAFHFNPR | P17931 | 425.21 | 3 |
| LEG9 | SILLSGTVLPSAQR | O00182 | 721.42 | 2 |
| LIMS2 | VIEGDVVSALNK | Q7Z4I7 | 622.35 | 2 |
| LKHA4 | DLSSHQLNEFLAQTLQR | P09960 | 667.34 | 3 |
| LKHA4 | TLTGTAALTVQSQEDNLR | P09960 | 640.00 | 3 |
| LKHA4 | FTRRTLT[Pho]GTAALTVQSQEDNLR | P09960 | 640.32 | 4 |
| LMNA | AAYEAELGDAR | P02545 | 583.28 | 2 |
| LMNA | LADALQELR | P02545 | 514.79 | 2 |
| LMNA | ITESEEVVSR | P02545 | 574.79 | 2 |
| LMNA | SLETENAGLR | P02545 | 545.28 | 2 |
| LMNA | EGDLIAAQAR | P02545 | 522.28 | 2 |
| LMNA | TALINSTGEEVAMR | P02545 | 746.38 | 2 |
| LMNB1 | AGGPTTPLSPTR | P20700 | 577.81 | 2 |
| LMNB1 | IQELEDLLAK | P20700 | 586.33 | 2 |
| LMNB1 | ALYETELADAR | P20700 | 626.31 | 2 |
| LMNB1 | LSSEMNTSTVNSAR | P20700 | 748.85 | 2 |
| LMNB1 | IESLSSQLSNLQK | P20700 | 723.89 | 2 |
| LMNB1 | LALDMEISAYR | P20700 | 641.33 | 2 |
| LMNB2 | GLESDVAELR | Q03252 | 544.78 | 2 |
| LONM | ILEFIAVSQLR | P36776 | 644.88 | 2 |
| LONM | DIIALNPLYR | P36776 | 594.34 | 2 |
| LPPRC | SSLLLGFR | P42704 | 446.77 | 2 |
| LPPRC | TVLDQQQTPSR | P42704 | 636.83 | 2 |
| LRMP | AEMLGAINQESR | Q12912 | 659.82 | 2 |
| LSM3 | GDGVVLVAPPLR | P62310 | 596.86 | 2 |
| LSM8 | TVAVITSDGR | O95777 | 509.78 | 2 |
| LSP1 | WETGEVQAQSAAK | P33241 | 702.84 | 2 |
| LSP1 | [1Ac]-AEASSDPGAEER | P33241 | 630.77 | 2 |
| LTOR3 | ELAPLFEELR | Q9UHA4 | 608.83 | 2 |
| LUM | FNALQYLR | P51884 | 512.78 | 2 |
| LYPA1 | ASFPQGPIGGANR | O75608 | 636.33 | 2 |
| LYPA1 | [1Ac]-STPLPAIVPAAR | O75608 | 617.86 | 2 |
| LYRIC | EMLSVGLGFLR | Q86UE4 | 611.34 | 2 |
| LYSC | STDYGIFQINSR | P61626 | 700.84 | 2 |
| M2OM | GIYTGLSAGLLR | Q02978 | 610.85 | 2 |
| M2OM | LGIYTVLFER | Q02978 | 605.85 | 2 |
| M2OM | [1Ac]-AATASAGAGGIDGKPR | Q02978 | 721.37 | 2 |
| MAOM | ALTSQLTDEELAQGR | P23368 | 816.42 | 2 |
| MAP4 | TTTLSGTAPAAGVVPSR | P27816 | 793.43 | 2 |
| MAP4 | TTTAAAVASTGPSSR | P27816 | 689.35 | 2 |
| MARE1 | [1Ac]-AVNVYSTSVTSDNLSR | Q15691 | 877.93 | 2 |
| MATR3 | DLSAAGIGLLAAATQSLSMPASLGR | P43243 | 791.09 | 3 |
| MATR3 | SQAFIEMETR | P43243 | 606.29 | 2 |
| MATR3 | GDADQASNILASFGLSAR | P43243 | 896.94 | 2 |
| MBB1A | VLDLVEVLVTK | Q9BQG0 | 614.38 | 2 |
| MBB1A | LLGAALPLLTK | Q9BQG0 | 555.37 | 2 |
| MCA3 | [1Ac]-AAAAELSLLEK | O43324 | 579.32 | 2 |
| MCM2 | ESLVVNYEDLAAR | P49736 | 739.88 | 2 |
| MCM3 | VQVVGTYR | P25205 | 461.26 | 2 |

| | | | | |
|---|---|---|---|---|
| MCM3 | VALLDVFR | P25205 | 466.78 | 2 |
| MCM3 | [1Ac]-AGTVVLDDVELR | P25205 | 664.86 | 2 |
| MCM4 | GILLQLFGGTR | P33991 | 587.85 | 2 |
| MCM4 | NLNPEDIDQLITISGMVIR | P33991 | 714.38 | 3 |
| MCM5 | IPGIIIAASAVR | P33992 | 590.87 | 2 |
| MCM5 | VLGIQVDTDGSGR | P33992 | 658.84 | 2 |
| MCM6 | IQETQAELPR | Q14566 | 592.82 | 2 |
| MCM6 | [1Ac]-MDLAAAAEPGAGSQHLEVR | Q14566 | 982.98 | 2 |
| MCM7 | IAQPGDHVSVTGIFLPILR | P33993 | 678.39 | 3 |
| MCM7 | GSSGVGLTAAVLR | P33993 | 594.34 | 2 |
| MCM7 | SLEQNIQLPAALLSR | P33993 | 826.97 | 2 |
| MDHC | DLDVAILVGSMPR | P40925 | 693.38 | 2 |
| MDHC | GEFVTTVQQR | P40925 | 582.80 | 2 |
| MDHC | FVEGLPINDFSR | P40925 | 697.36 | 2 |
| MDHM | IFGVTTLDIVR | P40926 | 617.36 | 2 |
| MDHM | IQEAGTEVVK | P40926 | 537.30 | 2 |
| MDHM | VAVLGASGGIGQPLSLLLK | P40926 | 897.05 | 2 |
| MDHM | AGAGSATLSMAYAGAR | P40926 | 727.86 | 2 |
| MDHM | LTLYDIAHTPGVAADLSHIETK | P40926 | 592.07 | 4 |
| MEP50 | SDGALLLGASSLSGR | Q9BQA1 | 702.38 | 2 |
| METK2 | FVIGGPQGDAGLTGR | P31153 | 722.88 | 2 |
| METK2 | TQVTVQYMQDR | P31153 | 684.83 | 2 |
| MFS10 | DAADLLSPLALLR | Q14728 | 684.40 | 2 |
| MGST3 | IASGLGLAWIVGR | O14880 | 656.89 | 2 |
| MGST3 | VLYAYGYYTGEPSK | O14880 | 805.89 | 2 |
| MIC19 | VAEELALEQAK | Q9NX63 | 600.83 | 2 |
| MIC60 | GIEQAVQSHAVAEEEAR | Q16891 | 608.63 | 3 |
| MIC60 | SEIQAEQDR | Q16891 | 538.25 | 2 |
| MICA1 | VAVELALLGAR | Q8TDZ2 | 556.35 | 2 |
| MICA1 | TVEETQVPEISGVAR | Q8TDZ2 | 807.92 | 2 |
| MK01 | [1Ac]-AAAAAAGAGPEMVR | P28482 | 642.82 | 2 |
| MK03 | [1Ac]-AAAAAQGGGGGEPR | P27361 | 606.29 | 2 |
| ML12B | FTDEEVDELYR | O14950 | 708.32 | 2 |
| MLEC | SNPEDQILYQTER | Q14165 | 796.88 | 2 |
| MOB1B | ELAPLQELIEK | Q7L9L4 | 641.87 | 2 |
| MOES | ISQLEMAR | P26038 | 474.25 | 2 |
| MOES | FYPEDVSEELIQDITQR | P26038 | 694.67 | 3 |
| MOES | EDAVLEYLK | P26038 | 540.28 | 2 |
| MOES | EVWFFGLQYQDTK | P26038 | 830.90 | 2 |
| MOES | TQEQLALEMAELTAR | P26038 | 852.44 | 2 |
| MOGS | LGPLLDILADSR | Q13724 | 641.87 | 2 |
| MPCP | IQTQPGYANTLR | Q00325 | 681.36 | 2 |
| MPCP | LPRPPPPEMPESLK | Q00325 | 529.96 | 3 |
| MPCP | FGFYEVFK | Q00325 | 518.76 | 2 |
| MPPA | GLDTVVALLADVVLQPR | Q10713 | 593.68 | 3 |
| MPPB | IDAVNAETIR | O75439 | 551.30 | 2 |
| MPRD | HTLADNFNPVSEER | P20645 | 543.59 | 3 |
| MPU1 | LLVPILLPEK | O75352 | 567.88 | 2 |
| MRP | GDVTAEEAAGASPAK | P49006 | 687.33 | 2 |
| MTCH2 | EEGILGFFAGLVPR | Q9Y6C9 | 752.91 | 2 |
| MTMRE | TPLFISLLR | Q8NCE2 | 530.33 | 2 |
| MVP | VPHNAAVQVYDYR | Q14764 | 511.26 | 3 |
| MVP | DAQGLVLFDVTGQVR | Q14764 | 809.43 | 2 |
| MVP | TAVFGFETSEAK | Q14764 | 643.82 | 2 |
| MVP | [1Ac]-ATEEFIIR | Q14764 | 510.77 | 2 |
| MYADM | ALTQPLGLLR | Q96S97 | 541.34 | 2 |
| MYH11 | TGVLAHLEEER | P35749 | 418.55 | 3 |
| MYH9 | VISGVLQLGNIVFK | P35579 | 743.95 | 2 |
| MYH9 | LQQELDDLLVDLDHQR | P35579 | 650.67 | 3 |
| MYH9 | AGVLAHLEEER | P35579 | 408.55 | 3 |
| MYH9 | VSHLLGINVTDFTR | P35579 | 524.62 | 3 |
| MYH9 | IIGLDQVAGMSETALPGAFK | P35579 | 673.36 | 3 |
| MYL6 | VLDFEHFLPMLQTVAK | P60660 | 630.01 | 3 |
| MYL6 | HVLVTLGEK | P60660 | 498.30 | 2 |
| MYL6 | DQGTYEDYVEGLR | P60660 | 772.85 | 2 |
| MYL9 | FTDEEVDEMYR | P24844 | 717.30 | 2 |
| MYO1C | TSFLLNLR | O00159 | 482.28 | 2 |
| MYO1F | LFDFLVEAINR | O00160 | 668.87 | 2 |
| MZB1 | ELSELVYTDVLDR | Q8WU39 | 776.40 | 2 |
| NAA15 | DLSLLQIQMR | Q9BXJ9 | 608.84 | 2 |
| NACA | DIELVMSQANVSR | Q13765 | 731.37 | 2 |
| NAGK | EGFLLALTQGR | Q9UJ70 | 602.84 | 2 |
| NAGK | SLGLSLSGGDQEDAGR | Q9UJ70 | 781.38 | 2 |
| NAGK | HIVAVLPEIDPVLFQGK | Q9UJ70 | 625.70 | 3 |

| | | | | |
|---|---|---|---|---|
| NAMPT | STQAPLIIRPDSGNPLDTVLK | P43490 | 745.75 | 3 |
| NB5R3 | STPAITLESPDIK | P00387 | 686.37 | 2 |
| NB5R3 | GPSGLLVYQGK | P00387 | 559.81 | 2 |
| NB5R3 | DILLRPELEELR | P00387 | 499.29 | 3 |
| NB5R3 | EIISHDTR | P00387 | 485.75 | 2 |
| NC2A | ALELFLESLLK | Q14919 | 638.38 | 2 |
| NCEH1 | IVQELPQLLDAR | Q6PIU2 | 697.90 | 2 |
| NCF1 | LLDGWWVIR | P14598 | 579.33 | 2 |
| NDE1 | ISALNIVGDLLR | Q9NXR1 | 642.39 | 2 |
| NDKA | NIIHGSDSVESAEK | P15531 | 495.91 | 3 |
| NDRG1 | TASGSSVTSLDGTR | Q92597 | 669.83 | 2 |
| NDUA9 | VFEISPFEPWITR | Q16795 | 810.92 | 2 |
| NDUA9 | LFLPFPLPLFAYR | Q16795 | 797.46 | 2 |
| NDUAC | [1Ac]-MELVQVLK | Q9UI09 | 501.29 | 2 |
| NDUB4 | GLIENPALLR | O95168 | 548.33 | 2 |
| NDUS1 | FASEIAGVDDLGTTGR | P28331 | 804.90 | 2 |
| NDUS2 | LVMELSGEMVR | O75306 | 632.33 | 2 |
| NDUS3 | VVAEPVELAQEFR | O75489 | 743.90 | 2 |
| NDUS3 | FEIVYNLLSLR | O75489 | 683.89 | 2 |
| NDUS8 | GLGMTLSYLFR | O00217 | 629.34 | 2 |
| NDUV2 | AAAVLPVLDLAQR | P19404 | 668.90 | 2 |
| NECP2 | AFIGIGFGDR | Q9NVZ3 | 526.78 | 2 |
| NEK9 | SSTVTEAPIAVVTSR | Q8TD19 | 759.41 | 2 |
| NEUL | LVNTGLLTLR | Q9BYT8 | 550.35 | 2 |
| NFKB1 | LMFTAFLPDSTGSFTR | P19838 | 895.94 | 2 |
| NH2L1 | QQIQSIQQSIER | P55769 | 729.39 | 2 |
| NH2L1 | [1Ac]-TEADVNPK | P55769 | 458.22 | 2 |
| NHRF1 | SVDPDSPAEASGLR | O14745 | 700.84 | 2 |
| NHRF1 | [1Ac]-SADAAAGAPLPR | O14745 | 569.80 | 2 |
| NID2 | VFALYNDEER | Q14112 | 628.30 | 2 |
| NIPA | FGMLPLDEPAILVSEFLDR | Q86WB0 | 721.38 | 3 |
| NIPS1 | MGPNIYELR | Q9BPW8 | 546.78 | 2 |
| NOLC1 | VVPSDLYPLVLGFLR | Q14978 | 844.49 | 2 |
| NONO | FAQPGSFEYEYAMR | Q15233 | 848.38 | 2 |
| NONO | NLPQYVSNELLEEAFSVFGQVER | Q15233 | 890.11 | 3 |
| NONO | MGQMAMGGAMGINNR | Q15233 | 769.84 | 2 |
| NOP2 | IQDIVGILR | P46087 | 513.82 | 2 |
| NOP56 | LSFYETGEIPR | O00567 | 656.33 | 2 |
| NOP58 | TQLYEYLQNR | Q9Y2X3 | 664.34 | 2 |
| NOP58 | YDAFGEDSSSAMGVENR | Q9Y2X3 | 917.88 | 2 |
| NP1L1 | LDGLVETPTGYIESLPR | P55209 | 930.49 | 2 |
| NP1L1 | GIPEFWLTVFK | P55209 | 668.87 | 2 |
| NP1L4 | VLAALQER | Q99733 | 450.27 | 2 |
| NPM | MTDQEAIQDLWQWR | P06748 | 910.43 | 2 |
| NPM | MSVQPTVSLGGFEITPPVVLR | P06748 | 743.08 | 3 |
| NPM | GPSSVEDIK | P06748 | 466.24 | 2 |
| NPM | DELHIVEAEAMNYEGSPIK | P06748 | 715.68 | 3 |
| NPM3 | [1Ac]-AAGTAAALAFLSQESR | O75607 | 803.42 | 2 |
| NSF | LLDYVPIGPR | P46459 | 571.83 | 2 |
| NSF1C | EFVAVTGAEEDR | Q9UNZ2 | 661.81 | 2 |
| NTM1A | LLLPLFR | Q9BV86 | 436.29 | 2 |
| NU160 | DLLILQQLLMR | Q12769 | 678.41 | 2 |
| NU160 | FVSSPQTIVELFFQEVAR | Q12769 | 699.71 | 3 |
| NU205 | YSFIQALVR | Q92621 | 548.81 | 2 |
| NU205 | MLALALLDR | Q92621 | 508.30 | 2 |
| NU5M | MILLTLTGQPR | P03915 | 621.87 | 2 |
| NUCB1 | DLELLIQTATR | Q02818 | 636.86 | 2 |
| NUCL | EVFEDAAEIR | P19338 | 589.79 | 2 |
| NUCL | NDLAVVDVR | P19338 | 500.77 | 2 |
| NUCL | TGISDVFAK | P19338 | 469.25 | 2 |
| NUCL | GLSEDTTEETLK | P19338 | 661.82 | 2 |
| NUCL | GFGFVDFNSEEDAK | P19338 | 781.34 | 2 |
| NUCL | [CRM]-VTQDELK[1Ac]EVFEDAAEIR | P19338 | 693.01 | 3 |
| NUCL | [1Ac]-VTQ[Dea]DELK[1Ac]EVFEDAAEIR | P19338 | 693.00 | 3 |
| NUP93 | NLQEIQQAGER | Q8N1F7 | 643.33 | 2 |
| ODO2 | GLVVPVIR | P36957 | 426.79 | 2 |
| ODO2 | NVEAMNFADIER | P36957 | 704.83 | 2 |
| ODPA | LEEGPPVTTVLTR | P08559 | 706.39 | 2 |
| ODPA | GPILMELQTYR | P08559 | 660.85 | 2 |
| ODPB | IMEGPAFNFLDAPAVR | P11177 | 874.45 | 2 |
| OST48 | SSLNPILFR | P39656 | 523.80 | 2 |
| OST48 | NTLLIAGLQAR | P39656 | 585.35 | 2 |
| OST48 | YSQTGNYELAVALSR | P39656 | 836.42 | 2 |
| OSTF1 | GYADIVQLLLAK | Q92882 | 652.38 | 2 |

| | | | | |
|---|---|---|---|---|
| OTUB1 | LLTSGYLQR | Q96FW1 | 525.80 | 2 |
| OTUB1 | IQQEIAVQNPLVSER | Q96FW1 | 862.47 | 2 |
| OXLA | LALDDVAALHGPVVR | Q96RQ9 | 515.96 | 3 |
| OXLA | IVGGWDLLPR | Q96RQ9 | 563.32 | 2 |
| OXLA | VIVVGAGVAGLVAAK | Q96RQ9 | 662.42 | 2 |
| OXSR1 | IPISLVLR | O95747 | 455.81 | 2 |
| P3H1 | DLSFFGGLLR | Q32P28 | 562.81 | 2 |
| P5CS | TPLFDQIIDMLR | P54886 | 731.39 | 2 |
| P5CS | MIDLIIPR | P54886 | 485.79 | 2 |
| PA1B2 | IIVLGLLPR | P68402 | 497.34 | 2 |
| PA1B3 | VVVLGLLPR | Q15102 | 483.33 | 2 |
| PA2G4 | ITSGPFEPDLYK | Q9UQ80 | 683.85 | 2 |
| PA2G4 | FDAMPFTLR | Q9UQ80 | 549.28 | 2 |
| PABP1 | FSPAGPILSIR | P11940 | 579.34 | 2 |
| PABP1 | GFGFVSFER | P11940 | 523.26 | 2 |
| PABP2 | TSLALDESLFR | Q86U42 | 626.33 | 2 |
| PABP2 | [1Ac]-AAAAAAAAAAGAAGGR | Q86U42 | 620.82 | 2 |
| PACN2 | AADAVEDLR | Q9UNF0 | 480.24 | 2 |
| PARK7 | GAEEMETVIPVDVMR | Q99497 | 838.41 | 2 |
| PARP1 | TTNFAGILSQGLR | P09874 | 689.38 | 2 |
| PARP1 | AEPVEVVAPR | P09874 | 533.80 | 2 |
| PARP1 | VVSEDFLQDVSASTK | P09874 | 812.91 | 2 |
| PARP1 | VGTVIGSNK | P09874 | 437.75 | 2 |
| PAXX | LAAAEETAVSPR | Q9BUH6 | 607.82 | 2 |
| PCBP1 | IITLTGPTNAIFK | Q15365 | 694.91 | 2 |
| PCBP1 | QGANINEIR | Q15365 | 507.77 | 2 |
| PCBP1 | IANPVEGSSGR | Q15365 | 543.78 | 2 |
| PCBP2 | IITLAGPTNAIFK | Q15366 | 679.91 | 2 |
| PCBP2 | IANPVEGSTDR | Q15366 | 579.79 | 2 |
| PDC10 | VNLSAAQTLR | Q9BUL8 | 536.81 | 2 |
| PDC6I | FYNELTEILVR | Q8WUM4 | 698.88 | 2 |
| PDC6I | LLDEEEATDNDLR | Q8WUM4 | 766.86 | 2 |
| PDCD4 | APQLVGQFIAR | Q53EL6 | 600.35 | 2 |
| PDCD6 | LSDQFHDILIR | O75340 | 452.91 | 3 |
| PDCD6 | SIISMFDR | O75340 | 484.75 | 2 |
| PDIA1 | LITLEEEMTK | P07237 | 603.82 | 2 |
| PDIA1 | EADDIVNWLK | P07237 | 601.81 | 2 |
| PDIA1 | ENLLDFIK | P07237 | 496.28 | 2 |
| PDIA1 | TVIDYNGER | P07237 | 533.76 | 2 |
| PDIA3 | ELSDFISYLQR | P30101 | 685.85 | 2 |
| PDIA3 | LAPEYEAAATR | P30101 | 596.30 | 2 |
| PDIA3 | TFSHELSDFGLESTAGEIPVVAIR | P30101 | 859.11 | 3 |
| PDIA3 | LNFAVASR | P30101 | 439.25 | 2 |
| PDIA3 | FVMQEEFSR | P30101 | 586.77 | 2 |
| PDIA3 | QAGPASVPLR | P30101 | 498.29 | 2 |
| PDIA4 | IDATSASVLASR | P13667 | 595.82 | 2 |
| PDIA6 | TGEAIVDAALSALR | Q15084 | 693.88 | 2 |
| PDIA6 | GSTAPVGGGAFPTIVER | Q15084 | 808.43 | 2 |
| PDIA6 | LAAVDATVNQVLASR | Q15084 | 764.43 | 2 |
| PDIA6 | GSFSEQGINEFLR | Q15084 | 742.36 | 2 |
| PDIA6 | ALDLFSDNAPPPELLEIINEDIAK | Q15084 | 879.79 | 3 |
| PDIA6 | NSYLEVLLK | Q15084 | 539.81 | 2 |
| PDLI1 | SAMPFTASPASSTTAR | O00151 | 791.88 | 2 |
| PDLI1 | [1Ac]-TTQQIDLQGPGPWGFR | O00151 | 921.96 | 2 |
| PDP1 | LLGLLMPFR | Q9P0J1 | 530.32 | 2 |
| PEA15 | RPDLLTMVVDYR | Q15121 | 493.27 | 3 |
| PEBB | AQQEDALAQQAFEEAR | Q13951 | 902.93 | 2 |
| PEBP1 | LYTLVLTDPDAPSR | P30086 | 780.92 | 2 |
| PEBP1 | YVWLVYEQDRPLK | P30086 | 570.31 | 3 |
| PEF1 | LSFEDFVTMTASR | Q9UBV8 | 752.36 | 2 |
| PEPD | VPLALFALNR | P12955 | 557.34 | 2 |
| PERM | VVLEGGIDPILR | P05164 | 640.88 | 2 |
| PFKAL | TNVLGHLQQGGAPTPFDR | P17858 | 636.66 | 3 |
| PFKAL | FDEATQLR | P17858 | 490.25 | 2 |
| PFKAL | SEWGSLLEELVAEGK | P17858 | 823.92 | 2 |
| PFKAL | ISMAAYVSGELEHVTR | P17858 | 588.30 | 3 |
| PGAM1 | ALPFWNEEIVPQIK | P18669 | 842.46 | 2 |
| PGAM1 | TLWTVLDAIDQMWLPVVR | P18669 | 719.39 | 3 |
| PGAM1 | FSGWYDADLSPAGHEEAK | P18669 | 660.63 | 3 |
| PGAM1 | HGESAWNLENR | P18669 | 438.21 | 3 |
| PGAM5 | AIETTDIISR | Q96HS1 | 559.81 | 2 |
| PGBM | LLSGPYFWSLPSR | P98160 | 761.91 | 2 |
| PGBM | LEGDTLIIPR | P98160 | 563.83 | 2 |
| PGBM | YELGSGLAVLR | P98160 | 589.33 | 2 |

| | | | | |
|---|---|---|---|---|
| PGK1 | ALESPERPFLAILGGAK | P00558 | 590.34 | 3 |
| PGK1 | VLPGVDALSNI | P00558 | 549.31 | 2 |
| PGK1 | YSLEPVAVELK | P00558 | 624.35 | 2 |
| PGK1 | LGDVYVNDAFGTAHR | P00558 | 545.60 | 3 |
| PGK1 | [CRM]-AK[1Ac]QIVWNGPVGVFEWEAFAR | P00558 | 797.07 | 3 |
| PGK1 | [1Ac]-AK[1Ac]QIVWN[Dea]GPVGVFEWEAFAR | P00558 | 797.07 | 3 |
| PGK1 | [1Ac]-AK[1Ac]Q[Dea]IVWN[Dea]GPVGVFEWEAFAR | P00558 | 797.40 | 3 |
| PGK1 | [1Ac]-AK[1Ac]Q[Dea]IVWNGPVGVFEWEAFAR | P00558 | 797.07 | 3 |
| PGM1 | LSGTGSAGATIR | P36871 | 545.80 | 2 |
| PGM1 | [1Ac]-GFARSM[Oxi]PTSGALDR | P36871 | 762.37 | 2 |
| PGM2 | IVLANDPDADR | Q96G03 | 599.81 | 2 |
| PGM2 | [1Ac]-AAPEGSGLGEDAR | Q96G03 | 636.30 | 2 |
| PGRC2 | GLGAGAGAGEESPATSLPR | O15173 | 849.43 | 2 |
| PGS1 | IQAIELEDLLR | P21810 | 656.88 | 2 |
| PHB | VLPSITTEILK | P35232 | 607.37 | 2 |
| PHB | FDAGELITQR | P35232 | 575.30 | 2 |
| PHB | IFTSIGEDYDER | P35232 | 722.83 | 2 |
| PHB | ILFRPVASQLPR | P35232 | 466.29 | 3 |
| PHB | NITYLPAGQSVLLQLPQ | P35232 | 928.02 | 2 |
| PHB2 | IGGVQQDTILAEGLHFR | Q99623 | 618.67 | 3 |
| PHB2 | VLPSIVNEVLK | Q99623 | 605.87 | 2 |
| PHB2 | LGLDYEER | Q99623 | 497.75 | 2 |
| PHB2 | VLSRPNAQELPSMYQR | Q99623 | 630.33 | 3 |
| PHB2 | ESVFTVEGGHR | Q99623 | 406.54 | 3 |
| PI42A | FLDFIGHILT | P48426 | 588.33 | 2 |
| PKHA2 | DNLFEIITSSR | Q9HB19 | 647.84 | 2 |
| PKN1 | SLGPVELLLR | Q16512 | 548.84 | 2 |
| PLCG2 | YPVTPELLER | P16885 | 608.83 | 2 |
| PLCG2 | EGSDSYAITFR | P16885 | 623.29 | 2 |
| PLCG2 | TGYVLQPESMR | P16885 | 640.82 | 2 |
| PLCG2 | VEELFEWFQSIR | P16885 | 791.90 | 2 |
| PLCG2 | MYVDPSEINPSMPQR | P16885 | 882.41 | 2 |
| PLD3 | AFLLSLAALR | Q8IV08 | 537.84 | 2 |
| PLEC | SLVPAAELLESR | Q15149 | 642.86 | 2 |
| PLEC | APVPASELLASGVLSR | Q15149 | 783.95 | 2 |
| PLEC | LTAEDLFEAR | Q15149 | 582.80 | 2 |
| PLEK | EDPAYLHYYDPAGAEDPLGAIHLR | P08567 | 671.58 | 4 |
| PLMN | FVTWIEGVMR | P00747 | 619.32 | 2 |
| PLP2 | HTAAPTDPADGPV | Q04941 | 624.80 | 2 |
| PLSL | NEALIALLR | P13796 | 506.81 | 2 |
| PLSL | QFVTATDVVR | P13796 | 568.31 | 2 |
| PLSL | ISFDEFIK | P13796 | 499.76 | 2 |
| PLSL | LNLAFIANLFNR | P13796 | 703.40 | 2 |
| PLSL | NWMNSLGVNPR | P13796 | 644.32 | 2 |
| PLSL | TLTLALIWQLMR | P13796 | 729.93 | 2 |
| PLSL | [1Ac]-GN[Dea]PK[1Ac]LNLAFIANLFNR | P13796 | 629.68 | 3 |
| PLSL | VNHLYSDLSDALVIFQLYEK[1Ac]IK[CRM] | P13796 | 674.11 | 4 |
| PLSL | [CRM]-EGK[1Ac]PYLVLGLLWQVIK | P13796 | 647.71 | 3 |
| PML | TGSALVQR | P29590 | 416.24 | 2 |
| PML | AETEELIR | P29590 | 480.75 | 2 |
| PMVK | EAYGAVTQTVR | Q15126 | 597.81 | 2 |
| PNPH | VFGFSLITNK | P00491 | 563.32 | 2 |
| PNPH | FEVGDIMLIR | P00491 | 596.82 | 2 |
| PNPH | LGADAVGMSTVPEVIVAR | P00491 | 892.98 | 2 |
| PNPH | DHINLPGFSGQNPLR | P00491 | 555.62 | 3 |
| PO210 | VGQALELPLR | Q8TEM1 | 548.33 | 2 |
| POSTN | AAAITSDILEALGR | Q15063 | 700.89 | 2 |
| POSTN | VLTQIGTSIQDFIEAEDDLSSFR | Q15063 | 862.10 | 3 |
| POSTN | FSTFLSLLEAADLK | Q15063 | 777.92 | 2 |
| PP1A | LNLDSIIGR | P62136 | 500.79 | 2 |
| PP1B | IVQMTEAEVR | P62140 | 588.31 | 2 |
| PP1R7 | SLETVVYLER | Q15435 | 555.30 | 2 |
| PP2AB | YSFLQFDPAPR | P62714 | 670.84 | 2 |
| PP2AB | ESNVQEVR | P62714 | 480.74 | 2 |
| PPIA | VSFELFADK | P62937 | 528.27 | 2 |
| PPIA | FEDENFILK | P62937 | 577.79 | 2 |
| PPIA | EGMNIVEAMER | P62937 | 639.79 | 2 |
| PPIA | [1Ac]-VNPTVFFDIAVDGEPLGR | P62937 | 994.51 | 2 |
| PPIA | [1Ac]-VNP[Oxi]T[Dhy]VFFDIAVDGEPLGR | P62937 | 993.50 | 2 |
| PPIA | [1Ac]-VN[Dea]PTVFFDIAVDGEPLGR | P62937 | 995.00 | 2 |
| PPIA | [1Ac]-VNPTVFFD[KXX]IAVDGEPLGR | P62937 | 675.99 | 3 |
| PPIA | VNPT[Pho]VFFDIAVDGEPLGR | P62937 | 675.99 | 3 |
| PPIA | [1Ac]-VN[Dea]P[Oxi]T[Dhy]VFFDIAVDGEPLGR | P62937 | 993.99 | 2 |
| PPIB | VLEGMEVVR | P23284 | 516.28 | 2 |

| | | | | |
|---|---|---|---|---|
| PPIB | TVDNFVALATGEK | P23284 | 682.86 | 2 |
| PPOX | SILLGLLLGAGR | P50336 | 591.88 | 2 |
| PRAF3 | TPMGIVLDALEQQEEGINR | O75915 | 705.02 | 3 |
| PRAF3 | AWDDFFPGSDR | O75915 | 656.78 | 2 |
| PRAF3 | [1Ac]-MDVNIAPLR | O75915 | 535.79 | 2 |
| PRDX1 | LVQAFQFTDK | Q06830 | 598.82 | 2 |
| PRDX1 | TIAQDYGVLK | Q06830 | 554.31 | 2 |
| PRDX1 | ADEGISFR | Q06830 | 447.72 | 2 |
| PRDX1 | ATAVMPDGQFK | Q06830 | 582.79 | 2 |
| PRDX1 | [1Ac]-SSGNAK[Frm]IGHPAPNFK | Q06830 | 797.90 | 2 |
| PRDX2 | EGGLGPLNIPLLADVTR | P32119 | 867.99 | 2 |
| PRDX2 | LSEDYGVLK | P32119 | 512.27 | 2 |
| PRDX2 | KEGGLGPLNIPLLADVTR | P32119 | 621.69 | 3 |
| PRDX2 | TDEGIAYR | P32119 | 462.72 | 2 |
| PRDX3 | DYGVLLEGSGLALR | P30048 | 731.90 | 2 |
| PRDX3 | GLFIIDPNGVIK | P30048 | 643.38 | 2 |
| PRDX3 | HLSVNDLPVGR | P30048 | 603.83 | 2 |
| PRDX3 | [1Ac]-HLSVN[Dea]DLPVGR | P30048 | 417.22 | 3 |
| PRDX4 | QITLNDLPVGR | Q13162 | 613.35 | 2 |
| PRDX5 | LLADPTGAFGK | P30044 | 545.30 | 2 |
| PRDX5 | ETDLLLDDSLVSIFGNR | P30044 | 953.99 | 2 |
| PRDX6 | FHDFLGDSWGILFSHPR | P30041 | 508.50 | 4 |
| PRDX6 | LPFPIIDDR | P30041 | 543.30 | 2 |
| PRDX6 | LSILYPATTGR | P30041 | 596.34 | 2 |
| PRDX6 | VVFVFGPDK | P30041 | 504.28 | 2 |
| PRDX6 | ELAILLGMLDPAEKDEK | P30041 | 629.00 | 3 |
| PRDX6 | VVISLQLTAEK | P30041 | 600.86 | 2 |
| PRKDC | VTELALTASDR | P78527 | 588.32 | 2 |
| PRKDC | LLEEALLR | P78527 | 478.79 | 2 |
| PRKDC | VTVMASLR | P78527 | 438.75 | 2 |
| PRKDC | [Frm]-YK[1Ac]EVYAAAAEVLGLILR | P78527 | 650.36 | 3 |
| PROF1 | TFVNITPAEVGVLVGK | P07737 | 822.47 | 2 |
| PROF1 | DSLLQDGEFSMDLR | P07737 | 813.38 | 2 |
| PROF1 | DSPSVWAAVPGK | P07737 | 607.31 | 2 |
| PROF1 | TLVLLMGK | P07737 | 437.78 | 2 |
| PRP16 | SLNTDVLFGLLR | Q92620 | 674.39 | 2 |
| PRS10 | VALDMTTLTIMR | P62333 | 682.87 | 2 |
| PRS10 | EVIELPLTNPELFQR | P62333 | 899.49 | 2 |
| PRS6A | VDILDPALLR | P17980 | 562.84 | 2 |
| PRS6B | [1Ac]-MEEIGILVEK | P43686 | 601.82 | 2 |
| PRS7 | FDDGAGGDNEVQR | P35998 | 690.29 | 2 |
| PRS8 | IDILDSALLRPGR | P62195 | 480.28 | 3 |
| PRS8 | EHAPSIIFMDEIDSIGSSR | P62195 | 702.01 | 3 |
| PRS8 | LEGGSGGDSEVQR | P62195 | 645.80 | 2 |
| PSA | LGLQNDLFSLAR | P55786 | 673.88 | 2 |
| PSA | VALSNMNVIDR | P55786 | 616.33 | 2 |
| PSA | YAAVTQFEATDAR | P55786 | 721.85 | 2 |
| PSA | VLGATLLPDLIQK | P55786 | 690.93 | 2 |
| PSA1 | ETLPAEQDLTTK | P25786 | 673.35 | 2 |
| PSA2 | YNEDLELEDAIHTAILTLK | P25787 | 734.38 | 3 |
| PSA2 | GYSFSLTTFSPSGK | P25787 | 739.86 | 2 |
| PSA2 | LAQQYYLVYQEPIPTAQLVQR | P25787 | 841.12 | 3 |
| PSA2 | HIGLVYSGMGPDYR | P25787 | 522.26 | 3 |
| PSA3 | HVGMAVAGLLADAR | P25788 | 460.92 | 3 |
| PSA3 | AVENSSTAIGIR | P25788 | 609.33 | 2 |
| PSA4 | LLDEVFFSEK | P25789 | 613.82 | 2 |
| PSA5 | GVNTFSPEGR | P28066 | 532.26 | 2 |
| PSA5 | LFQVEYAIEAIK | P28066 | 712.40 | 2 |
| PSA5 | ITSPLMEPSSIEK | P28066 | 716.37 | 2 |
| PSA6 | HITIFSPEGR | P60900 | 578.81 | 2 |
| PSA6 | ILTEAEIDAHLVALAERD | P60900 | 660.35 | 3 |
| PSA6 | AINQGGLTSVAVR | P60900 | 643.36 | 2 |
| PSA6 | LYQVEYAFK | P60900 | 580.80 | 2 |
| PSA7 | LTVEDPVTVEYITR | O14818 | 817.94 | 2 |
| PSB1 | DVFISAAER | P20618 | 504.26 | 2 |
| PSB1 | AMTTGAIAAMLSTILYSR | P20618 | 624.33 | 3 |
| PSB1 | GAVYSFDPVGSYQR | P20618 | 773.37 | 2 |
| PSB10 | LPFTALGSGQDAALAVLEDR | P40306 | 682.03 | 3 |
| PSB2 | FILNLPTFSVR | P49721 | 653.88 | 2 |
| PSB2 | VAASNIVQMK | P49721 | 530.79 | 2 |
| PSB2 | NGYELSPTAAANFTR | P49721 | 806.39 | 2 |
| PSB2 | [1Ac]-MEYLIGIQGPDYVLVASDR | P49721 | 727.70 | 3 |
| PSB2 | [1Ac]-MEYLIGIQ[Dea]GPDYVLVASDR | P49721 | 728.03 | 3 |
| PSB3 | LYIGLAGLATDVQTVAQR | P49720 | 630.35 | 3 |

| | | | | |
|---|---|---|---|---|
| PSB3 | FGPYYTEPVIAGLDPK | P49720 | 883.95 | 2 |
| PSB3 | DAVSGMGVIVHIIEK | P49720 | 523.29 | 3 |
| PSB6 | LAAIAESGVER | P28072 | 558.31 | 2 |
| PSB8 | ASAGSYISALR | P28062 | 548.29 | 2 |
| PSB9 | VSAGEAVVNR | P28065 | 501.27 | 2 |
| PSB9 | FTTDAIALAMSR | P28065 | 648.83 | 2 |
| PSB9 | EGGQVYGTLGGMLTR | P28065 | 769.89 | 2 |
| PSD11 | LYDNLLEQNLIR | O00231 | 752.41 | 2 |
| PSD12 | LQEVIETLLSLEK | O00232 | 757.94 | 2 |
| PSD12 | VEFILEQMR | O00232 | 582.81 | 2 |
| PSD12 | WSTLVEDYGMELR | O00232 | 799.88 | 2 |
| PSD12 | TASDMVSTSR | O00232 | 527.75 | 2 |
| PSD13 | VNPLSLVEIILHVVR | Q9UNM6 | 567.69 | 3 |
| PSD13 | LYENFISEFEHR | Q9UNM6 | 528.59 | 3 |
| PSMD1 | AAVESLGFILFR | Q99460 | 661.88 | 2 |
| PSMD2 | MLVTFDEELRPLPVSVR | Q13200 | 667.70 | 3 |
| PSMD2 | AVPLALALISVSNPR | Q13200 | 760.96 | 2 |
| PSMD2 | AELATEEFLPVTPILEGFVILR | Q13200 | 819.79 | 3 |
| PSMD3 | HDADGQATLLNLLLR | O43242 | 550.64 | 3 |
| PSMD5 | [1Ac]-AAQALALLR | Q16401 | 484.80 | 2 |
| PSMD6 | IGLFYMDNDLITR | Q15008 | 785.90 | 2 |
| PSMD6 | GAEILEVLHSLPAVR | Q15008 | 535.31 | 3 |
| PSMD6 | LDIVFYLLR | Q15008 | 576.34 | 2 |
| PSMD6 | IHAYSQLLESYR | Q15008 | 493.92 | 3 |
| PSMD9 | SDVDLYQVR | O00233 | 547.78 | 2 |
| PSME1 | NAYAVLYDIILK | Q06323 | 698.40 | 2 |
| PSME1 | DVIEQLNLVTTWLQLQIPR | Q06323 | 760.43 | 3 |
| PSME1 | IEDGNNFGVAVQEK | Q06323 | 760.37 | 2 |
| PSME2 | QNLFQEAEEFLYR | Q9UL46 | 843.91 | 2 |
| PSME2 | AFYAELYHIISSNLEK | Q9UL46 | 633.33 | 3 |
| PSME2 | DEAAYGELR | Q9UL46 | 512.24 | 2 |
| PSME2 | IIYLNQLLQEDSLNVADLTSLR | Q9UL46 | 844.46 | 3 |
| PSME3 | TVESEAASYLDQISR | P61289 | 834.91 | 2 |
| PSME3 | MWVQLLIPR | P61289 | 578.34 | 2 |
| PTBP1 | IAIPGLAGAGNSVLLVSNLNPER | P26599 | 759.10 | 3 |
| PTBP1 | KLPIDVTEGEVISLGLPFGK | P26599 | 704.74 | 3 |
| PTBP1 | IIVENLFYPVTLDVLHQIFSK | P26599 | 830.13 | 3 |
| PTBP1 | VTPQSLFILFGVYGDVQR | P26599 | 680.37 | 3 |
| PTCA | AELGSTDNDLER | Q14761 | 660.31 | 2 |
| PTMA | AAEDDEDDDVDTK | P06454 | 719.28 | 2 |
| PTMA | [1Ac]-SDAAVDTSSEITTK | P06454 | 733.85 | 2 |
| PTN1 | MGLIQTADQLR | P18031 | 623.33 | 2 |
| PTN1 | FSYLAVIEGAK | P18031 | 599.33 | 2 |
| PTN1 | QLELENLTTQETR | P18031 | 787.90 | 2 |
| PTN6 | YTVGGLETFDSLTDLVEHFK | P29350 | 757.71 | 3 |
| PTN6 | DLSGLDAETLLK | P29350 | 637.85 | 2 |
| PTN6 | TLQVSPLDNGDLIR | P29350 | 770.92 | 2 |
| PTPA | LVALLNTLDR | Q15257 | 564.34 | 2 |
| PTPA | WIDETPPVDQPSR | Q15257 | 770.38 | 2 |
| PTPRC | LFLAEFQSIPR | P08575 | 660.87 | 2 |
| PTPRC | YVDILPYDYNR | P08575 | 715.85 | 2 |
| PTPRC | DPPSEPSPLEAEFQR | P08575 | 849.90 | 2 |
| PTPRC | SEAAHQGVITWNPPQR | P08575 | 597.64 | 3 |
| PTSS1 | ILFIGGITAPTVR | P48651 | 679.41 | 2 |
| PUF60 | VYVGSIYYELGEDTIR | Q9UHX1 | 938.97 | 2 |
| PUR2 | AIAFLQQPR | P22102 | 522.30 | 2 |
| PUR2 | ENLISALEEAK | P22102 | 608.82 | 2 |
| PUR6 | ASILNTWISLK | P22234 | 623.36 | 2 |
| PUR6 | [1Ac]-ATAEVLNIGK | P22234 | 529.30 | 2 |
| PUR8 | NALDLLLPK | P30566 | 498.81 | 2 |
| PUR9 | DVSELTGFPEMLGGR | P31939 | 804.39 | 2 |
| PUR9 | TGLVEFAR | P31939 | 446.75 | 2 |
| PYGB | GLAGLGDVAEVR | P11216 | 578.82 | 2 |
| PYR1 | LALGIPLPELR | P27708 | 596.38 | 2 |
| PYR1 | MALLATVLGR | P27708 | 522.82 | 2 |
| PYR1 | TLGVDLVALATR | P27708 | 614.87 | 2 |
| PYR1 | VLGTSPEAIDSAENR | P27708 | 779.89 | 2 |
| PYR1 | [1Ac]-AALVLEDGSVLR | P27708 | 642.86 | 2 |
| QCR1 | SGMFWLR | P31930 | 448.73 | 2 |
| QCR2 | YEDFSNLGTTHLLR | P22695 | 555.95 | 3 |
| RAB10 | AFLTLAEDILR | P61026 | 631.36 | 2 |
| RAB14 | GAAGALMVYDITR | P61106 | 669.35 | 2 |
| RAB14 | LTSEPQPQR | P61106 | 528.28 | 2 |
| RAB18 | GAQGVILVYDVTR | Q9NP72 | 695.89 | 2 |

| | | | | |
|---|---|---|---|---|
| RAB1A | MGPGATAGGAEK | P62820 | 523.75 | 2 |
| RAB1B | MGPGAASGGERPNLK | Q9H0U4 | 481.25 | 3 |
| RAB21 | VNLAIWDTAGQER | Q9UL25 | 736.88 | 2 |
| RAB21 | MIETAQVDER | Q9UL25 | 596.29 | 2 |
| RAB21 | FHALGPIYYR | Q9UL25 | 412.89 | 3 |
| RAB21 | [1Ac]-AAAGGGGGGAAAAGR | Q9UL25 | 557.27 | 2 |
| RAB2A | GAAGALLVYDITR | P61019 | 660.37 | 2 |
| RAB2A | FQPVHDLTIGVEFGAR | P61019 | 595.98 | 3 |
| RAB2A | DTFNHLTTWLEDAR | P61019 | 573.61 | 3 |
| RAB4B | GAAGALLVYDITSR | P61018 | 703.89 | 2 |
| RAB5A | GVDLTEPTQPTR | P20339 | 657.34 | 2 |
| RAB5C | GVDLQENNPASR | P51148 | 650.32 | 2 |
| RAB5C | GAQAAIVVYDITNTDTFAR | P51148 | 676.01 | 3 |
| RAB7A | FQSLGVAFYR | P51149 | 594.31 | 2 |
| RAB7A | LVTMQIWDTAGQER | P51149 | 824.41 | 2 |
| RAB7A | DEFLIQASPR | P51149 | 588.31 | 2 |
| RAB7A | DPENFPFVVLGNK | P51149 | 738.38 | 2 |
| RAB7A | EAINVEQAFQTIAR | P51149 | 795.42 | 2 |
| RAB7L | LQLWDIAGQER | O14966 | 664.85 | 2 |
| RAB8A | ANINVENAFFTLAR | P61006 | 790.41 | 2 |
| RAB8B | NIEEHASSDVER | Q92930 | 693.32 | 2 |
| RAC2 | LAPITYPQGLALAK | P15153 | 728.43 | 2 |
| RAGP1 | LEGNTVGVEAAR | P46060 | 608.32 | 2 |
| RALY | VFIGNLNTALVK | Q9UKM9 | 644.88 | 2 |
| RALY | SNIDALLSR | Q9UKM9 | 494.77 | 2 |
| RALY | STAVTTSSAK | Q9UKM9 | 476.75 | 2 |
| RAN | SNYNFEKPFLWLAR | P62826 | 595.64 | 3 |
| RAN | HLTGEFEK | P62826 | 480.74 | 2 |
| RAN | YVATLGVEVHPLVFHTNR | P62826 | 513.78 | 4 |
| RAN | [1Ac]-AAQGEPQVQFK | P62826 | 622.82 | 2 |
| RAP2B | ASVDELFAEIVR | P61225 | 674.86 | 2 |
| RASL3 | VALALEELDAPR | Q86YV0 | 648.86 | 2 |
| RASL3 | AQALVTDLGTAELAR | Q86YV0 | 764.92 | 2 |
| RB11B | GAVGALLVYDIAK | Q15907 | 645.38 | 2 |
| RB11B | STIGVEFATR | Q15907 | 540.79 | 2 |
| RB11B | AQIWDTAGQER | Q15907 | 637.81 | 2 |
| RB11B | DHADSNIVIMLVGNK | Q15907 | 542.62 | 3 |
| RB11B | [1Ac]-GTRDDEYDYLFK | Q15907 | 782.35 | 2 |
| RBBP7 | TVALWDLR | Q16576 | 487.28 | 2 |
| RBBP7 | LMIWDTR | Q16576 | 467.74 | 2 |
| RBGPR | VILLDVAR | Q9H2M9 | 449.79 | 2 |
| RBM3 | GFGFITFTNPEHASVAMR | P98179 | 661.32 | 3 |
| RBM39 | DLEEFFSTVGK | Q14498 | 636.31 | 2 |
| RBM39 | LYVGSLHFNITEDMLR | Q14498 | 636.66 | 3 |
| RBM39 | TDASSASSFLDSDELER | Q14498 | 915.41 | 2 |
| RBMX | IVEVLLMK | P38159 | 472.80 | 2 |
| RBMX | LFIGGLNTETNEK | P38159 | 718.38 | 2 |
| RBMX | GFAFVTFESPADAK | P38159 | 743.86 | 2 |
| RBMX | YDDYSSSR | P38159 | 496.70 | 2 |
| RBMX | DSYESYGNSR | P38159 | 589.24 | 2 |
| RCC1 | VFLWGSFR | P18754 | 506.27 | 2 |
| RCC2 | VFSWGFGGYGR | Q9P258 | 616.80 | 2 |
| RCC2 | LFDFPGR | Q9P258 | 426.22 | 2 |
| RD23B | AVEYLLMGIPGDR | P54727 | 717.38 | 2 |
| RDH14 | [1Ac]-AVATAAAVLAALGGALWLAAR | Q9HBH5 | 990.08 | 2 |
| RECQ1 | VAGVVAPTLPR | P46063 | 540.33 | 2 |
| RFA3 | [1Ac]-VDMMDLPR | P35244 | 509.74 | 2 |
| RFTN1 | FLEFTTLSAAELPGSSAVR | Q14699 | 998.52 | 2 |
| RFTN1 | NQSPEPSSGPR | Q14699 | 578.27 | 2 |
| RHG01 | VPATLQVLQTLPEENYQVLR | Q07960 | 771.09 | 3 |
| RHG01 | FLLDHQGELFPSPDPSGL | Q07960 | 984.99 | 2 |
| RHG04 | LPAPVLVVLR | P98171 | 538.86 | 2 |
| RIC8A | LLFLLTALR | Q9NPQ8 | 530.35 | 2 |
| RINI | ELSLAGNELGDEGAR | P13489 | 765.87 | 2 |
| RIR2B | [1Ac]-GDPERPEAAGLDQDER | Q7LG56 | 898.91 | 2 |
| RL10 | VHIGQVIMSIR | P27635 | 418.24 | 3 |
| RL10 | FNADEFEDMVAEK | P27635 | 772.83 | 2 |
| RL10A | AVDIPHMDIEALK | P62906 | 484.59 | 3 |
| RL10A | YDAFLASESLIK | P62906 | 678.86 | 2 |
| RL11 | VLEQLTGQTPVFSK | P62913 | 773.93 | 2 |
| RL11 | [1Ac]-AQDQGEKENPMR | P62913 | 722.83 | 2 |
| RL12 | IGPLGLSPK | P30050 | 441.28 | 2 |
| RL12 | QAQIEVVPSASALIIK | P30050 | 833.99 | 2 |
| RL13 | LATQLTGPVMPVR | P26373 | 691.89 | 2 |

| | | | | |
|---|---|---|---|---|
| RL13 | TIGISVDPR | P26373 | 479.27 | 2 |
| RL13 | STESLQANVQR | P26373 | 616.82 | 2 |
| RL13A | MVVPAALK | P40429 | 414.75 | 2 |
| RL13A | YQAVTATLEEK | P40429 | 626.82 | 2 |
| RL13A | [1Ac]-AEVQVLVLDGR | P40429 | 620.85 | 2 |
| RL14 | LVAIVDVIDQNR | P50914 | 677.89 | 2 |
| RL15 | FFEVILIDPFHK | P61313 | 502.28 | 3 |
| RL15 | VLNSYWVGEDSTYK | P61313 | 830.90 | 2 |
| RL18 | ILTFDQLALDSPK | Q07020 | 730.90 | 2 |
| RL18 | TAVVVGTITDDVR | Q07020 | 673.37 | 2 |
| RL18A | NFGIWLR | Q02543 | 453.25 | 2 |
| RL18A | FWYFVSQLK | Q02543 | 609.32 | 2 |
| RL19 | LLADQAEAR | P84098 | 493.77 | 2 |
| RL1D1 | LLSSFDFFLTDAR | O76021 | 766.39 | 2 |
| RL21 | VYNVTQHAVGIVVNK | P46778 | 547.64 | 3 |
| RL21 | HGVVPLATYMR | P46778 | 622.33 | 2 |
| RL22 | AGNLGGGVVTIER | P35268 | 621.84 | 2 |
| RL22 | ITVTSEVPFSK | P35268 | 604.33 | 2 |
| RL23 | LPAAGVGDMVMATVK | P62829 | 487.26 | 3 |
| RL23A | LAPDYDALDVANK | P62750 | 702.85 | 2 |
| RL24 | AITGASLADIMAK | P83731 | 631.34 | 2 |
| RL24 | QINWTVLYR | P83731 | 596.83 | 2 |
| RL26 | DDEVQVVR | P61254 | 480.24 | 2 |
| RL27 | YSVDIPLDK | P61353 | 525.28 | 2 |
| RL27A | TGAAPIIDVVR | P46776 | 556.33 | 2 |
| RL27A | LWTLVSEQTR | P46776 | 616.84 | 2 |
| RL29 | AQAAAPASVPAQAPK | P47914 | 689.38 | 2 |
| RL29 | LAYIAHPK | P47914 | 456.77 | 2 |
| RL31 | LYTLVTYVPVTTFK | P62899 | 822.97 | 2 |
| RL31 | SAINEVVTR | P62899 | 494.77 | 2 |
| RL35 | VLTVINQTQK | P42766 | 572.34 | 2 |
| RL35A | DETEFYLGK | P18077 | 551.26 | 2 |
| RL36 | EELSNVLAAMR | Q9Y3U8 | 616.82 | 2 |
| RL38 | IEEIKDFLLTAR | P63173 | 483.28 | 3 |
| RL4 | APIRPDIVNFVHTNLR | P36578 | 466.27 | 4 |
| RL4 | AAAAAAALQAK | P36578 | 478.78 | 2 |
| RL4 | MINTDLSR | P36578 | 475.24 | 2 |
| RL6 | ASITPGTILIILTGR | Q02878 | 763.47 | 2 |
| RL6 | AIPQLQGYLR | Q02878 | 579.84 | 2 |
| RL6 | QLASGLLLVTGPLVLNR | Q02878 | 882.54 | 2 |
| RL6 | HQEGEIFDTEK | Q02878 | 666.81 | 2 |
| RL6 | [CRM]-VVFLK[1Ac]QLASGLLLVTGPLVLNR | Q02878 | 812.50 | 3 |
| RL7 | IALTDNALIAR | P18124 | 585.85 | 2 |
| RL7A | AGVNTVTTLVENK | P62424 | 673.37 | 2 |
| RL7A | NFGIGQDIQPK | P62424 | 608.82 | 2 |
| RL7A | VAPAPAVVK | P62424 | 426.27 | 2 |
| RL7A | LKVPPAINQFTQALDR | P62424 | 604.34 | 3 |
| RL8 | AVVGVVAGGGR | P62917 | 471.28 | 2 |
| RL8 | ASGNYATVISHNPETK | P62917 | 563.61 | 3 |
| RL9 | DFNHINVELSLLGK | P32969 | 533.62 | 3 |
| RLA0 | GTIEILSDVQLIK | P05388 | 714.92 | 2 |
| RLA0 | TSFFQALGITTK | P05388 | 657.36 | 2 |
| RLA0 | IIQLLDDYPK | P05388 | 609.34 | 2 |
| RLA0 | GNVGFVFTK | P05388 | 484.76 | 2 |
| RLA0 | VLALSVETDYTFPLAEK | P05388 | 948.50 | 2 |
| RLA1 | AAGVNVEPFWPGLFAK | P05386 | 851.95 | 2 |
| RLA2 | ILDSVGIEADDDR | P05387 | 709.34 | 2 |
| RM47 | FFALPYVDHFLR | Q9HD33 | 508.94 | 3 |
| RO60 | ALLQEMPLTALLR | P10155 | 734.93 | 2 |
| ROA0 | LFIGGLNVQTSESGLR | Q13151 | 845.96 | 2 |
| ROA1 | IEVIEIMTDR | P09651 | 609.82 | 2 |
| ROA1 | LFIGGLSFETTDESLR | P09651 | 892.96 | 2 |
| ROA1 | GFAFVTFDDHDSVDK | P09651 | 567.26 | 3 |
| ROA1 | EDSQRPGAHLTVK | P09651 | 479.92 | 3 |
| ROA2 | GGGGNFGPGPGSNFR | P22626 | 689.32 | 2 |
| ROA2 | GFGFVTFDDHDPVDK | P22626 | 565.93 | 3 |
| ROA2 | QEMQEVQSSR | P22626 | 611.28 | 2 |
| ROA2 | TLETVPLER | P22626 | 529.30 | 2 |
| ROA2 | NMGGPYGGGNYGPGGSGGSGGYGGR | P22626 | 730.64 | 3 |
| ROA3 | IETIEVMEDR | P51991 | 617.80 | 2 |
| ROA3 | LFIGGLSFETTDDSLR | P51991 | 885.95 | 2 |
| ROA3 | GFAFVTFDDHDTVDK | P51991 | 571.93 | 3 |
| ROA3 | EDTEEYNLR | P51991 | 584.76 | 2 |
| ROA3 | SSGSPYGGGYGSGGGSGGYGSR | P51991 | 955.90 | 2 |

| | | | | |
|---|---|---|---|---|
| ROAA | GFGFILFK | Q99729 | 464.77 | 2 |
| ROAA | GFVFITFK | Q99729 | 479.77 | 2 |
| ROAA | EVYQQQQYGSGGR | Q99729 | 750.35 | 2 |
| RPAB3 | IEGDETSTEAATR | P52434 | 690.32 | 2 |
| RPAB3 | [1Ac]-AGILFEDIFDVK | P52434 | 704.87 | 2 |
| RPF2 | SLLIDFFR | Q9H7B2 | 505.79 | 2 |
| RPIA | FIVIADFR | P49247 | 490.78 | 2 |
| RPN1 | ATSFLLALEPELEAR | P04843 | 830.45 | 2 |
| RPN1 | SEDLLDYGPFR | P04843 | 656.31 | 2 |
| RPN1 | VTAEVVLAHLGGGSTSR | P04843 | 551.97 | 3 |
| RPN1 | ALTSEIALLQSR | P04843 | 651.37 | 2 |
| RPN1 | TILPAAAQDVYYR | P04843 | 740.89 | 2 |
| RPN2 | SIVEEIEDLVAR | P04844 | 686.87 | 2 |
| RPN2 | YIANTVELR | P04844 | 539.80 | 2 |
| RPN2 | YHVPVVVVPEGSASDTHEQAILR | P04844 | 626.58 | 4 |
| RRBP1 | SIEALLEAGQAR | Q9P2E9 | 629.34 | 2 |
| RRBP1 | EQEITAVQAR | Q9P2E9 | 572.80 | 2 |
| RRP44 | NVIVLQTVLQEVR | Q9Y2L1 | 755.95 | 2 |
| RS10 | IAIYELLFK | P46783 | 555.33 | 2 |
| RS10 | DYLHLPPEIVPATLR | P46783 | 578.66 | 3 |
| RS10 | AEAGAGSATEFQFR | P46783 | 721.34 | 2 |
| RS10 | HFYWYLTNEGIQYLR | P46783 | 668.33 | 3 |
| RS12 | [1Ac]-AEEGIAAGGVM[Oxi]DVNTALQEVLK | P25398 | 758.38 | 3 |
| RS13 | GLTPSQIGVILR | P62277 | 627.38 | 2 |
| RS13 | GLAPDLPEDLYHLIK | P62277 | 565.31 | 3 |
| RS13 | DSHGVAQVR | P62277 | 484.75 | 2 |
| RS14 | IEDVTPIPSDSTR | P62263 | 715.36 | 2 |
| RS14 | TPGPGAQSALR | P62263 | 527.79 | 2 |
| RS14 | ADRDESSPYAAMLAAQDVAQR | P62263 | 755.69 | 3 |
| RS15 | GVDLDQLLDMSYEQLMQLYSAR | P62841 | 863.42 | 3 |
| RS15A | MNVLADALK | P62244 | 487.77 | 2 |
| RS15A | HGYIGEFEIIDDHR | P62244 | 567.61 | 3 |
| RS15A | QFGFIVLTTSAGIMDHEEAR | P62244 | 741.37 | 3 |
| RS16 | LLEPVLLLGK | P62249 | 547.86 | 2 |
| RS16 | TLLVADPR | P62249 | 442.76 | 2 |
| RS16 | GPLQSVQVFGR | P62249 | 594.33 | 2 |
| RS16 | GGGHVAQIYAIR | P62249 | 414.56 | 3 |
| RS18 | VITIMQNPR | P62269 | 536.30 | 2 |
| RS18 | AGELTEDEVER | P62269 | 624.29 | 2 |
| RS18 | IPDWFLNR | P62269 | 530.78 | 2 |
| RS18 | IAFAITAIK | P62269 | 474.30 | 2 |
| RS18 | YAHVVLR | P62269 | 429.25 | 2 |
| RS19 | IAGQVAAANK | P39019 | 471.77 | 2 |
| RS19 | ELAPYDENWFYTR | P39019 | 852.39 | 2 |
| RS2 | SLEEIYLFSLPIK | P15880 | 776.44 | 2 |
| RS2 | GTGIVSAPVPK | P15880 | 513.30 | 2 |
| RS20 | LIDLHSPSEIVK | P60866 | 450.93 | 3 |
| RS20 | TPVEPEVAIHR | P60866 | 416.56 | 3 |
| RS21 | MGESDDSILR | P63220 | 561.76 | 2 |
| RS21 | [1Ac]-MQNDAGEFVDLYVPR | P63220 | 898.42 | 2 |
| RS21 | [1Ac]-MQN[Dea]DAGEFVDLYVPR | P63220 | 898.91 | 2 |
| RS21 | [1Ac]-MQ[Dea]NDAGEFVDLYVPR | P63220 | 898.91 | 2 |
| RS23 | VANVSLLALYK | P62266 | 595.86 | 2 |
| RS23 | KGHAVGDIPGVR | P62266 | 402.56 | 3 |
| RS24 | TTPDVIFVFGFR | P62847 | 699.87 | 2 |
| RS24 | [1Ac]-MNDTVTIR | P62847 | 496.25 | 2 |
| RS25 | AALQELLSK | P62851 | 486.79 | 2 |
| RS26 | NIVEAAAVR | P62854 | 471.77 | 2 |
| RS26 | DISEASVFDAYVLPK | P62854 | 827.42 | 2 |
| RS28 | EGDVLTLLESER | P62857 | 680.85 | 2 |
| RS3 | TEIIILATR | P23396 | 515.32 | 2 |
| RS3 | AELNEFLTR | P23396 | 546.79 | 2 |
| RS3 | FGFPEGSVELYAEK | P23396 | 786.88 | 2 |
| RS3 | IMLPWDPTGK | P23396 | 579.30 | 2 |
| RS3 | ELAEDGYSGVEVR | P23396 | 712.34 | 2 |
| RS30 | FVNVVPTFGK | P62861 | 554.31 | 2 |
| RS3A | VFEVSLADLQNDEVAFR | P61247 | 976.49 | 2 |
| RS3A | ADGYEPPVQESV | P61247 | 645.80 | 2 |
| RS4X | LSNIFVIGK | P62701 | 495.80 | 2 |
| RS5 | [1Ac]-TEWETAAPAVAETPDIK | P46782 | 935.96 | 2 |
| RS6 | DIPGLTDTTVPR | P62753 | 642.84 | 2 |
| RS6 | MATEVAADALGEEWK | P62753 | 810.88 | 2 |
| RS7 | TLTAVHDAILEDLVFPSEIVGK | P62081 | 789.77 | 3 |
| RS7 | AIIIFVPVPQLK | P62081 | 669.43 | 2 |

| | | | | |
|---|---|---|---|---|
| RS7 | DVNFEFPEFQL | P62081 | 692.82 | 2 |
| RS8 | IIDVVYNASNNELVR | P62241 | 859.96 | 2 |
| RS8 | ADGYVLEGK | P62241 | 476.24 | 2 |
| RS9 | LFEGNALLR | P46781 | 516.80 | 2 |
| RS9 | IEDFLER | P46781 | 461.24 | 2 |
| RS9 | QVVNIPSFIVR | P46781 | 636.38 | 2 |
| RS9 | LDYILGLK | P46781 | 467.78 | 2 |
| RSMB | VLGLVLLR | P14678 | 441.81 | 2 |
| RSSA | AIVAIENPADVSVISSR | P08865 | 870.98 | 2 |
| RSSA | LLVVTDPR | P08865 | 456.78 | 2 |
| RSSA | FTPGTFTNQIQAAFR | P08865 | 849.93 | 2 |
| RSSA | FAAATGATPIAGR | P08865 | 602.33 | 2 |
| RSSA | [1Ac]-SGALDVLQMK | P08865 | 552.29 | 2 |
| RTN4 | GPLPAAPPVAPER | Q9NQC3 | 636.36 | 2 |
| RU17 | T[Pho]QFLPP[Oxi]NLLALFAPR | P08621 | 598.65 | 3 |
| RU2A | NAIANASTLAEVER | P09661 | 729.88 | 2 |
| RU2A | SLTYLSILR | P09661 | 533.32 | 2 |
| RUVB1 | GTEDITSPHGIPLDLLDR | Q9Y265 | 650.34 | 3 |
| RUVB1 | ALESSIAPIVIFASNR | Q9Y265 | 844.47 | 2 |
| RUVB1 | AVLLAGPPGTGK | Q9Y265 | 540.82 | 2 |
| RUVB2 | IGLETSLR | Q9Y230 | 444.76 | 2 |
| RUVB2 | AAGVVLEMIR | Q9Y230 | 529.80 | 2 |
| RUVB2 | VYSLFLDESR | Q9Y230 | 614.81 | 2 |
| RUVB2 | GLGLDDALEPR | Q9Y230 | 578.30 | 2 |
| RUVB2 | ALESDMAPVLIMATNR | Q9Y230 | 866.44 | 2 |
| RUVB2 | AVLIAGQPGTGK | Q9Y230 | 556.33 | 2 |
| RUXE | VMVQPINLIFR | P62304 | 665.39 | 2 |
| RUXF | GVEEEEEDGEMRE | P62306 | 769.30 | 2 |
| RUXGL | GNSIIMLEALER | A8MWD9 | 673.36 | 2 |
| S10A4 | TDEAAFQK | P26447 | 455.22 | 2 |
| S10A6 | LMEDLDR | P06703 | 446.22 | 2 |
| S10A8 | ALNSIIDVYHK | P05109 | 424.90 | 3 |
| S10A9 | NIETIINTFHQYSVK | P06702 | 602.98 | 3 |
| S2540 | FGAVTVISPLELIR | Q8TBP6 | 757.95 | 2 |
| SAC1 | TNVIQSLLAR | Q9NTJ5 | 557.83 | 2 |
| SAE2 | ADPEAAWEPTEAEAR | Q9UBT2 | 821.87 | 2 |
| SAHH | VADIGLAAWGR | P23526 | 564.81 | 2 |
| SAMH1 | IIDTPQFQR | Q9Y3Z3 | 559.30 | 2 |
| SAMH1 | FENLGVSSLGER | Q9Y3Z3 | 654.33 | 2 |
| SAMP | VGEYSLYIGR | P02743 | 578.80 | 2 |
| SAMP | AYSLFSYNTQGR | P02743 | 703.84 | 2 |
| SAP | EIVDSYLPVILDIIK | P07602 | 865.50 | 2 |
| SARNP | FGLNVSSISR | P82979 | 540.30 | 2 |
| SARNP | FGIVTSSAGTGTTEDTEAK | P82979 | 936.45 | 2 |
| SARNP | [1Ac]-ATETVELHK | P82979 | 535.28 | 2 |
| SART3 | IQLIFER | Q15020 | 459.77 | 2 |
| SC11A | VGEIVVFR | P67812 | 459.77 | 2 |
| SC11A | GDLLFLTNR | P67812 | 524.79 | 2 |
| SC11A | MLSLDFLDDVR | P67812 | 662.33 | 2 |
| SC22B | IMVANIEEVLQR | O75396 | 707.89 | 2 |
| SC22B | NLGSINTELQDVQR | O75396 | 793.91 | 2 |
| SC22B | GEALSALDSK | O75396 | 495.76 | 2 |
| SC22B | VADGLPLAASMQEDEQSGR | O75396 | 987.47 | 2 |
| SC23A | MVVPVAALFTPLK | Q15436 | 693.41 | 2 |
| SC23B | AESEEGPDVLR | Q15437 | 601.29 | 2 |
| SC24C | AVITSLLDQIPEMFADTR | P53992 | 674.02 | 3 |
| SC24C | SLLDFLPR | P53992 | 480.78 | 2 |
| SC31A | EQTLSPTITSGLHNIAR | O94979 | 613.33 | 3 |
| SC31A | MADAIILAIAGGQELLAR | O94979 | 609.34 | 3 |
| SCAM3 | TAAANAAAGAAENAFR | O14828 | 738.86 | 2 |
| SCFD1 | [1Ac]-AAAAAATAAAAASIR | Q8WVM8 | 650.35 | 2 |
| SCMC1 | DYFLFNPVTDIEEIIR | Q6NUK1 | 992.51 | 2 |
| SCOT1 | MVSSYVGENAEFER | P55809 | 809.36 | 2 |
| SCRB2 | VEEVGPYTYR | Q14108 | 606.80 | 2 |
| SCRN1 | AIIESDQEQGR | Q12765 | 623.31 | 2 |
| SDHA | LGANSLLDLVVFGR | P31040 | 737.42 | 2 |
| SDHA | GEGGILINSQGER | P31040 | 665.34 | 2 |
| SEPT1 | LTQTLAIER | Q8WYJ6 | 522.81 | 2 |
| SEPT1 | GLRPLDVAFLR | Q8WYJ6 | 419.59 | 3 |
| SEPT2 | TIISYIDEQFER | Q15019 | 757.38 | 2 |
| SEPT2 | TVQIEASTVEIEER | Q15019 | 802.41 | 2 |
| SEPT7 | ILEQQNSSR | Q16181 | 537.78 | 2 |
| SEPT9 | YLQEEVNINR | Q9UHD8 | 639.33 | 2 |
| SERA | DLPLLLFR | O43175 | 493.81 | 2 |

| | | | | |
|---|---|---|---|---|
| SERPH | LYGPSSVSFADDFVR | P50454 | 830.40 | 2 |
| SERPH | DTQSGSLLFIGR | P50454 | 647.34 | 2 |
| SERPH | LFYADHPFIFLVR | P50454 | 546.63 | 3 |
| SET | VEVTEFEDIK | Q01105 | 604.81 | 2 |
| SET | LNEQASEEILK | Q01105 | 637.34 | 2 |
| SET | IDFYFDENPYFENK | Q01105 | 920.91 | 2 |
| SF01 | AYIVQLQIEDLTR | Q15637 | 781.43 | 2 |
| SF3A2 | TGSGGVASSSESNR | Q15428 | 648.29 | 2 |
| SF3A3 | YMEVSGNLR | Q12874 | 534.76 | 2 |
| SF3A3 | [1Ac]-METILEQQR | Q12874 | 595.30 | 2 |
| SF3B1 | EVMLILIR | O75533 | 493.81 | 2 |
| SF3B2 | VGEPVALSEEER | Q13435 | 657.83 | 2 |
| SF3B3 | TVLDPVTGDLSDTR | Q15393 | 744.88 | 2 |
| SF3B3 | MQGQEAVLAMSSR | Q15393 | 704.34 | 2 |
| SF3B3 | IVILEYQPSK | Q15393 | 595.34 | 2 |
| SFPQ | LFVGNLPADITEDEFKR | P23246 | 655.34 | 3 |
| SFPQ | NLSPYVSNELLEEAFSQFGPIER | P23246 | 880.44 | 3 |
| SFPQ | FGQGGAGPVGGQGPR | P23246 | 671.34 | 2 |
| SFPQ | LFVGNLPADITEDEFK | P23246 | 904.46 | 2 |
| SFPQ | FAQHGTFEYEYSQR | P23246 | 588.27 | 3 |
| SFXN1 | NILLTNEQLESAR | Q9H9B4 | 750.90 | 2 |
| SGT1 | [1Ac]-AAAAAGTATSQR | Q9Y2Z0 | 559.28 | 2 |
| SH3L1 | GDYDAFFEAR | O75368 | 595.76 | 2 |
| SH3L1 | VYIASSSGSTAIK | O75368 | 642.35 | 2 |
| SH3L3 | VYSTSVTGSR | Q9H299 | 528.77 | 2 |
| SH3L3 | IQYQLVDISQDNALRDEMR | Q9H299 | 769.72 | 3 |
| SH3L3 | IQYQLVDISQDNALR | Q9H299 | 888.47 | 2 |
| SHIP1 | FTHLFWFGDLNYR | Q92835 | 572.62 | 3 |
| SMAP2 | LYEAYLPETFR | Q8WU79 | 701.36 | 2 |
| SMAP2 | RPQIDPAVEGFIR | Q8WU79 | 499.94 | 3 |
| SMC1A | TALFEEISR | Q14683 | 533.28 | 2 |
| SMC1A | AFVSMVYSEEGAEDR | Q14683 | 845.37 | 2 |
| SMCA5 | LLNILMQLR | O60264 | 557.34 | 2 |
| SMD1 | NREPVQLETLSIR | P62314 | 518.96 | 3 |
| SMD2 | GDSVIVVLR | P62316 | 479.29 | 2 |
| SMD3 | FLILPDMLK | P62318 | 545.32 | 2 |
| SMD3 | VAQLEQVYIR | P62318 | 609.85 | 2 |
| SMU1 | LMALLGQALK | Q2TAY7 | 529.33 | 2 |
| SNAA | LDQWLTTMLLR | P54920 | 695.38 | 2 |
| SND1 | LGTLSPAFSTR | Q7KZF4 | 575.32 | 2 |
| SND1 | VLPAQATEYAFAFIQVPQDDDAR | Q7KZF4 | 855.76 | 3 |
| SND1 | TDAVDSVVR | Q7KZF4 | 481.25 | 2 |
| SND1 | SEAVVEYVFSGSR | Q7KZF4 | 715.35 | 2 |
| SND1 | SSHYDELLAAEAR | Q7KZF4 | 487.90 | 3 |
| SND1 | LRPLYDIPYMFEAR | Q7KZF4 | 595.31 | 3 |
| SND1 | [1Ac]-ASSAQSGGSSGGPAVPTVQR | Q7KZF4 | 921.95 | 2 |
| SNG2 | FLTQPQVVAR | O43760 | 579.84 | 2 |
| SNG2 | AGGSFDLR | O43760 | 411.71 | 2 |
| SNG2 | DVLVGADSVR | O43760 | 515.78 | 2 |
| SNP23 | [1Ac]-MDNLSSEEIQQR | O00161 | 746.34 | 2 |
| SNX2 | AVNTQALSGAGILR | O60749 | 685.89 | 2 |
| SODM | AIWNVINWENVTER | P04179 | 872.44 | 2 |
| SODM | GDVTAQIALQPALK | P04179 | 712.91 | 2 |
| SODM | [CRM]-NVRPDYLK[1Ac]AIWNVINWENVTER | P04179 | 704.36 | 4 |
| SON | [1Ac]-ATNIEQIFR | P18583 | 567.30 | 2 |
| SORCN | LMVSMLDR | P30626 | 482.75 | 2 |
| SP16H | YEEEEEQSR | Q9Y5B9 | 599.75 | 2 |
| SPB6 | IAELLSPGSVDPLTR | P35237 | 784.44 | 2 |
| SPB9 | AQLLELPYAR | P50453 | 587.33 | 2 |
| SPB9 | AFQSLLTEVNK | P50453 | 625.34 | 2 |
| SPCS2 | [1Ac]-AAAAVQGGR | Q15005 | 421.73 | 2 |
| SPEE | AAFVLPEFAR | P19623 | 560.81 | 2 |
| SPEE | YQDILVFR | P19623 | 527.29 | 2 |
| SPTB2 | LTTLELLEVR | Q01082 | 593.86 | 2 |
| SPTB2 | VQAVVAVAR | Q01082 | 456.78 | 2 |
| SPTB2 | DLMLWMEDVIR | Q01082 | 710.85 | 2 |
| SPTB2 | LWEYLLELLR | Q01082 | 674.39 | 2 |
| SPTN1 | AALLELWELR | Q13813 | 607.35 | 2 |
| SPTN1 | ELPTAFDYVEFTR | Q13813 | 794.39 | 2 |
| SPTN1 | DLASVQALLR | Q13813 | 543.32 | 2 |
| SQRD | GYWGGPAFLR | Q9Y6N5 | 562.29 | 2 |
| SQRD | TAAAVAAQSGILDR | Q9Y6N5 | 672.37 | 2 |
| SQRD | IMYLSEAYFR | Q9Y6N5 | 646.82 | 2 |
| SRRM2 | TAAALAPASLTSAR | Q9UQ35 | 650.86 | 2 |

| | | | | |
|---|---|---|---|---|
| SRS10 | GFAYVQFEDVR | O75494 | 665.82 | 2 |
| SRS10 | YGPIVDVYVPLDFYTR | O75494 | 958.99 | 2 |
| SRSF1 | DGTGVVEFVR | Q07955 | 539.78 | 2 |
| SRSF1 | GGPPFAFVEFEDPR | Q07955 | 782.88 | 2 |
| SRSF1 | SHEGETAYIR | Q07955 | 581.78 | 2 |
| SRSF2 | VGDVYIPR | Q01130 | 459.76 | 2 |
| SRSF2 | DAEDAMDAMDGAVLDGR | Q01130 | 876.36 | 2 |
| SRSF2 | [1Ac]-SYGRPPPDVEGMTSLK | Q01130 | 888.44 | 2 |
| SRSF3 | AFGYYGPLR | P84103 | 522.27 | 2 |
| SRSF4 | GESENAGTNQETR | Q08170 | 696.80 | 2 |
| SRSF5 | GFGFVEFEDPR | Q13243 | 650.30 | 2 |
| SRSF6 | TNEGVIEFR | Q13247 | 532.77 | 2 |
| SRSF7 | VELSTGMPR | Q16629 | 495.26 | 2 |
| SRSF7 | VYVGNLGTGAGK | Q16629 | 568.31 | 2 |
| SRSF9 | IYVGNLPTDVR | Q13242 | 623.84 | 2 |
| SSBP | VGQDPVLR | Q04837 | 442.25 | 2 |
| SSRD | FFDEESYSLLR | P51571 | 703.34 | 2 |
| STAT1 | EGAITFTWVER | P42224 | 654.83 | 2 |
| STAT1 | FSLENNFLLQHNIR | P42224 | 582.31 | 3 |
| STAT1 | FHDLLSQLDDQYSR | P42224 | 579.61 | 3 |
| STAT1 | TFSLFQQLIQSSFVVER | P42224 | 677.03 | 3 |
| STAT1 | [1Ac]-SQWYELQQLDSK | P42224 | 783.88 | 2 |
| STAT2 | LTTLIELLLPK | P52630 | 627.41 | 2 |
| STAT3 | SIVSELAGLLSAMEYVQK | P40763 | 646.68 | 3 |
| STIP1 | LAYINPDLALEEK | P31948 | 744.90 | 2 |
| STIP1 | LMDVGLIAIR | P31948 | 550.83 | 2 |
| STML2 | ILEPGLNILIPVLDR | Q9UJZ1 | 838.01 | 2 |
| STML2 | ATVLESEGTR | Q9UJZ1 | 531.77 | 2 |
| STMN1 | ASGQAFELILSPR | P16949 | 694.88 | 2 |
| STOM | AMAAEAEASR | P27105 | 503.73 | 2 |
| STOM | VQNATLAVANITNADSATR | P27105 | 965.50 | 2 |
| STOM | VIAAEGEMNASR | P27105 | 624.31 | 2 |
| STT3A | VGQAMASTEEK | P46977 | 575.77 | 2 |
| STT3B | FGEMQLDFR | Q8TCJ2 | 571.77 | 2 |
| STT3B | ESDYFTPQGEFR | Q8TCJ2 | 738.33 | 2 |
| STX7 | TLNQLGTPQDSPELR | O15400 | 834.93 | 2 |
| SUCB2 | ETYLAILMDR | Q96I99 | 612.82 | 2 |
| SUN2 | LTTAASLLDVFVLTR | Q9UH99 | 810.47 | 2 |
| SURF4 | NLALGGGLLLLLAESR | O15260 | 805.49 | 2 |
| SWP70 | LQTQVELQAR | Q9UH65 | 593.33 | 2 |
| SYAC | AVFDETYPDPVR | P49588 | 704.84 | 2 |
| SYAC | IVAVTGAEAQK | P49588 | 543.81 | 2 |
| SYAC | [1Ac]-MDSTLTASEIR | P49588 | 633.31 | 2 |
| SYDC | FGAPPHAGGGIGLER | P14868 | 479.25 | 3 |
| SYDC | LPLQLDDAVRPEAEGEEEGR | P14868 | 741.70 | 3 |
| SYDC | ESIVDVEGVVR | P14868 | 601.32 | 2 |
| SYDM | IIDISDVFR | Q6PI48 | 539.30 | 2 |
| SYFA | VVDSMEDEVQR | Q9Y285 | 653.80 | 2 |
| SYFB | DLLFQALGR | Q9NSD9 | 516.80 | 2 |
| SYG | TFFSFPAVVAPFK | P41250 | 729.40 | 2 |
| SYIC | FLIQNVLR | P41252 | 501.81 | 2 |
| SYIC | LYLINSPVVR | P41252 | 587.35 | 2 |
| SYIC | LLILMEAR | P41252 | 479.79 | 2 |
| SYK | YLDLILNDFVR | Q15046 | 690.88 | 2 |
| SYK | MLVVGGIDR | Q15046 | 480.27 | 2 |
| SYLC | FDDPLLGPR | Q9P2J5 | 515.27 | 2 |
| SYLC | VDIGDTIIYLVH | Q9P2J5 | 679.37 | 2 |
| SYNC | FLTWILNR | O43776 | 531.81 | 2 |
| SYNC | NLMFLVLR | O43776 | 503.30 | 2 |
| SYNC | IFDSEEILAGYK | O43776 | 692.85 | 2 |
| SYPL1 | TVTATFGYPFR | Q16563 | 630.32 | 2 |
| SYQ | LFTLTALR | P47897 | 467.79 | 2 |
| SYQ | DRPMEESLLLFEAMR | P47897 | 612.97 | 3 |
| SYRC | LFEFAGYDVLR | P54136 | 665.35 | 2 |
| SYVC | ALSPLEEWLR | P26640 | 607.33 | 2 |
| SYVC | SSAQDPQAVLGALGR | P26640 | 735.39 | 2 |
| SYVC | LSAAVTEAFVR | P26640 | 582.32 | 2 |
| SYWC | ALIEVLQPLIAEHQAR | P23381 | 601.02 | 3 |
| SYWC | ISFPAIQAAPSFSNSFPQIFR | P23381 | 775.74 | 3 |
| SYWC | [+1K]-SEP[Oxi]AS[Pho]LLELFNSIATQGELVR | P23381 | 833.42 | 3 |
| SYYC | TVVSGLVQFVPK | P54577 | 637.38 | 2 |
| SYYC | APWELLELR | P54577 | 563.82 | 2 |
| TADBP | TSDLIVLGLPWK | Q13148 | 671.39 | 2 |
| TADBP | FTEYETQVK | Q13148 | 572.78 | 2 |

| | | | | |
|---|---|---|---|---|
| TAGL | TLMALGSLAVTK | Q01995 | 602.85 | 2 |
| TAGL | LGFQVWLK | Q01995 | 495.79 | 2 |
| TAGL | EFTESQLQEGK | Q01995 | 648.31 | 2 |
| TAGL2 | TLMNLGGLAVAR | P37802 | 608.35 | 2 |
| TAGL2 | NVIGLQMGTNR | P37802 | 601.82 | 2 |
| TAGL2 | DDGLFSGDPNWFPK | P37802 | 797.86 | 2 |
| TAGL2 | ENFQNWLK | P37802 | 539.77 | 2 |
| TALDO | LFVLFGAEILK | P37837 | 625.38 | 2 |
| TALDO | [CRM]-NAIDK[1Ac]LFVLFGAEILK | P37837 | 626.02 | 3 |
| TAP1 | IFSLLVPTALPLLR | Q03518 | 776.99 | 2 |
| TAP1 | QVAAVGQEPQVFGR | Q03518 | 743.39 | 2 |
| TAP1 | ELISWGAPGSADSTR | Q03518 | 773.88 | 2 |
| TAP2 | EAVGGLQTVR | Q03519 | 515.29 | 2 |
| TAP2 | EQLFSSLLR | Q03519 | 546.81 | 2 |
| TBA1A | LIGQIVSSITASLR | Q71U36 | 729.44 | 2 |
| TBA1A | [1Ac]-LIGQIVSSITASLR | Q71U36 | 500.63 | 3 |
| TBA4A | EIIDPVLDR | P68366 | 535.30 | 2 |
| TBA4A | AVFVDLEPTVIDEIR | P68366 | 858.46 | 2 |
| TBB2A | YLTVAAIFR | Q13885 | 527.31 | 2 |
| TBB4B | AVLVDLEPGTMDSVR | P68371 | 801.41 | 2 |
| TBB4B | INVYYNEATGGK | P68371 | 664.83 | 2 |
| TBB5 | MAVTFIGNSTAIQELFK | P07437 | 935.49 | 2 |
| TBB5 | ISVYYNEATGGK | P07437 | 651.32 | 2 |
| TBB5 | FWEVISDEHGIDPTGTYHGDSDLQLDR | P07437 | 776.36 | 4 |
| TBB5 | [AAR]-GLK[1Ac]M[Oxi]AVTFIGNSTAIQELFK | P07437 | 752.08 | 3 |
| TBCA | RLEAAYLDLQR | O75347 | 449.92 | 3 |
| TBCA | LEAAYLDLQR | O75347 | 596.32 | 2 |
| TBCB | AQQEAEAAQR | Q99426 | 551.27 | 2 |
| TCEA1 | EESTSSGNVSNR | P23193 | 633.78 | 2 |
| TCPA | EQLAIAEFAR | P17987 | 574.31 | 2 |
| TCPA | FATEAAITILR | P17987 | 603.35 | 2 |
| TCPA | YINENLIVNTDELGR | P17987 | 881.95 | 2 |
| TCPA | AFHNEAQVNPER | P17987 | 471.23 | 3 |
| TCPA | SQNVMAAASIANIVK | P17987 | 758.91 | 2 |
| TCPA | [1Ac]-MEGPLSVFGDR | P17987 | 625.30 | 2 |
| TCPB | GATQQILDEAER | P78371 | 665.83 | 2 |
| TCPB | LAVEAVLR | P78371 | 435.77 | 2 |
| TCPB | MLPTIIADNAGYDSADLVAQLR | P78371 | 783.07 | 3 |
| TCPB | LTSFIGAIAIGDLVK | P78371 | 759.45 | 2 |
| TCPB | [1Ac]-ASLSLAPVNIFK | P78371 | 651.38 | 2 |
| TCPB | [1Ac]-ASLSLAPVNIFKAGADEERA | P78371 | 701.04 | 3 |
| TCPD | ALIAGGGAPEIELALR | P50991 | 775.95 | 2 |
| TCPD | DALSDLALHFLNK | P50991 | 486.26 | 3 |
| TCPD | VIDPATATSVDLR | P50991 | 679.37 | 2 |
| TCPD | AYILNLVK | P50991 | 467.29 | 2 |
| TCPD | ETLLNSATTSLNSK | P50991 | 739.89 | 2 |
| TCPE | WVGGPEIELIAIATGGR | P48643 | 869.98 | 2 |
| TCPE | IADGYEQAAR | P48643 | 547.27 | 2 |
| TCPE | QMAEIAVNAVLTVADMER | P48643 | 654.33 | 3 |
| TCPE | [1Ac]-ASMGTLAFDEYGRPFLIIK | P48643 | 724.38 | 3 |
| TCPG | AVAQALEVIPR | P49368 | 583.85 | 2 |
| TCPG | TAVETAVLLLR | P49368 | 593.36 | 2 |
| TCPG | IVLLDSSLEYK | P49368 | 640.36 | 2 |
| TCPH | LPIGDVATQYFADR | Q99832 | 783.40 | 2 |
| TCPH | GGAEQFMEETER | Q99832 | 692.30 | 2 |
| TCPH | TATQLAVNK | Q99832 | 473.27 | 2 |
| TCPH | [1Ac]-MMPTPVILLK | Q99832 | 592.84 | 2 |
| TCPQ | DIDEVSSLLR | P50990 | 573.80 | 2 |
| TCPQ | LATNAAVTVLR | P50990 | 564.84 | 2 |
| TCPQ | LVPGGGATEIELAK | P50990 | 677.88 | 2 |
| TCPQ | LFVTNDAATILR | P50990 | 667.38 | 2 |
| TCPQ | FAEAFEAIPR | P50990 | 575.80 | 2 |
| TCPQ | GSTDNLMDDIER | P50990 | 683.30 | 2 |
| TCPZ | IITEGFEAAK | P40227 | 539.79 | 2 |
| TCPZ | AQLGVQAFADALLIIPK | P40227 | 884.52 | 2 |
| TCTP | DLISHDEMFSDIYK | P13693 | 571.60 | 3 |
| TENA | APTAQVESFR | P24821 | 553.29 | 2 |
| TENA | LEELENLVSSLR | P24821 | 701.38 | 2 |
| TENA | VATYLPAPEGLK | P24821 | 629.86 | 2 |
| TENA | ITAQGQYELR | P24821 | 589.81 | 2 |
| TERA | IVSQLLTLMDGLK | P55072 | 715.92 | 2 |
| TERA | EVDIGIPDATGR | P55072 | 621.82 | 2 |
| TERA | WALSQSNPSALR | P55072 | 665.35 | 2 |
| TERA | EMVELPLR | P55072 | 493.77 | 2 |

| | | | | |
|---|---|---|---|---|
| TERA | GILLYGPPGTGK | P55072 | 586.84 | 2 |
| TERA | LDQLIYIPLPDEK | P55072 | 778.93 | 2 |
| TFR1 | SSGLPNIPVQTISR | P02786 | 734.91 | 2 |
| TFR1 | ILNIFGVIK | P02786 | 508.83 | 2 |
| TFR1 | YNSQLLSFVR | P02786 | 613.83 | 2 |
| TGM2 | ALLVEPVINSYLLAER | P21980 | 900.52 | 2 |
| TGM2 | [1Ac]-AEELVLER | P21980 | 500.77 | 2 |
| THIC | ILVTLLHTLER | Q9BWD1 | 436.61 | 3 |
| THIK | AEELGLPILGVLR | P09110 | 690.42 | 2 |
| THIL | EVVIVSATR | P24752 | 487.29 | 2 |
| THIL | NEQDAYAINSYTR | P24752 | 772.85 | 2 |
| THIO | TAFQEALDAAGDK | P10599 | 668.82 | 2 |
| THMS2 | [1Ac]-MEPVPLQDFVR | Q5TEJ8 | 686.85 | 2 |
| TIF1B | ADVQSIIGLQR | Q13263 | 600.34 | 2 |
| TIM50 | VLLDLSAFLK | Q3ZCQ8 | 559.84 | 2 |
| TINAL | LDGAWWFLR | Q9GZM7 | 582.30 | 2 |
| TIPRL | VMPSSFFLLLR | O75663 | 655.37 | 2 |
| TKT | VLDPFTIKPLDR | P29401 | 471.94 | 3 |
| TKT | ILTVEDHYYEGGIGEAVSSAVVGEPGITVTHLAVNR | P29401 | 938.98 | 4 |
| TKT | LDNLVAILDINR | P29401 | 684.90 | 2 |
| TKT | DAIAQAVR | P29401 | 422.24 | 2 |
| TKT | NMAEQIIQEIYSQIQSK | P29401 | 675.01 | 3 |
| TLN1 | GVGAAATAVTQALNELLQHVK | Q9Y490 | 697.72 | 3 |
| TLN1 | VAGSVTELIQAAEAMK | Q9Y490 | 809.43 | 2 |
| TLN1 | TLAESALQLLYTAK | Q9Y490 | 761.43 | 2 |
| TLN1 | VMVTNVTSLLK | Q9Y490 | 602.85 | 2 |
| TLN1 | TMLESAGGLIQTAR | Q9Y490 | 724.38 | 2 |
| TM109 | EAPVDVLTQIGR | Q9BVC6 | 649.36 | 2 |
| TM109 | ASGAQLEAK | Q9BVC6 | 437.74 | 2 |
| TMED9 | FSLFAGGMLR | Q9BVK6 | 549.79 | 2 |
| TMM33 | ALLANALTSALR | P57088 | 607.37 | 2 |
| TMX1 | VDVTEQPGLSGR | Q9H3N1 | 629.33 | 2 |
| TNAP2 | VLLVELPAFLR | Q03169 | 635.40 | 2 |
| TNPO1 | FSDQFPLPLK | Q92973 | 596.32 | 2 |
| TNPO1 | ALVMLLEVR | Q92973 | 522.32 | 2 |
| TNPO1 | ATVGILITTIASK | Q92973 | 644.40 | 2 |
| TOIP1 | SQPAILLLTAAR | Q5JTV8 | 627.38 | 2 |
| TOM22 | LWGLTEMFPER | Q9NS69 | 689.84 | 2 |
| TOM40 | LPPLPLTLALGAFLNHR | O96008 | 615.04 | 3 |
| TOP2B | NTVEITELPVR | Q02880 | 635.85 | 2 |
| TPD54 | TSAALSTVGSAISR | O43399 | 660.86 | 2 |
| TPIS | VVLAYEPVWAIGTGK | P60174 | 801.95 | 2 |
| TPIS | SNVSDAVAQSTR | P60174 | 617.80 | 2 |
| TPM3 | AADAEAEVASLNR | P06753 | 658.83 | 2 |
| TPM3 | KLVIIEGDLER | P06753 | 428.92 | 3 |
| TPM4 | IQALQQQADEAEDR | P67936 | 807.89 | 2 |
| TPM4 | KIQALQQQADEAEDR | P67936 | 581.63 | 3 |
| TPM4 | [1Ac]-AGLNSLEAVK | P67936 | 522.29 | 2 |
| TPP1 | ILSGRPPLGFLNPR | O14773 | 512.97 | 3 |
| TPR | VLLMELEEAR | P12270 | 601.83 | 2 |
| TPR | ASTALSNEQQAR | P12270 | 638.32 | 2 |
| TPR | [1Ac]-AAVLQQVLER | P12270 | 584.84 | 2 |
| TPSN | VSLMPATLAR | O15533 | 529.80 | 2 |
| TRA2A | GFAFVYFER | Q13595 | 568.28 | 2 |
| TRA2A | YGPLSGVNVVYDQR | Q13595 | 783.90 | 2 |
| TRA2B | RPHTPT[Pho]PGIYMGR | P62995 | 521.58 | 3 |
| TRA2B | RPHT[Pho]PTPGIYMGR | P62995 | 521.58 | 3 |
| TRAP1 | AQLLQPTLEINPR | Q12931 | 746.93 | 2 |
| TRFE | MYLGYEYVTAIR | P02787 | 739.87 | 2 |
| TRFE | TAGWNIPMGLLYNK | P02787 | 789.41 | 2 |
| TRFE | APNHAVVTR | P02787 | 482.77 | 2 |
| TSN | EAVTEILGIEPDR | Q15631 | 721.38 | 2 |
| TSN | [1Ac]-SVSEIFVELQGFLAAEQDIR | Q15631 | 765.06 | 3 |
| TSN | [1Ac]-SVSEIFVELQ[Dea]GFLAAEQDIR | Q15631 | 765.39 | 3 |
| TSNAX | VTPVDYLLGVADLTGELMR | Q99598 | 688.03 | 3 |
| TTC38 | VLELLLPIR | Q5R3I4 | 533.36 | 2 |
| TXND5 | GYPTLLLFR | Q8NBS9 | 540.32 | 2 |
| TXND5 | GYPTLLWFR | Q8NBS9 | 576.81 | 2 |
| TXND5 | EYVESQLQR | Q8NBS9 | 576.29 | 2 |
| TXTP | FFVMTSLR | P53007 | 500.77 | 2 |
| TXTP | FGMFEFLSNHMR | P53007 | 505.90 | 3 |
| TXTP | GLSSLLYGSIPK | P53007 | 617.86 | 2 |
| TYPH | ALPLALVLHELGAGR | P19971 | 510.64 | 3 |
| TYPH | MLAAQGVDPGLAR | P19971 | 649.85 | 2 |

| | | | | |
|---|---|---|---|---|
| TYPH | ALQEALVLSDR | P19971 | 607.84 | 2 |
| TYPH | VAAALDDGSALGR | P19971 | 608.32 | 2 |
| U2AF1 | NPQNSSQSADGLR | Q01081 | 687.32 | 2 |
| U2AF1 | [1Ac]-AEYLASIFGTEK | Q01081 | 685.85 | 2 |
| U520 | SGGPVVVLVQLER | O75643 | 676.90 | 2 |
| U520 | WTELGALDILQMLGR | O75643 | 858.46 | 2 |
| U520 | EGSASTEVLR | O75643 | 524.77 | 2 |
| U520 | NALLQLTDSQIADVAR | O75643 | 864.47 | 2 |
| U5S1 | SFVEFILEPLYK | Q15029 | 742.91 | 2 |
| UB2L3 | ADLAEEYSK | P68036 | 513.24 | 2 |
| UB2V1 | WTGMIIGPPR | Q13404 | 564.31 | 2 |
| UBA1 | LQTSSVLVSGLR | P22314 | 630.37 | 2 |
| UBA1 | LAGTQPLEVLEAVQR | P22314 | 812.46 | 2 |
| UBA1 | SLVASLAEPDFVVTDFAK | P22314 | 955.00 | 2 |
| UBA1 | LDQPMTEIVSR | P22314 | 644.83 | 2 |
| UBA1 | ALPAVQQNNLDEDLIR | P22314 | 904.98 | 2 |
| UBC12 | GGYIGSTYFER | P61081 | 625.30 | 2 |
| UBE2K | GEIAGPPDTPYEGGR | P61086 | 758.36 | 2 |
| UBE2N | LLAEPVPGIK | P61088 | 518.82 | 2 |
| UBE2N | TNEAQAIETAR | P61088 | 602.30 | 2 |
| UBP14 | AQLFALTGVQPAR | P54578 | 686.39 | 2 |
| UBP14 | ASGEMASAQYITAALR | P54578 | 820.41 | 2 |
| UBP14 | RVEIMEEESEQ | P54578 | 689.81 | 2 |
| UBP5 | DGLGGLPDIVR | P45974 | 556.31 | 2 |
| UBXN1 | [1Ac]-AELTALESLIEMGFPR | Q04323 | 909.97 | 2 |
| UCHL5 | FNLMAIVSDR | Q9Y5K5 | 583.31 | 2 |
| UGGG1 | ILASPVELALVVMK | Q9NYU2 | 741.95 | 2 |
| URP2 | VFVGEEDPEAESVTLR | Q86UX7 | 888.94 | 2 |
| URP2 | VVLAGGVAPALFR | Q86UX7 | 635.39 | 2 |
| URP2 | ILEAHQNVAQLSLAEAQLR | Q86UX7 | 702.06 | 3 |
| URP2 | IDLAVGDVVK | Q86UX7 | 514.80 | 2 |
| USP9X | LAQQISDEASR | Q93008 | 609.31 | 2 |
| VA0D1 | LYPEGLAQLAR | P61421 | 615.85 | 2 |
| VAMP3 | ADALQAGASQFETSAAK | Q15836 | 833.41 | 2 |
| VAMP8 | [1Ac]-MEEASEGGGNDR | Q9BV40 | 647.25 | 2 |
| VASP | EEIIEAFVQELR | P50552 | 738.39 | 2 |
| VAT1 | LPPLPVTPGMEGAGVVIAVGEGVSDR | Q99536 | 839.78 | 3 |
| VATA | TALVANTSNMPVAAR | P38606 | 758.40 | 2 |
| VATA | GVNVSALSR | P38606 | 451.76 | 2 |
| VATA | LAEMPADSGYPAYLGAR | P38606 | 891.43 | 2 |
| VATA | FTMVQVWPVR | P38606 | 631.84 | 2 |
| VATB2 | IPQSTLSEFYPR | P21281 | 719.37 | 2 |
| VATB2 | TPVSEDMLGR | P21281 | 552.77 | 2 |
| VATB2 | TVFETLDIGWQLLR | P21281 | 845.96 | 2 |
| VATB2 | QIYPPINVLPSLSR | P21281 | 798.96 | 2 |
| VATE1 | ARDDLITDLLNEAK | P36543 | 529.62 | 3 |
| VATG1 | [1Ac]-ASQSQGIQQLLQAEK | O75348 | 835.94 | 2 |
| VDAC1 | VTQSNFAVGYK | P21796 | 607.31 | 2 |
| VDAC1 | WTEYGLTFTEK | P21796 | 687.83 | 2 |
| VIGLN | LVGEIMQETGTR | Q00341 | 667.34 | 2 |
| VIME | ILLAELEQLK | P08670 | 585.36 | 2 |
| VIME | SLYASSPGGVYATR | P08670 | 714.86 | 2 |
| VIME | ISLPLPNFSSLNLR | P08670 | 785.95 | 2 |
| VIME | EEAENTLQSFR | P08670 | 662.31 | 2 |
| VIME | DNLAEDIMR | P08670 | 538.76 | 2 |
| VIME | LGDLYEEEMR | P08670 | 627.79 | 2 |
| VINC | MALLMAEMSR | P18206 | 576.78 | 2 |
| VINC | ELTPQVVSAAR | P18206 | 585.83 | 2 |
| VINC | VDQLTAQLADLAAR | P18206 | 742.91 | 2 |
| VINC | AIPDLTAPVAAVQAAVSNLVR | P18206 | 692.73 | 3 |
| VKORL | GFGLLGSIFGK | Q8N0U8 | 548.31 | 2 |
| VP26A | LFLAGYDPTPTMR | O75436 | 741.38 | 2 |
| VPS35 | LSQLEGVNVER | Q96QK1 | 622.34 | 2 |
| VPS35 | NIIIALIDR | Q96QK1 | 520.83 | 2 |
| VPS4B | GILLFGPPGTGK | O75351 | 578.84 | 2 |
| VTNC | DVWGIEGPIDAAFTR | P04004 | 823.91 | 2 |
| VTNC | DWHGVPGQVDAAMAGR | P04004 | 556.26 | 3 |
| VTNC | FEDGVLDPDYPR | P04004 | 711.83 | 2 |
| WASF2 | DVVGNDVATILSR | Q9Y6W5 | 679.87 | 2 |
| WDFY4 | DSGLLGLLLAQLR | Q6ZS81 | 684.91 | 2 |
| WDR1 | IAVVGEGR | O75083 | 400.73 | 2 |
| XPO1 | ETLVYLTHLDYVDTER | O14980 | 656.33 | 3 |
| XPO1 | IYLDMLNVYK | O14980 | 636.34 | 2 |
| XPO1 | FLVTVIK | O14980 | 410.27 | 2 |

| | | | | |
|---|---|---|---|---|
| XPO1 | FLNVPMFR | O14980 | 512.28 | 2 |
| XPO2 | LVLDAFALPLTNLFK | P55060 | 838.00 | 2 |
| XPP1 | IENVVLVVPVK | Q9NQW7 | 604.88 | 2 |
| XRCC5 | HLMLPDFDLLEDIESK | P13010 | 638.99 | 3 |
| XRCC5 | HIEIFTDLSSR | P13010 | 659.34 | 2 |
| XRCC6 | DIISIAEDEDLR | P12956 | 694.85 | 2 |
| XRCC6 | DTGIFLDLMHLK | P12956 | 468.25 | 3 |
| XRCC6 | SDSFENPVLQQHFR | P12956 | 568.61 | 3 |
| XRCC6 | LGSLVDEFK | P12956 | 504.27 | 2 |
| XRCC6 | TFNTSTGGLLLPSDTK | P12956 | 826.43 | 2 |
| YBOX1 | GAEAANVTGPGGVPVQGSK | P67809 | 848.44 | 2 |
| YBOX1 | SVGDGETVEFDVVEGEK | P67809 | 898.42 | 2 |
| ZCCHV | FVVLETGGEAGITR | Q7Z2W4 | 724.89 | 2 |
